# Decision analysis in support of proactive planning for chronic wasting disease in Vermont, USA

**DOI:** 10.64898/2026.07.17.739197

**Authors:** Jonathan D. Cook, Annabelle Stanley, Brittany Mosher, Nicholas Fortin, Katherina Gieder, David Sausville, John Austin, Steve Agius, Tim Appleton, Jaclyn Comeau, Kristin Haas, Paul Hamelin, Melanie Kunkel, Natalie Kwit, Megan Cahill, Shawn Langston, Matt Leonard, Kaitlynn Levine, Katherine McNamara, Meredith Naughton, Fredrick Pogmore, Justin Stedman, Ken Sturm, Michael C. Runge

**Affiliations:** U.S. Geological Survey, Eastern Ecological Science Center, Laurel, MD; University of Vermont, Rubenstein School of the Environment and Natural Resources, Burlington, VT; Vermont Department of Fish and Wildlife, Agency of Natural Resources, Montpelier, VT; U.S. Fish and Wildlife Service, National Wildlife Refuge System, Brunswick, VT; Vermont Agency of Agriculture, Food and Markets, Montpelier, VT; Northeast Association of Fish and Wildlife Agencies, Ithaca, NY; Vermont Department of Health, Infectious Disease Epidemiology, Waterbury, VT; U.S. Forest Service, Forest Service Green Mountain & Finger Lakes National Forests, East Mendon, VT; Department of Forests, Parks and Recreation, Agency of Natural Resources, Essex, VT; United States Department of Agriculture Animal and Plant Health Inspection Service, Wildlife Services, VT District Office, Berlin, VT; U.S. Fish and Wildlife Service National Wildlife Refuge System, Swanton, VT

## Abstract

1

Chronic wasting disease (CWD), a fatal, transmissible disease in white-tailed deer (*Odocoileus virginianus*) and related species, is spreading across North America but has not yet been detected in Vermont, United States (U.S.). The Vermont Department of Fish and Wildlife, along with partner agencies, wants to develop a proactive prevention and response plan in anticipation of the eventual detection of the disease. Between September 2023 and September 2025, staff from the U.S. Geological Survey and the University of Vermont facilitated a structured decision-making (SDM) process with seven State and Federal agencies that have jurisdiction over some aspect of CWD management in Vermont. The aim of this process was to generate and evaluate alternative response plans against a range of long-term objectives important to the agencies. To aid in the evaluation of the alternatives, we developed a linked set of models for white-tailed deer population and disease dynamics, hunter participation and health, forest health, economic consequences, and agricultural opportunities related to the actions being contemplated as part of the response plan. We evaluated over 256 different permutations of management actions and used multi-criteria decision analysis, a branch of decision analysis designed to help decision makers navigate tradeoffs among competing objectives, to summarize the performance of those alternative strategies against the desired outcomes. Proactive actions—those designed to slow the arrival of CWD to Vermont—were moderately effective, but the most important proactive action was surveillance to detect the disease early after arrival, which triggered response actions after detection. With the insights generated by the SDM process and the results of the analyses, the participating agencies were able to identify a preferred strategy and outline the elements of a proactive response plan. This report describes the SDM process, the technical details of the modeling work, and the results of the analyses. It is intended to serve as the technical basis for Vermont’s response plan.

**Executive Summary:** Chronic wasting disease (CWD) is a fatal disease of deer, moose, and other cervids that is spreading across the United States (U.S.). As of December 2025, the nearest infected wild deer population to Vermont was in Pennsylvania, about 225 miles away; however, the disease has been detected closer to the state in New York and Quebec, Canada, but those detections are assumed to have been isolated and removed without additional spread in wild populations or captive facilities. The primary pathways of arrival to Vermont are natural spread in wild deer, import of infected live animals to captive cervid facilities, or disposal of imported infected deer parts on the Vermont landscape. Once CWD arrives, significant changes to many ecological, social, and economic resources are expected.

Many State and Federal agencies could have a role in managing CWD, before and after its arrival.

- Vermont Department of Fish and Wildlife (VDFW) has statutory responsibility for managing the wild deer and moose populations in the State, including through hunting;
- Vermont Agency of Agriculture, Food and Markets (VAAFM) is responsible for oversight of the nine captive cervid facilities in the State;
- Vermont Department of Forests, Parks and Recreation (VFPR) manages State Forests and Parks, which provide habitat for deer and recreational access for hunters;
- Vermont Department of Health (VDH) provides information and guidance to the public that enables them to make informed decisions that can affect their health, including whether and how CWD poses any risk to humans;
- U.S. Fish and Wildlife Service (USFWS) manages two National Wildlife Refuges in the State, which provide public access, including to hunters;
- U.S. Department of Agriculture Wildlife Services can provide contractual support for wildlife management, including targeted reduction of deer populations; and
- U.S. Forest Service manages extensive forest resources in the State, which provide habitat for deer and access for hunters.

These agencies have expressed interest in co-development of a long-term collaborative plan for management of CWD. From September 2023 through September 2025, they engaged in a series of meetings, facilitated using the principles of structured decision making to explore potential proactive and reactive management strategies, and how well those strategies might perform against the fundamental objectives being sought by the agencies. As part of this work, the U.S. Geological Survey and the University of Vermont developed quantitative predictive models to allow evaluation of alternative strategies against the fundamental objectives.

The agencies identified seven fundamental objectives for a long-term (20-year) CWD management plan that arise from their enabling legislation and missions:

- Maximize herd health of native cervid populations in the state of Vermont;
- Maximize the public benefit of cervids (consumptive and non-consumptive);
- Minimize harm to other species, unique ecosystems, and forests;
- Minimize costs, staff time, and workload associated with CWD management;
- Minimize the effect (or potential effect) that CWD has on public health;
- Minimize the effect of CWD and its management on local economies; and
- Maintain opportunities for private landowners to engage in agricultural pursuits.

In initial work to explore the effects of actions on the objectives, six broadly different alternative strategies were developed, where the strategies differed in philosophy, investment, timing, and proactive versus reactive emphasis.

- Under the *Status Quo* strategy, existing protocols would remain in place until CWD is detected in the State, at which point an increase in surveillance and harvest opportunity would be implemented.
- The *Heavy Outreach* strategy would rely primarily on outreach and communication with the public, notably hunters, to delay arrival, slow transmission, and mitigate impacts.
- The *Reduce Arrival* strategy would invest primarily in proactive measures including both outreach and regulatory changes to slow arrival of CWD to the State.
- The *Prepare and React* strategy would invest primarily in reactive measures after CWD arrives, with preparatory measures to detect it as early as possible and have in place the management tools to deploy upon arrival.
- The *Prevent and React* strategy would make a moderate investment in both proactive and reactive measures.
- Under the *Kitchen Sink* strategy, extensive investment would be made in both proactive and reactive measures.

The estimated mean time to arrival of CWD in the State is 7.7 years under the *Status Quo* strategy. Concerted efforts to delay arrival could increase this time to 10.6 years, by significantly reducing two of the three arrival pathways. It will, however, take an average of 15−30 years after arrival to detect CWD in the State assuming effort is spread evenly throughout the State, owing to its slow growth in the deer population. Investment in surveillance leads to earlier detection and hence, an earlier opportunity to implement reactive measures.

There are notable trade-offs among the alternative strategies in their ability to achieve outcomes of interest to the State, based on the analysis described in this report.

- The arrival and spread of CWD can be slowed quite significantly, particularly with a combination of proactive and reactive measures, with the *Kitchen Sink* showing a 6-fold reduction in prevalence over 50 years compared to the *Status Quo* strategy.
- The number of CWD-positive deer consumed by hunters, their families, and their friends largely tracks the prevalence of CWD in the population, although increased outreach efforts also contribute to reduced consumption.
- Deer harvest, and the economic benefits associated with it, is expected to increase under aggressive CWD management strategies, because the strategies encourage higher harvest to manage the disease, and lower prevalence of CWD keeps more hunters participating.
- The positive outcomes associated with more aggressive proactive and reactive measures are counterbalanced by increases in costs to agencies, for surveillance, sharpshooting, outreach, and captive cervid management.

Captive cervid farm operations in Vermont were estimated to be a low-risk introduction pathway because of the stringent regulations, agency oversight of importation, and disease testing that is already occurring. However, actions by VDFW could reduce the risk of spread from wild to captive deer. The more aggressive CWD management strategies reduce CWD prevalence and increase the long-term opportunity for captive cervid operations. The one exception to this is the *Kitchen Sink* strategy, which seeks to reduce the risk of arrival of CWD by proactively closing captive cervid facilities.

The size of the statewide deer population is expected to be about the same in 20 years across all strategies, with all alternatives resulting in a decrease of about 5% compared to current population size estimates. In some strategies, the deer population decreases because of the effects of CWD; in others, the deer population decreases because of increased hunting that slows the transmission of CWD.

After considering the results from the analysis of the six initial strategies, the participating agencies were interested in more carefully exploring the tradeoffs among three of the objectives: long-term prevalence of CWD in the State; the cost of management of CWD; and the opportunity for private farms to participate in captive cervid farming. An additional 256 strategies, which were permutations of the constituent actions under consideration, were created and analyzed. These results showed that the most effective way to keep CWD prevalence low over the long term was extensive surveillance to allow early detection, followed by sharpshooting or other targeted removal at a local level, to continually eliminate or reduce new incursions of the disease. This approach, however, is quite costly, so the balance between effort and effect was important to the agencies. Regarding captive cervid operations, proactive closing of captive cervid facilities does slightly lower the long-term prevalence of CWD and substantially diminishes the long-term costs of closing infected captive cervid farms in the future, but comes at the cost of farming opportunity.

In September 2025, the participating agencies expressed a collective interest in a strategy that preserves captive cervid farming opportunities, invests heavily in surveillance for early detection of the disease, and prepares for targeted removal of deer when detections occur. Such a strategy, while potentially costly, achieves many of the objectives sought by the participating agencies on behalf of the people of Vermont. The agencies, without yet making any commitments, intend to undertake work to develop the details of this plan, discuss its aspects with stakeholders, and build support for its long-term implementation.

## 3 Introduction

Chronic wasting disease (CWD) is a transmissible spongiform encephalopathy that affects cervids (Williams and Young, 1980), including white-tailed deer (*Odocoileus virginianus*) and moose (*Alces alces*). The disease is caused by a misfolded prion protein (PrP^CWD^) that accumulates in neural tissue causing cell death, declining neural function, and eventual host death (Prusiner, 1982). Chronic wasting disease is transmitted via direct contact with an infectious individual or indirectly through contact with a contaminated environment (Mathiason and others, 2006). Misfolded prions are shed in bodily fluids and can persist in the environment for years (Johnson and others, 2006). The disease spreads across the landscape and increases in prevalence slowly (Wasserberg and others, 2009). While it has not yet been detected in Vermont, it occurs in 37 other U.S. states, five Canadian provinces, and is detected moving closer to the Stateeach year. The closest known location of CWD in wild populations is found in Pennsylvania (Richards, 2021). The disease affects both wild and captive cervids.

Once introduced into a new area, CWD is extremely challenging to eradicate because of its prolonged period of infectivity in hosts, stability in the environment, and lack of visually apparent symptoms that would allow for targeted removal. Because the disease is always fatal with no individual immunity, CWD can cause population declines in areas where prevalence is high (Edmunds and others, 2016; DeVivo and others, 2017; Gaya and others, 2026). Population declines of widespread and popular game species can affect hunting, viewing and farming opportunities, state conservation budgets, and the composition and configuration of forested habitats. Finally, there is a potential for hunters to consume CWD-affected meat, as nearly the entire hunting public in Vermont pursues white-tailed deer (96%, Vermont Fish and Wildlife Department, 2021b). While there has not been documented transmission of CWD from deer to humans, the U.S. Centers for Disease Control and Prevention (CDC) recommend testing game in areas where CWD is present and avoiding consumption of CWD-positive meat (https://www.cdc.gov/chronic-wasting/about/index.html).

The magnitude of future CWD impacts on multiple Vermont sectors is unknown, as is the efficacy of actions that could mitigate disease-associated concerns. Proactive actions, early detection, and an aggressive response could improve outcomes, but it is unclear how much success is possible and whether the associated costs can be met. To explore these uncertainties and identify the proactive, reactive, and surveillance elements of a CWD management plan for the state of Vermont, the Vermont Department of Fish and Wildlife (VDFW), U.S. Fish and Wildlife Service (USFWS), U.S. Forest Service (USFS), U.S. Department of Agriculture (USDA), Vermont Department of Health (VDH), Vermont Agency of Agriculture, Food and Markets (VAAFM), Vermont Department of Forests, Parks and Recreation (VFPR), U.S. Geological Survey (USGS), and the University of Vermont (UVM) have been working together to frame and evaluate a range of CWD management strategies in anticipation of the disease’s eventual detection in the state. While many of the State and Federal agencies directly contributed to discerning the viability of management actions, USGS and UVM contributed to facilitation and technical tasks solely. The purpose of this report is to describe the technical work that can be used to support the multi-agency plan, including the use of structured decision-making (SDM), mathematical modeling, and agency deliberation that could ultimately lead to a management strategy for CWD in Vermont deer and moose.

### 3.1 Background

Vermont has a rich history of managing white-tailed deer, moose, and wildlife diseases using the best available science and in consideration of its stewardship mission (Vermont Fish and Wildlife Department, 2021b). The agencies are guided by a broad set of legal and regulatory authorities that cover free-ranging and captive cervids, agricultural production, land management, and public health. The management of big game is described in the Vermont Big Game Management Plan 2020–2030, including the natural history of deer and moose and specific goals for their management (Vermont Fish and Wildlife Department, 2021b). There are other documents, such as the VFPR Division of Forests Strategic Plan 2025–2028 (VFPR, 2025), VDH Strategic Plan 2024−2029 (VDH, 2024), and Vermont Agriculture and Food System Strategic Plan 2021–2030 (Claro and others, 2021) that provide additional context in support of this work. We briefly draw on these plans to offer background and then describe the specific legal, regulatory, and political contexts of the agencies involved in this planning effort.

#### 3.1.1 Ecological Context

White-tailed deer are broadly distributed and abundant throughout Vermont. They occur in all 14 counties of the state and population objectives range from <10 to 18 deer per square mile (Vermont Fish and Wildlife Department, 2021b). While deer are highly regarded, the abundance of white-tailed deer exceeds desirable levels in some locations and has led to habitat damage and increasing wildlife conflict, including damage to gardens and agricultural crops and increasing rates of human vector-borne diseases (Vermont Fish and Wildlife Department, 2021b). In terms of harvest and deer management, Vermont hunters take an average of about 17,000 deer during the fall hunting seasons, providing over 3 million venison meals per year (Vermont Fish and Wildlife Department, 2020; 2021a; 2022; 2023; 2024). Hunting license sales have declined 2−3% per year, on average, over the last two decades and this may affect the ability to manage deer in the future. In terms of captive deer, farming opportunities are valued, but relative to other states (e.g., Pennsylvania and New York) are rare; there are a total of nine captive deer facilities that house red deer (*Cervus elaphus*), fallow deer (*Dama dama*), reindeer (*Rangifer tarandus*), and elk (*Cervus canadensis*).

Compared to deer, moose are much less abundant and do not occur statewide. They are primarily restricted to the central and northeastern parts of Vermont where their densities fall between 0.5 and 1.0 moose per square mile (Vermont Fish and Wildlife Department, 2021b). The current management objective for moose is targeted to keep moose density below 1.0 per square mile to avoid damage to forest regeneration and excessive mortality caused by winter ticks. In terms of CWD, moose are susceptible to the disease but there is little empirical data on disease dynamics with overlapping deer and moose populations. Nonetheless, some evidence exists that moose CWD cases are less common and less likely to transition to epidemic outbreaks (Mysterud, 2023). As a result, the primary focus of this technical work is on white-tailed deer with any targeted actions for CWD also expected to benefit moose.

Chronic wasting disease is broadly distributed throughout North America and is expanding in spatial distribution through at least three mechanisms: natural disease dispersal in wild populations (through animal movement and disease spread); human transport of infected captive animals to new locations; and human-mediated movement of infected cervid parts (carcasses, skins, and fluids from harvested animals) (Schuler and others, 2016; Cook and others, 2022).

There has been extensive study of white-tailed deer spatial dynamics and movement, including in conjunction with CWD modeling, but there is not yet a predictive model that provides the expected time-to-arrival to northeastern states. As a result, it is unclear how long until, and by which mechanism, CWD is likely to arrive in Vermont. In other states where CWD occurs, the interaction between wild and captive cervids has affected disease spread dynamics. Captive cervid facilities, even those that are USDA-certified, can be a source of disease to wild populations as animals contact each other across fence lines and possibly escape their enclosures (e.g., https://www.aphis.usda.gov/sites/default/files/status-of-captive-herds.pdf). Transportation of live animals between captive herds has led to long-distance CWD dispersal events and outbreaks that are far from previously known cases. In addition, there is concern among captive cervid owners that infected wild deer could transmit CWD to their herds, which could result in a requirement to depopulate a herd. Finally, Vermonters who hunt out of state, taxidermists, and processors could introduce CWD through the disposal of infected cervid parts on the landscape.

Chronic wasting disease dynamics and the efficacy of management interventions are only partly understood. There is uncertainty about the relative risk of introduction and spread associated with different pathways, including but not limited to, captive cervid facilities, out-of-state hunting, and taxidermists and processors. There is uncertainty about the level of compliance with existing regulations (e.g., bans on natural lures and baiting), and, to a lesser extent, about the rate of deer migration and movement. Each preventative or responsive management action has uncertainty around the effectiveness of the action owing to a combination of action efficacy and compliance. The slow development of the disease has confounded the ability of scientists to resolve many of these uncertainties.

#### 3.1.2 Legal, Regulatory, and Political Context

Under U.S. laws, the authority and responsibility to manage terrestrial wildlife belongs to the states, except for species that are listed under the Federal Endangered Species Act of 1973 (16 U.S.C. §1531 et seq.). In Vermont, much of the relevant statutory authority is found in Title 10 of the Vermont Statutes Annotated, particularly in Chapter 103, which establishes the Department of Fish and Wildlife (10 V.S.A. §4041 et seq.). The VDFW manages and protects free-ranging wildlife under Vermont law. “The State, through the Commissioner of Fish and Wildlife, shall safeguard the fish, wildlife, and fur-bearing animals of the State for the people of the State, and the State shall fulfill this duty with a constant and continual vigilance” (10 V.S.A. §4081(a)(2)). Deer are specifically mentioned in the statutory mission of VDFW, “An abundant, healthy deer herd is a primary goal of fish and wildlife management” (10 V.S.A. §4081(c)). The VDFW has broad powers to promulgate and enforce regulations meant to conserve wildlife and provide recreational and hunting opportunities to the citizens of Vermont.

The VAAFM is established under Title 6 of the Vermont Statutes and is implicated in CWD management because deer are kept in fenced enclosures for meat production, recreational viewing, and as a hobby. Broadly, “The Secretary [of Agriculture, Food and Markets] shall promote the agricultural interests and education throughout the state” (6 V.S.A. §3). Among a wide set of authorities, VAAFM oversees captive deer operations. The Rules Governing Captive Cervids require CWD testing of any carcasses, annual inventories, redundant animal identification systems, and sufficient facilities to ensure containment and safe handling of herds (6 V.S.A. §1165). In addition, VAAFM can quarantine domestic animals or premises if they have been affected by or exposed to contagious disease (6 V.S.A. §1157) and may condemn or order destroyed infected or exposed animals (6 V.S.A. §1159). The movement of domestic animals into and out of the State is also regulated by VAAFM (6 V.S.A. §1461), which maintains strict standards for captive cervids.

The VDH has “power to supervise and direct the execution of all laws relating to public health […]” (18 V.S.A. §1). This includes identifying any public health hazards and mitigating their risk (18 V.S.A. §2), as well as managing communicable diseases (18 V.S.A. §§1001-1141). The Department collaborates with other health agencies, like CDC, on matters of human health that cross State boundaries. Chronic wasting disease infections have not been detected in humans. Current public health recommendations include refraining from handling and consuming animals that look sick, act strangely, or are found dead, and strongly consider testing deer and elk meat prior to consuming if the hunt occurred in an area with CWD activity (Centers for Disease Control and Prevention, 2026).

The VFPR was created under Title 10, Chapter 83 of the Vermont Statutes Annotated (10 V.S.A. §2600) and is responsible for “manag[ing] and plan[ning] for the use of publicly owned forests and park lands” (10 V.S.A. §2603 (b)). The VFPR assists private landowners and lumber operators with forest management and can support VDFW in gaining access to future locations affected by CWD as well as understanding the role that deer have in altering forest structure and composition.

In terms of Federal agency involvement, the USDA Animal and Plant Health Inspection Services (USDA-APHIS) Wildlife Services Program’s statutory authority comes from the Act of March 2, 1931 as amended (46 Stat. 1468-69; 7 U.S.C. §§8351-8352), and the Act of December 22, 1987 (Public Law No. 100-202 §101(k), 101 Stat. 1329-331, 7 U.S.C. §8353). The USDA-APHIS Wildlife Services has expertise in culling wild animals and can contract with State agencies to undertake a reduction in a wild deer population. The USFS, within the USDA, sustains healthy forests (16 U.S.C. §472). A major landowner in the state of Vermont, USFS manages the Green Mountain National Forest, which was created in 1932 and encompasses more than 400,000 acres.

The USFWS manages the National Wildlife Refuge System. The Mission of the Refuge System is “To administer a network of lands and waters for the conservation, management, and where appropriate, restoration of the fish, wildlife, and plant resources and their habitats within the United States for the benefit of present and future generations of Americans” (National Wildlife Refuge System Improvement Act of 1997, Public Law 105-57). Units and divisions of the Missisquoi National Wildlife Refuge and Silvio O. Conte National Fish and Wildlife Refuge are in Vermont. A wide variety of laws, regulations, executive orders, and policies dictate how National Wildlife Refuges are administered. Key concepts and guidance for the System are included in the National Wildlife Refuge System Administration Act of 1966, The Refuge Recreation Act of 1962, Title 50 of the Code of Federal Regulations, the Fish and Wildlife Service Manual, Executive Order 12996 (March 23, 1996) and, most recently, the National Wildlife Refuge System Improvement Act of 1997.

#### 3.1.3 Current CWD Status

As of December 2025, CWD has not been detected in Vermont. There was a multi-animal detection in a single captive facility in Quebec, Canada, in 2018, and a small outbreak in captive and wild deer in eastern New York in 2005, but both have been contained and eliminated. Additionally, there was a detection in a captive facility in Herkimer County, New York in 2024 that is believed to be contained. As of 2025, the closest-known infected wild population is approximately 225 miles away in Luzerne and Carbon Counties, PA, whereas the closest known captive population is in Wayne County, PA, which is 125 miles away from southwestern Vermont (fig. 1).

**Figure 1.**
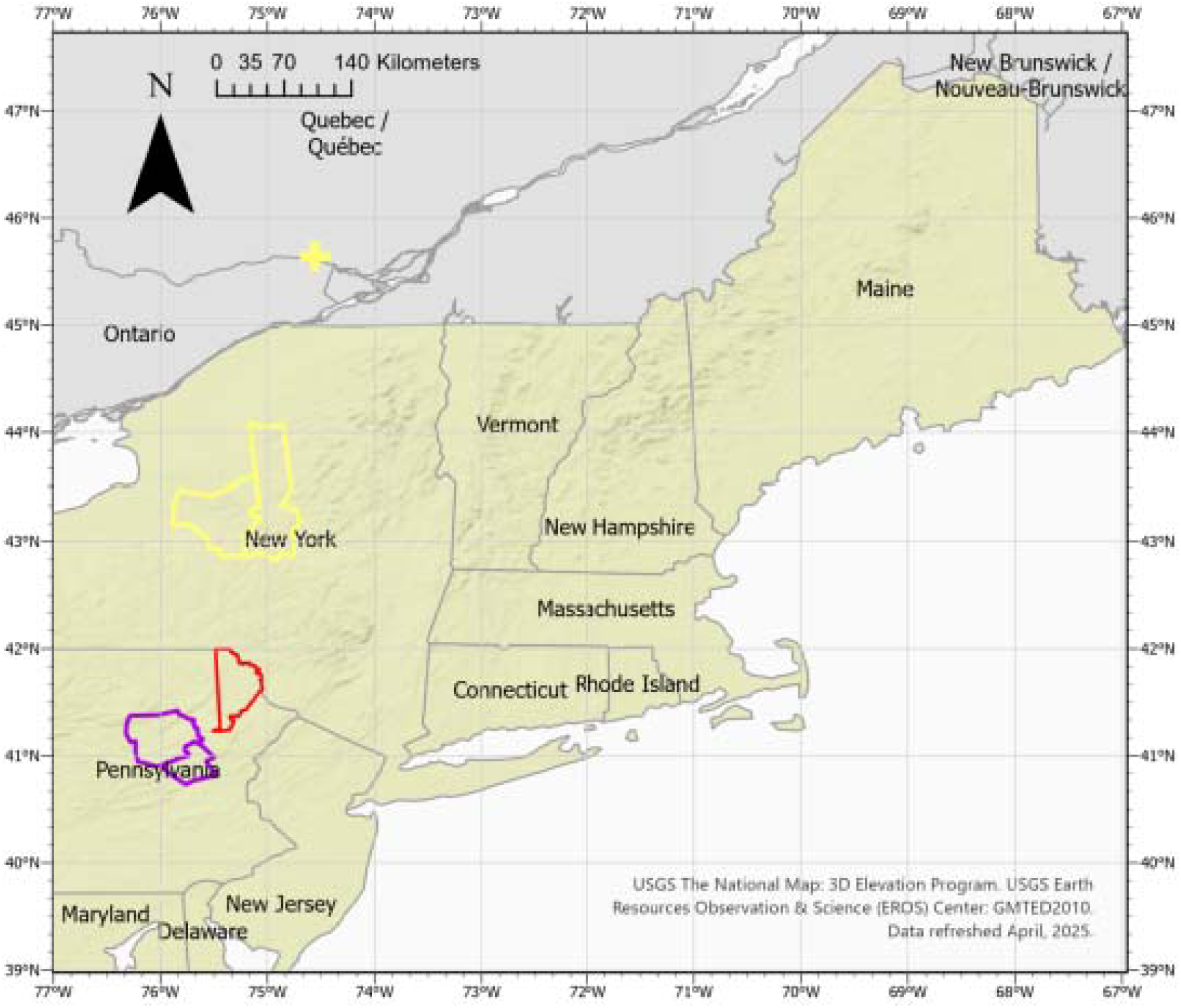
Map of the northeastern United States including Vermont, the nearest active chronic wasting disease (CWD) positive U.S. counties for captive cervids (red outline) and free-ranging deer (purple outlines). Also included are prior detections that are believed to be no longer spreading, including positive captive facilities in Herkimer County, NY and Oneida County, NY (yellow outlines) and another captive facility in Quebec, Canada (yellow cross).

The current CWD surveillance protocol in Vermont is to test suspect animals reported by the public; the number of samples varies but averages less than 10 animals per year. Clinical-suspect animals are those that display behaviors and condition that are typical of disease, including loss of fear of humans, appearing disoriented, and appearing emaciated. Current education and outreach efforts related to CWD consist of 1−2 pages in the Hunting Regulation Digest and one webpage (https://www.vtfishandwildlife.com/learn-more/living-with-wildlife/wildlife-diseases/chronic-wasting-disease). At the time of this report there was no active in-person outreach campaign, but in 2019, the VDFW organized presentations around the State. The Fish and Wildlife Board has a rule that there is to be no importation of carcasses or high-risk parts (i.e., head or spinal cord) from wild deer or elk harvested in CWD-positive jurisdictions, or from captive deer or elk from farms or hunt facilities (10 Appendix V.S.A. §17). In addition, VDFW prohibits the use of natural urine or any bodily fluid-based lure to attract deer while hunting, and no baiting or feeding of deer is allowed (10 Appendix. V.S.A. §37). VDFW wardens are charged with enforcing the bans and regulations.

In captive deer, VAAFM has actively managed the risk of CWD introduction into captive deer facilities through annual inventories and inspections of the existing facilities, implementing redundant forms of individual animal identification that allow individual tracking, and testing all mortalities for CWD (VAAFM, 2005). Additionally, VAAFM has strict guidelines about animal importation, including a valid veterinary inspection certificate, requirements that the originating location is free from CWD in free-ranging and captive deer, and that the receiving facility has a history of compliance with regulations and a strong justification for importation. If CWD were to be detected in a captive facility, VAAFM protocols include immediate depopulation of all CWD-susceptible species in a facility, implementation of procedures to clean and restrict use of the potentially contaminated lands, and conducting epidemiological tracing to limit spread in other facilities (if necessary).

## 4 Decision Analysis

The management of CWD is complex and requires decision makers to navigate challenges such as uncertainty in statutory authorities and disease dynamics, and tradeoffs in the ability to address all management objectives simultaneously. The use of decision analysis can help overcome these challenges by clarifying the scope, scale, and specific detail of management. It also invites articulation of the full range of concerns held by one or more decision makers in a manner that guides formal evaluation and detailed deliberation. Structured decision making, a normative process for conducting decision analysis, has been used successfully in natural resource settings (Runge and others, 2020), including applications to wildlife diseases (McEachran and others, 2024), including CWD (Cook and others, 2023; Cook and others, 2025). This section describes the basic components of decision analysis applied to CWD management in Vermont.

### 4.1 Structured Decision-Making

Structured decision-making is practiced using five steps: describing the **pr**oblem, articulating the **o**bjectives, developing **a**lternatives, evaluating the **c**onsequences, and navigating **t**radeoffs (PrOACT; Gregory and others, 2012). The problem describes the details of the decision to be made, the decision maker and their authority, as well as the spatial and temporal extent. The objectives articulate the values of the decision maker and stakeholders that are important to consider. Each objective can be quantified by a performance metric, which captures how it performs under the actions being considered and is fully described by an appropriate scale, units, and direction. The alternatives are the discrete options from which the decision maker can choose. In natural resource management, the alternatives are often quite complex strategies themselves, consisting of many component actions in various combinations. Following a description of the problem, objectives, and alternatives, the next step is to formally evaluate the performance of the alternatives against all the objectives, using the best available science. Then, deliberations can focus on specific performance of alternatives and can take advantage of analytical tools that simplify this task, such as multi-criteria decision analysis (MCDA).

### 4.2 Multicriteria Decision Analysis

Multi-criteria decision analysis is a branch of decision analysis that helps a decision maker navigate competing objectives under different alternatives. For example, while an aggressive disease management response may perform well at keeping prevalence low, it might suffer from large financial expenditures. In contrast, there may be other alternatives that perform poorly on disease but are more financially feasible. Without the use of deliberative tools, like MCDA, it is difficult to decide among those strategies. Multi-criteria decision analysis tools provide a transparent way to navigate differences, articulate values positions, and select the appropriate course of action.

## 5 Vermont CWD Management Plan Framework

To develop the framework for a state-wide CWD management plan, we used tools from decision analysis and quantitative assessment in an iterative prototype approach, increasing the sophistication of the analysis with each prototype until a satisfactory framework emerged. We began by first framing the CWD management problem collaboratively, including a shared problem statement, a set of objectives, and a draft set of actions. We then used a series of quantitative and decision analysis techniques to illustrate and navigate necessary tradeoffs across a range of management strategies. During the deliberation phase, we crafted alternatives using two approaches (sections 5 & 6): one that focused on a discrete set of six contrasting alternatives; and another that sought to optimize a larger set of alternatives for a more limited set of objectives.

### 5.1 Problem Statement

Chronic wasting disease, a fatal disease of cervids, is spreading across North America. While it has been detected in the northeastern region of the U.S., it has not yet been detected in Vermont. The Vermont Department of Fish and Wildlife, along with the Vermont Department of Forests, Parks and Recreation, the Vermont Agency of Agriculture, Food and Markets, the Vermont Department of Health, the U.S. Department of Agriculture, the U.S. Fish and Wildlife Service National Refuge System, and the U.S. Forest Service, desire to have a robust prevention and response plan that identifies actions that reduce the risk of CWD introduction into free-ranging (white-tailed deer and moose) and captive (including red deer and fallow deer) cervid populations in the State, and to reduce the effects of CWD once it arrives. The intent of VDFW is to build on a CWD Response Framework drafted in 2019 and include proactive, reactive, and surveillance measures that can be taken cooperatively by participating agencies to best meet their collective objectives. The agencies plan to use a deliberative structured decision-making process and cover statewide strategies evaluated across 25- and 50-year time horizons.

### 5.2 Objectives and Performance Metrics

The framework included two strategic objectives and one process objective. Strategic objectives are broadly defined and help describe the general mission and goals of an agency. Process objectives help describe important aspects of how decisions are made.

**Strategic Objective 1**. Ensure public awareness about diseases that affect (or may affect) humans, plants, and animals of Vermont.
**Strategic Objective 2.** Maintain public satisfaction in the ability of agencies to manage deer that are healthy and in balance with their ecosystems.
**Process Objective 1.** Develop a multi-agency CWD prevention and response plan that considers all objectives and clearly defines actions, including roles and responsibilities of partners and stakeholders.

The fundamental objectives are the key values driving the decision-making process. In total, there were seven fundamental objectives and eight performance metrics used to measure the performance of alternatives (table 1).

**Table 1.** Table of seven fundamental objectives and eight performance metrics used in the decision analysis applied to the management of chronic wasting disease (CWD) in Vermont.

| Fundamental objective number | Fundamental objective | Performance metric number | Performance metric |
| --- | --- | --- | --- |
| 1 | Maximize herd health of native cervid populations in the state of Vermont | 1a | Size of deer population in 25 and 50 years |
|  |  | 1b | CWD prevalence in 25 and 50 years |
| 2 | Maximize the public benefit of cervids (consumptive and non-consumptive) | 2 | Average annual number of active hunters over 25 and 50 years |
| 3 | Minimize harm to other species, unique ecosystems, and forests | 3 | Density of deciduous saplings at year 25 and 50 (saplings/hectare) |
| 4 | Minimize costs, staff time, and workload associated with CWD management | 4 | The average cost per year over 25 and 50 years |
| 5 | Minimize the effect (or potential effect) that CWD has on public health | 5 | The total number of CWD positive deer harvested over 25 and 50 years |
| 6 | Minimize the effect of CWD and its management on local economies | 6 | Average annual net revenue for hunting-related businesses over 25 and 50 years |
| 7 | Maintain opportunities for private landowners to engage in agricultural pursuits | 7 | Cumulative captive cervid farm-years over 25 and 50 years |

**Fundamental Objective 1.** *Maximize health of native cervid populations in Vermont.*—Deer management is codified by Vermont statute and requires VDFW to manage for an abundant and healthy deer herd as a primary goal of management (10 V.S.A. §4081(c)). Further, VDFW aims to “maintain the deer population at levels that are socially acceptable and ecologically sustainable” and to “maintain a healthy moose population in Vermont’s moose management regions” (Vermont Fish and Wildlife Department, 2021b). Chronic wasting disease and its management have the potential to affect the health and abundance of white-tailed deer and moose in Vermont. Performance metric 1a.— The size of the white-tailed deer population in Vermont in 25 and 50 years. Performance metric 1b.— The prevalence of CWD in white-tailed deer at year 25 and year 50. While the State was interested in both white-tailed deer and moose, we only estimated the effects of management strategies on deer and assumed that any benefits of reduced prevalence would also benefit moose.
**Fundamental Objective 2.** *Maximize the consumptive and non-consumptive benefits of native cervids.*— Public perception about deer density can be based on individual preferences, recreational or economic interests, and environmental concerns. Hunting contributes to rural economies, is the primary source of funds for wildlife conservation, and has a rich history in Vermont. The VDFW manages deer to maintain hunting opportunities and satisfaction by, “Provid[ing] a quality deer hunting experience for as many hunters as possible” (Vermont Fish and Wildlife Department, 2021b). Wildlife viewing is an activity that Vermont residents and nonresidents value; maintenance of this opportunity is important to the State agencies. Performance metric 2.— The average annual number of active Vermont deer hunters over 25 and 50 years.
**Fundamental Objective 3.** *Minimize harm to other species, unique ecosystems, and forests.*— The management of Vermont’s lands, waters, and species is accomplished by State and Federal agencies. The VDFW’s mandate necessitates wildlife and habitat conservation. Similarly, VFPR, USFWS, and USFS conserve species, ecosystems, and forests. Chronic wasting disease and actions taken to manage it can have direct and indirect effects on other species, lands, and waters. One of the major ways that deer affect other species is through their browsing effects on forest structure and composition. Performance metric 3.— The density of deciduous saplings, defined as trees with diameter at breast height greater than one centimeter and less than nine centimeters, at year 25 and year 50.
**Fundamental Objective 4***. Minimize costs, staff time, and workload associated with CWD and its management.—*Actions to manage deer and CWD can be labor intensive and costly. Agencies must operate within budgetary constraints and allocate State and Federal resources efficiently and effectively. Any staff and financial costs allocated to the management of CWD have a direct effect on agency budgets and could redirect staff time to CWD management in a manner that reduces resources available to other agency programs. Performance metric 4.— The average annual cost (in 2024 U.S. dollars) of CWD management to Vermont agencies over 25 and 50 years.
**Fundamental Objective 5***. Minimize the effect (or potential effect) that CWD has on public health.—* CWD infections in humans have not been reported. If humans can be infected with CWD, consuming infected meat would be the likely route of transmission. To reduce possible risk, CDC and jurisdictions with CWD activity recommend precautions including testing meat prior to consuming it and not hunting and consuming animals that appear sick. In Vermont, the harvest and distribution of venison by hunters provides approximately 3 million servings of protein per year. Hunters also donate venison through the “Venison for Vermonters” program. A decrease in the deer population, and subsequently a decrease in the sharing of venison, could affect food security. Performance metric 5.— The total number of CWD-positive deer available for human consumption (that is, CWD-positive deer that are harvested but not tested for CWD within the simulated population) over 25 and 50 years.
**Fundamental Objective 6***. Minimize the effect of CWD and its management on local economies.—* Many businesses, such as those that operate check stations, process carcasses for meat, prepare deer for mounting, or maintain captive cervid facilities, are affected by deer management. Deer harvest check stations are closely tied to the hunting community and provide marketing opportunities for participating businesses. Likewise, processors and taxidermists rely on hunters to generate income. Larger captive cervid facilities in the State have economic interests in meat production. Changes in CWD and its management may affect deer harvest and the number of hunters and could lead to restrictions that affect businesses. Performance metric 6.— Average annual net revenue for hunting-related businesses over 25 and 50 years.
**Fundamental Objective 7***. Maintain opportunities for private landowners to engage in agricultural pursuits.—* Private landowners engage in a diversity of agricultural activities on their lands, including the holding of captive cervids for economic and recreational opportunities. In Vermont, captive cervid facilities can range from larger scale (100−200 individual deer) to smaller scale (<10 individuals) operations. While some facilities may have economic interests in meat production, other facilities and landowners enjoy hobby farming. VAAFM seeks to support the right of Vermonters to engage in diverse agricultural pursuits, including captive cervid farming. Performance metric 7.— Cumulative captive cervid farm-years in the next 25 and 50 years (calculated as the summed product of the number of captive cervid farms multiplied by the time that each farm is operational).

### 5.3 Actions

#### 5.3.1 Overview

To develop a set of well-defined CWD management strategies to evaluate, we first conducted a series of unstructured brainstorming sessions intended to promote creative thinking. Participants included agency staff with technical knowledge about their programs and the potential effects of CWD; and therefore, discussion focused on aspects related to wild and captive cervid management, forestry practices, human public health risks and messaging, and agency budgets. We then organized those actions into distinct categories and further subdivided the brainstormed list to include those actions that could be taken prior to detection (preventative or proactive actions), after detection (responsive or reactive actions), or as part of detection (i.e., surveillance). We consider surveillance as a separate category of actions from proactive or reactive actions because it serves as a source of information that can direct or trigger targeted actions, but in isolation does not alter disease dynamics.

#### 5.3.2 Proactive Actions

We considered a suite of proactive actions spanning major themes including hunting regulations and population management, carcass management, captive cervid management, outreach and education, and agency coordination. Within hunting regulations and population management actions, VDFW, with support of the other agencies, can act to change deer harvest by sex and age, methods of take, length of season, and access, in anticipation of the arrival of CWD to the state. They can also direct targeted actions to reduce local deer density in locations where hunting access might be limited. These actions would be intended to lower direct and indirect contacts of deer by reducing density in locations of possible CWD emergence.

For carcass management, VDFW, again supported by other agencies, can develop protocols that reduce the likelihood that infectious carcasses or tissues end up in locations where Vermont deer and moose can contact them. These actions are designed to reduce the introduction of CWD via improper disposal of out-of-state carcasses. Lastly, VAAFM can alter their already stringent captive cervid regulations to further restrict the importation of CWD-susceptible species, require double fencing of facilities, or move to close existing facilities within the state. They could also restrict importation of feed or supplies, however most of these resources are already acquired within the State. This set of actions might reduce the potential arrival and spread of CWD via the captive cervid industry. All these actions have the potential to be further enhanced by outreach and education, and agency coordination.

#### 5.3.3 Reactive Actions

The reactive actions mostly followed from the same set of categories as the proactive actions but were triggered after the detection of CWD. For example, changes in hunting regulations, carcass management, and captive cervid management would act to reduce the risk of CWD spread, albeit targeted to locations surrounding recent detections within Vermont’s borders. For the purposes of this evaluation, we assumed that reactive actions were triggered upon the detection of disease within the State, however, in practice, the State could implement reactive actions when the disease moves within a pre-determined distance from Vermont’s border. Additionally, the context surrounding outreach and education and agency coordination would change in comparison to pre-detection strategies, but the elements would remain the same.

#### 5.3.4 Surveillance Actions

To be effective, reactive actions need to be coupled to a program of surveillance that can detect CWD at an appropriate stage and provide any information that is necessary to direct actions; however, surveillance is one of the costliest CWD-related actions that an agency can take (Chiavacci, 2022). As a result, there are a range of surveillance designs that are possible and that may improve the efficiency of sample collection, storage, diagnostic, and database management costs. Instead of a detailed assessment of surveillance design, we focused on questions about detection probability as a function of sample size, and how successful detection affected the performance metrics of interest. We accounted for the expense of surveillance by including a per-sample cost that was derived from other large-scale CWD testing programs. We expect that future iterations of the CWD management planning process in Vermont will further evaluate questions surrounding the exact approach necessary to collect appropriate sample sizes.

## 6 Alternative CWD Management Strategies – Phase I

### 6.1 Developing Strategies

A CWD management plan is a combination of many different actions, including proactive actions, surveillance activities, and reactive actions. The development of a management plan, then, is the choice among all the possible permutations of the individual actions. To evaluate approaches to CWD management, we developed alternative *strategies*, that is, alternative portfolios of actions. In Phase I, we developed a small set of alternative strategies that were guided by differing prevention and response philosophies designed to span the decision space. In Phase II, we examined a more systematic set of alternative strategies.

### 6.2 Alternative Strategies

In the first phase of analysis (Phase I), the technical team developed six distinct alternatives from the set of actions available. The alternatives were designed to elicit strong contrasts in the performance of the objectives under the actions and how surveillance supported that outcome. As such, we designed alternatives to focus on lax and aggressive variations in proactive, reactive, and surveillance actions in differing combinations.

#### 6.2.1 Status Quo

The *Status Quo* alternative embeds the actions that Vermont agencies are already taking. These actions include existing VDFW prohibitions on baiting, naturally derived urine lures, and imports of carcasses from CWD-positive jurisdictions; VAAFM inspections of captive facilities, import restrictions of live cervids, and processes for depopulation, facility closure, and decontamination in the event of a CWD-positive captive animal; and VDH distribution of public health guidance. Surveillance for CWD by VDFW is minimal under the status quo (∼5−10 clinical-suspect deer per year); this strategy assumes surveillance would increase to provide voluntary testing of carcasses after first detection, as would hunting opportunity (to reduce deer density in CWD-affected areas).

#### 6.2.2 Heavy Outreach

The *Heavy Outreach* alternative is focused on outreach and education efforts before and after CWD detection that emphasize important information about CWD but otherwise have limited actions that act directly on disease dynamics. By developing and disseminating outreach materials, Vermont agencies would provide pertinent information that may inform voluntary action by hunters and the public, but would maintain status quo conditions for harvest, carcass disposal, and cervid management. Additional resources would be provided for voluntary CWD testing and additional sampling to ensure that the information shared with the public is current and accurate.

#### 6.2.3 Reduce Risk of CWD Arrival

The *Reduce Risk of CWD Arrival* alternative emphasizes a suite of actions that occur before CWD is detected in Vermont. The goal is to implement aggressive preventative actions that are intended to delay the arrival of CWD as long as possible. The primary mechanisms whereby these preventative actions occur is by reducing the risk associated with hunter-mediated carcass movements; enhancing guidelines for the proper disposal of carcasses by hunters, taxidermists, and processors; and limiting the potential of captive-to-wild deer transmission. Surveillance under this strategy would be limited to voluntary testing only because there are few reactive measures planned.

#### 6.2.4 Prepare and React to CWD

The *Prepare and React to CWD* alternative creates opportunities for an aggressive reactive response to CWD detection and then implements actions targeting hunter harvest, carcass management, cervid management, outreach and education, and interagency coordination at first detection. Because the efficacy of response depends on early detection of CWD and information to target ongoing responses, there is also a large investment in surveillance under this alternative. Pre-detection actions are limited to regulatory changes that allow for additional post-detection options in addition to other preparatory actions that allow for an intensive response.

#### 6.2.5 Prevent and React (Moderate Intensity Management Before and After Detection)

The *Prevent and React* alternative calls for moderate investments in proactive and reactive resources and to support these efforts through intermediate expenditures in CWD surveillance. There are increased regulations intended to: delay the arrival of CWD from carcass and live-animal transport; increase hunting access before CWD detection; and reduce deer density after detection. Finally, there are supporting efforts to increase outreach and education to hunters, members of the public, and captive cervid facility owners.

#### 6.2.6 Kitchen Sink (Maximum Intensity Management Before and After Detection)

The *Kitchen Sink* alternative calls for all agencies to implement the most aggressive proactive and reactive actions that they can. The goal is to explore the efficacy of an alternative that takes all possible actions, and to understand the positive and negative implications of such an alternative on the suite of management objectives. The *Kitchen Sink* strategy would invest heavily in surveillance so that the agencies could implement post-detection actions early after the arrival of CWD.

## 7 Alternative CWD Management Strategies – Phase II

During Phase I of the project, the six alternative strategies were evaluated against the eight performance metrics, and the results were presented to representatives from all six agencies that were collaborating on the CWD management plan. During this meeting, agency decision makers were presented with model descriptions, quantitative results, interpretations, and a consequence table that provided a visual representation of emerging findings. The participants were then guided through an MCDA exercise that elicited important values-based feedback that allowed for additional targeting of any pressing questions. What emerged from the meeting was a central question about whether there were other combinations of actions, beyond the six alternatives analyzed, that would perform better across performance metrics, particularly cost and CWD prevalence. Within that central question, there were a series of related topics of inquiry, specifically, how important are proactive, reactive, and surveillance efforts at reducing long-term disease prevalence; how efficiently can the important actions be accomplished; and, finally, how important was captive cervid farming to disease in wild deer populations.

To evaluate those pressing questions, we expanded the range of alternatives considered into a set of 256 strategies that allowed for a systematic evaluation of important tradeoffs. The first set of 40 scenarios (the “proactive set”) investigated the contributions of proactive actions at two different levels of reactive actions.

The proactive actions were:

- Captive management (5 levels): status quo (captive farms open, no double-fencing, maintain rate of captive imports); reduce rate of captive imports only; proactively double-fence all farms; double-fence and reduce captive imports; proactively close all captive farms.
- Carcass import restrictions (2 levels): status quo; further restrict and enforce import of carcasses by Vermont residents hunting out-of-state.
- Carcass disposal guidance (2 levels): status quo; increased guidance to hunters, taxidermists, and meat processors on carcass disposal practices that reduce the risk of CWD transmission to Vermont deer.

The two reactive settings for this set of proactive scenarios were:

- Passive reaction to a new detection (pre-detection surveillance of 10 carcasses per year, no sharpshooting, no increase in harvest rate post-detection).
- Aggressive reaction to a new detection (pre-detection surveillance of 463 carcasses per year, 250 deer/year removed by sharpshooting post-detection, increase harvest rate by 20% post-detection).

The second set of 216 scenarios (the “reactive set”) examined the contributions of pre-detection surveillance and post-detection responsive actions at three levels of proactive actions. The surveillance and reactive actions were:

- Pre-detection surveillance (4 levels): 10, 250, 463, or 1,000 carcasses sampled per year. The levels were calculated as: the maximum number of clinical-suspects that Vermont tests annually (10 per year); approximately half of the level required to detect CWD at 1% prevalence with 99% certainty (250); the level required to detect CWD at 1% prevalence with 99% certainty plus 5 clinical-suspect animals (463); and double the level required to detect CWD at 1% prevalence with 99% certainty (1,000).
- Post-detection harvest rate (3 levels): status quo (based on 2024 harvest regulations); 10% increase of all sex/age classes; 20% increase of all sex/age classes.
- Post-detection sharpshooting (6 levels): 0, 50, 100, 250, 750, and 1,500 deer removed per year in the area around the positive detections.

The three proactive settings for this set of reactive scenarios were: low (status quo captive management, status quo carcass import restrictions, status quo carcass disposal guidance); medium (restrict captive import, restrict carcass import, increase carcass disposal guidance); high (proactively close all captive facilities, restrict carcass import, increase carcass disposal guidance).

We then used Pareto-based methods to identify alternatives that were Pareto-efficient and used those to further deliberate the tradeoff between cost and CWD prevalence under each of the 256 scenarios. Pareto efficiency occurs when an improvement in one objective cannot occur without a loss on another objective; these efficient strategies essentially require a decision maker to then determine how much any gains are worth (in the currency of the competing objective; Runge and others, 2020; Williams and Kendall, 2017).

## 8 Quantitative Methods

To evaluate the suites of management strategies in Phases I and II against the eight performance metrics, a set of interconnected predictive models were built. The development of the models arose out of the decision framing elements described in sections 5–7, and coordination with a technical team made up of professionals with expertise in wild deer behavior and movement, population dynamics including harvest, CWD dynamics, public health, human dimensions, and agricultural practices. This section describes the methods, including a description of each of the modeling approaches, and how they were linked to estimate outcomes of interest. Each alternative was analyzed with 1,000 replicates, and the results were summarized cumulatively across time or at 25- and 50-year endpoints, depending on the performance metric.

### 8.1 Modeling Overview

In combination, the models simulate the spread of CWD across the northeastern U.S. from its currently known distribution in Pennsylvania, the arrival of CWD to and subsequent growth in Vermont deer, and ultimately the effects of CWD on the performance metrics (fig. 2). Also included was an observation process whereby VDFW conducts CWD surveillance to detect disease arrival and then switches to the set of reactive management actions upon first detection. The model structures relied on existing literature, conversations with members of the technical teams, and consultation with outside experts. To inform the necessary set of parameters within the models, we used existing data from literature as well as expert judgment. Because of limited evidence for some parameters, particularly disease transmission rates and management efficacy, we further confirmed the performance of models through a series of meetings with the technical experts.

**Figure 2.**
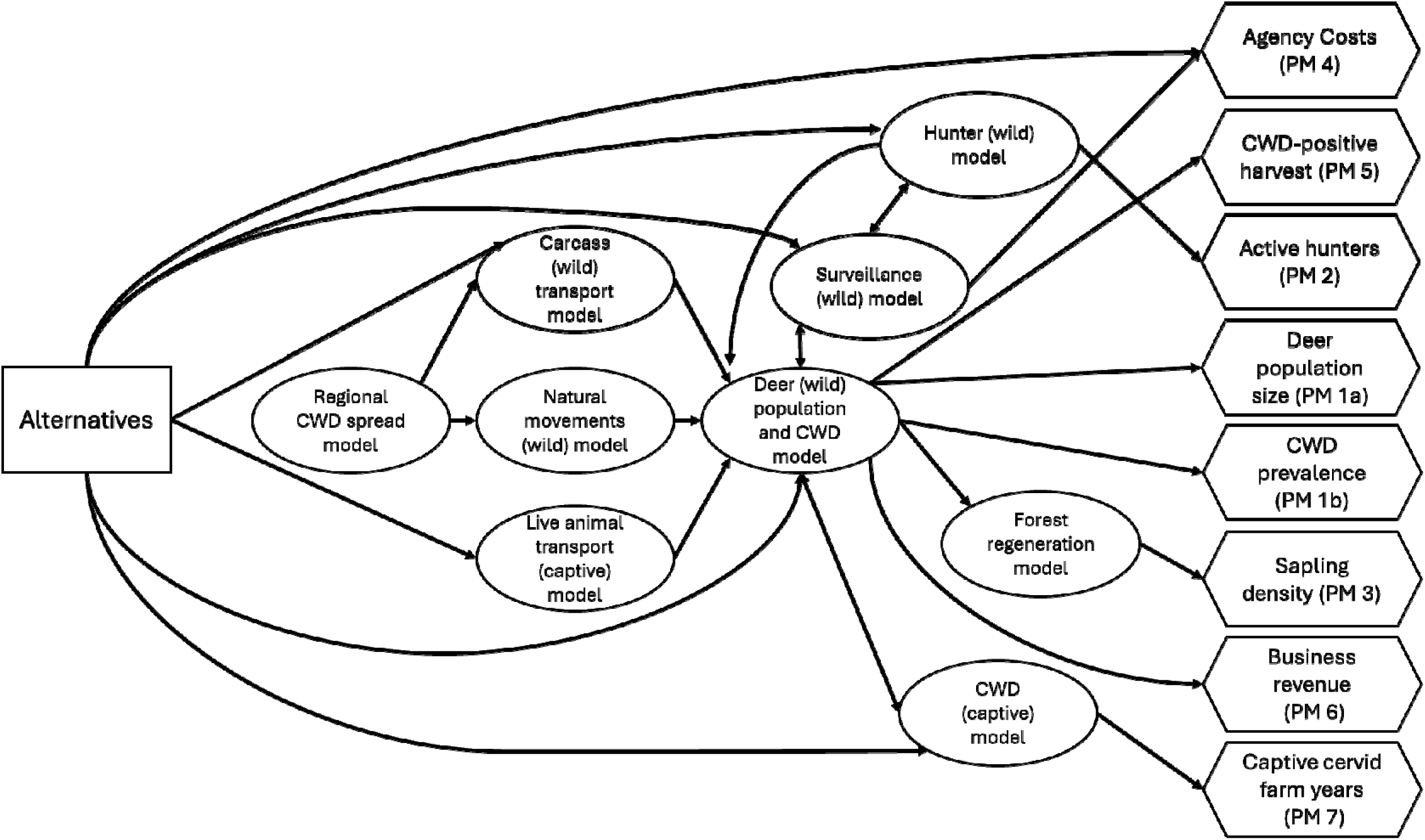
Computer modeling schematic, including alternatives (rectangle), models of dynamic processes (ovals), and connections to performance metrics (hexagons). [CWD, chronic wasting disease; PM, performance metric]

### 8.2 Regional CWD Spread and Arrival in Vermont

#### 8.2.1 Overview

Vermont is currently believed to be free from CWD (VDFW, 2026). However, the disease continues to spread in wild and captive deer populations across North America and could arrive in the State through several pathways. We considered three primary pathways: the importation and disposal of infected carcasses of hunter-harvested wild deer; natural deer movements that lead to geographic spread and eventual arrival in Vermont; and finally, the importation of a live captive cervid into one of the nine existing farms in Vermont. While it is possible that other pathways contribute to the spread of CWD (e.g., contaminated equipment, feed, or other fomite), these routes were considered to be lower risk and therefore were not considered further.

The total number of wild Vermont deer newly infected by the three pathways annually (*I_intro,t_*), was thus given by

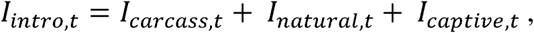

where *I_carcass,t_* is the total number of new wild Vermont deer infections in each year *t* from CWD-infected carcasses imported from outside of the state; *I_natural,t_* is the total number of new wild Vermont deer infections from live, CWD-infected deer moving into Vermont; and *I_captive,t_* is the total number of new wild Vermont deer infections from fenceline transmissions between susceptible wild deer and infected captive deer.

#### 8.2.2 Regional CWD Spread Dynamics

We modeled spread and growth across the entire northeastern U.S. (Connecticut, Massachusetts, Maine, New Hampshire, New Jersey, NY, PA, Rhode Island, VT) on an annual time step using a hexagonal spatial structure with a cell size of 1,040 km^2^ (34.7 km between cell centers). We chose this area based on rough agreement with the maximum dispersal distance of white-tailed bucks (Diefenbach and others, 2008). We then used localized apparent prevalence estimates from 2024 Pennsylvania surveillance data as initial conditions in each of the spatial units (Pennsylvania Game Commission, 2026). The county-level surveillance data were reclassified to match the spatial units of the model by calculating the proportion overlap between each hexagonal grid cell and Pennsylvania counties and then taking a weighted average from that overlap.

After estimating initial prevalence, we then simulated local growth of disease within units as a logistic process, with a diffusion process describing the subsequent spread of disease across unit boundaries.

The logistic growth process was given by

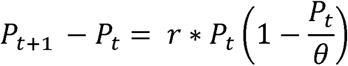

where

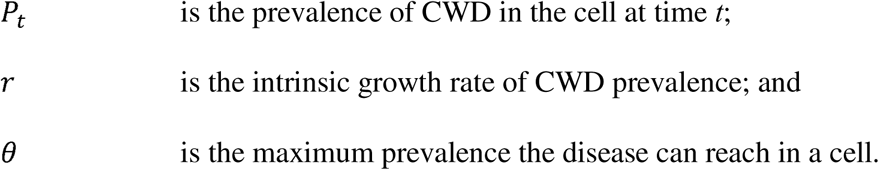

To represent uncertainty about the maximum CWD prevalence (*θ*, an asymptote) that can be reached in each spatial unit, we considered *θ* to be a random variable with a uniform distribution between 0.4 and 0.5 (Samuel, 2023). For the growth rate (*r*), our expectation for how quickly CWD would grow within spatial units was based on empirical data from locations with a long history of CWD surveillance (e.g., Samuel, 2023; minimum 0.1 and maximum 0.6).

To represent the spread of disease across different spatial units, our diffusion model was given by

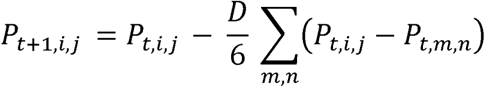

where,

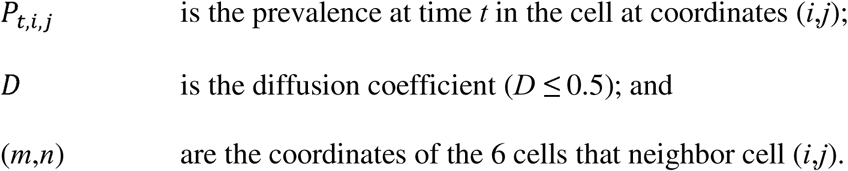

Adapted from Astor and Adami (1998), the above equation shows the updated prevalence for each cell after each time step. This is found by taking the difference between the starting prevalence value and the net diffusion change. Because our cells are hexagons, there are typically six neighboring cells for the calculation. We set two boundary conditions: 1) for a hard boundary such as the Atlantic Ocean along the east coast, we allow prevalence to accumulate and rebound back into the landscape, and 2) for soft boundaries which were artificially created by the spatial extent of our problem, such as the border between Canada and the U.S., we allow the disease to diffuse out of the study area. We assume that diffusion is constant across the landscape and set D as a uniform distribution between 0.05 and 0.4.

#### 8.2.3 Wild Deer Movements

To model the introduction risk from wild deer movements, we developed an approximation to the full spatial spread model that was computationally more tractable to integrate into the rest of the simulation. Consider a line that connects the leading edge of CWD infection in wild deer and the border of Vermont and that represents the shortest distance between them. The distance along this line, *x* (in kilometers), represents the distance from Vermont (with *x* = 0 located on the Vermont border). At time 0 of the simulation, let the leading edge of CWD infection be located at *x* = *L*(0), where the leading edge is defined as the place where the CWD prevalence is half of the potential maximum prevalence. We expect that the position of the leading edge will decrease over time, as CWD spreads across the landscape. The location of the leading edge over time is represented as *L*(*t*) and described by

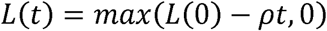

where the rate of spread, *ρ*, describes how quickly the leading edge advances toward Vermont.

The CWD prevalence along the line between the leading edge and the Vermont border is described with a logistic function, such that

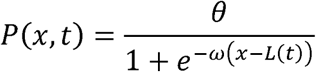

where *θ* is the maximum prevalence of CWD, and ω describes how quickly the prevalence attenuates with distance from the leading edge. Note that at *x* = *L*(*t*), that is, at the leading edge, the prevalence is *θ*/2. As the front advances toward Vermont, *L*(*t*) decreases and the prevalence of CWD at any given distance increases. If the density of deer on this landscape is given by 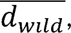 then the density of infected deer as a function of distance to the Vermont border is given by 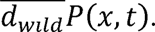

At any point in time, an individual deer may make a dispersal movement. If the dispersal distance is greater than the distance to the Vermont border, and the deer is infected with CWD, that would represent introduction of CWD into the State. If the dispersal kernel for white-tailed deer is given by *f*(*z*), where *z* is the dispersal distance and *f* represents the probability density function for dispersal, then the probability of a deer at a distance *x* from Vermont making the jump into Vermont is given by

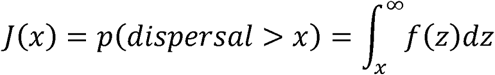

Putting this all together and integrating over the distance between the State and the initial leading edge, the expected number of infected wild animals entering the State in year t is given by

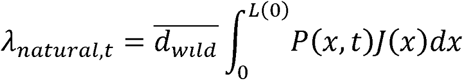

Finally, to represent the stochastic nature of these jumps, in any given year, we represent the number of infected deer that actually enter the State with a Poisson distribution:

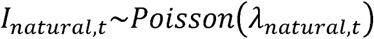

Where possible, the parameters were estimated from the published literature, and parametric uncertainty was included in the simulation. The current location of the leading edge of CWD infection, *L*(0), was estimated from 2024 surveillance data from Pennsylvania, with uncertainty expressed as a uniform distribution between 500 and 600 km. The rate of spread of CWD, *ρ*, was described with a lognormal distribution with mean 2.6 and standard deviation 0.25 (Hefley and others, 2017); this represents a mean spread of about 13.9 km/year. The maximum prevalence of CWD, *θ*, was described with uniform distribution between 0.4 and 0.5, based on observations of CWD prevalence in States where CWD has been established for a long time. The slope of the prevalence relationship as a function of distance, ω, was sampled from a uniform distribution between 0.01 and 0.02. The values were selected to approximate the declining density of disease across space, as one moves farther from a core area, while also conservatively assuming that there is a nonzero probability that CWD already exists in areas between the PA cases and the State. The average density of deer across this landscape, 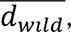 was represented as a uniform distribution between 5 and 25 deer/km^2^ (Walters and others, 2016). The dispersal kernel, *f*(*z*), was described with a log-normal distribution with mean 1.5 km and standard deviation 0.75 km (Long and others, 2015).

#### 8.2.4 Hunter-Harvest Carcass Transport and Disposal

The carcass introduction pathway was modeled as a random Poisson process whose rate was governed by a series of mutually exclusive and time-sensitive probabilistic events.

The rate of annual infections of wild deer in Vermont from CWD positive carcasses was given by

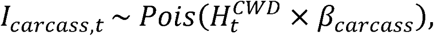

where,

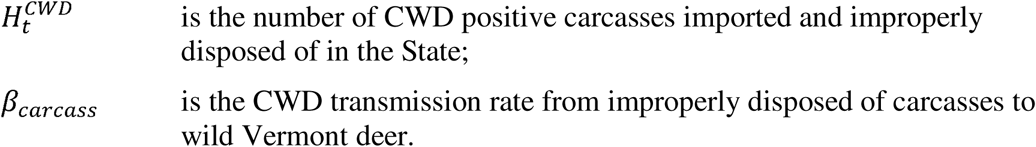

The number of CWD-positive carcasses deposited on the Vermont landscape was calculated as a function of the number of Vermont hunters that travel out of state for deer harvest opportunities, successfully harvest a deer that is CWD positive, transport it back to Vermont, and dispose of it improperly themselves, or following taxidermy or processing. The likelihood that a harvested deer was CWD positive is based on the spread model presented in Section 8.2.2, and specifically the average prevalence at time, *t*, for all spatial units in CT, MA, ME, NH, NJ, NY, PA, RI, and VT.

#### 8.2.5 Captive Cervid Importation

The last introduction pathway involves the importation of CWD-positive captive cervids to Vermont cervid farms and subsequent spread to wild deer via fenceline interactions. We assumed a Poisson distribution for the number of new animals imported annually and a binomial process governing whether imported animals were infected. We used historical importation data from 2020–2024 to calculate the average annual importation rate (2.87 animals imported per year) and assumed that the CWD prevalence in those imports was 2%, given VAAFM’s rigorous existing protocols.

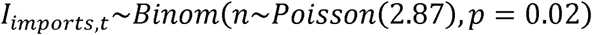

We assumed that all 9 cervid farms were identical, consisting of 100 animals on a 50-acre parcel, and these farms were individually tracked. Any infected imports each year were assigned to a single captive cervid farm. Following importation of a CWD positive animal, we assumed that CWD would subsequently spread within the farm and describe that process in section 8.4.2.

### 8.3 White-tailed Deer Population and CWD Dynamics in Vermont

#### 8.3.1 Overview

Along with the annual disease arrival processes described previously, we simulated deer population dynamics and disease (once it arrives) at annual time steps. The deer population and in-state CWD model was developed as a discrete time, multi-stage, SEI (susceptible-exposed-infected) model that simulated Vermont deer on an annual time step through a series of notable life-history events that occur throughout the biological year (June 1 – May 31) as well as disease transmission processes.

#### 8.3.2 Deer Population Model Structure

The deer population model included a total of 14 stages that included two sex classes (male and female), three age classes (fawns, juveniles, and adults), and three disease classes (susceptible, exposed, infectious) (fig. 3). We begin with a description of the transitions in the susceptible disease class (*S*). The first age class, fawns (*S_f_*), was assumed to enter the population with a 50-50 sex ratio at birth conditional on annual doe survival (*ϕ*) throughout the prior year. Then, fawns were assumed to survive their first year at a rate of 0.35 (natural survival without harvest), with different survival probabilities for neonates during summer (0.52) and first winter (0.68) based on data from VDFW (DOEPOP model, Vermont Department of Fish and Wildlife, written commun., 2024; fig. 3). We also assumed a small, additive rate of harvest of one percent (SD = 0.004) for fawns based on harvest data reported by VDFW.

**Figure 3.**
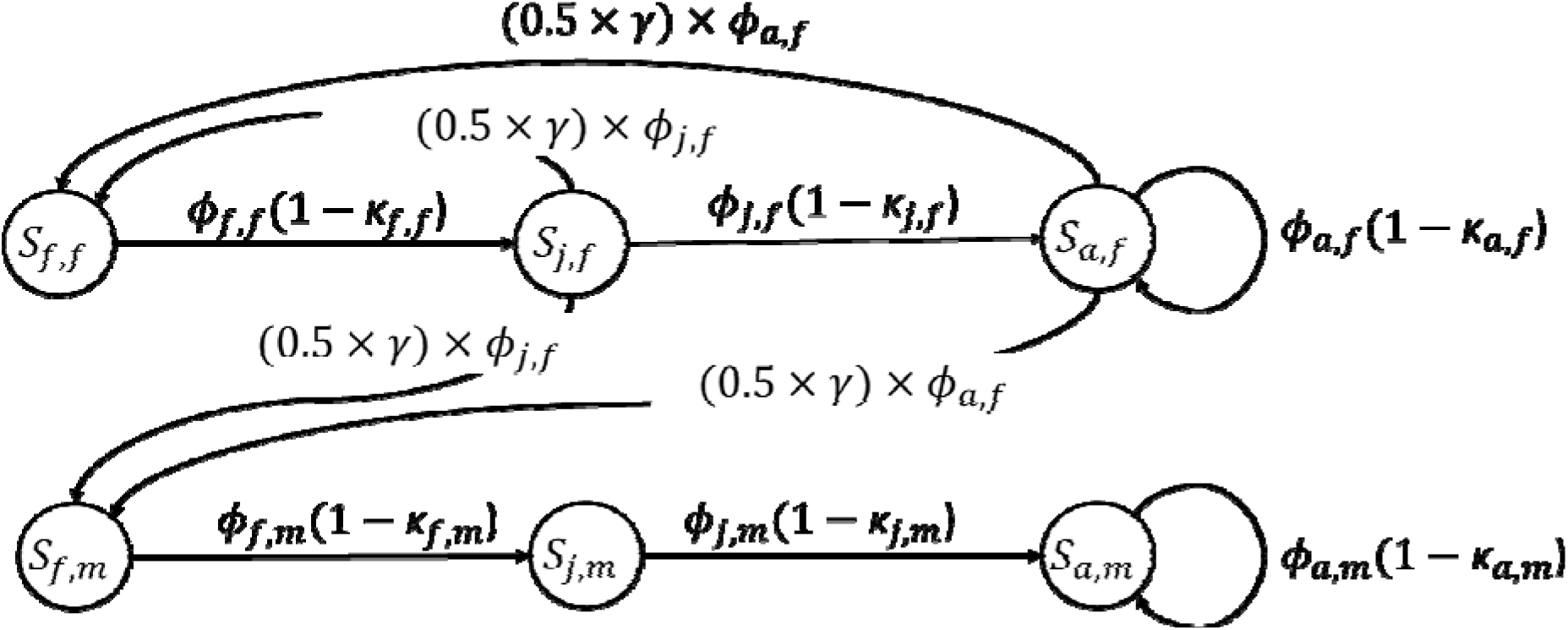
Life-history diagram for the disease-free (susceptible, S) white-tailed deer population dynamics model for two sex classes (male and female) and three age classes (fawns, juveniles, and adults).

Following the first year of life, deer then transitioned into juveniles of both sexes (fig. 3). Juvenile females then have the potential to reproduce, and in a disease-free population, suffered from natural and harvest mortality. Finally, juveniles transition into adults (2+ years old) and follow the same ordering of life-history events, including reproduction and natural and harvest mortality. Juveniles and adults can also enter exposed (*E*) and infectious (*I*) compartments, according to rates of transmission and disease progression. The rates can be found in table 2 and were taken from literature as well as VDFW modeled count and harvest data.

**Table 2.** White-tailed deer vital rate estimates used in the sex- and stage-structured population model. [SD, standard deviation; VDFW, Vermont Department of Fish and Wildlife; DOEPOP, doe-based deer population model]

| **Vital rate** | **Mean** | **Parametric SD** | **Source(s)** |
| --- | --- | --- | --- |
| Fawn survival | 0.35 | 0.04 | VDFW (DOEPOP) |
| Juvenile female survival | 0.87 | 0.02 | VDFW (DOEPOP) |
| Juvenile male survival | 0.87 | 0.02 | Assumed; VDFW (DOEPOP) |
| Adult female survival | 0.93 | 0.02 | Patterson and others, 2002 |
| Adult male survival | 0.93 | 0.03 | Assumed based on Patterson and others, 2002 |
| Reproduction rate | 1.60 | 0.01 | VDFW (DOEPOP) |
| Fawn harvest rate (both sexes)<sup>1</sup> | 0.01 | 0.004 | VDFW harvest data (2021–24) |
| Juvenile female harvest rate<sup>1</sup> | 0.10 | 0.03 | VDFW harvest data (2021–24) |
| Juvenile male harvest rate<sup>1</sup> | 0.40 | 0.12 | VDFW harvest data (2021–24) |
| Adult female harvest rate<sup>1</sup> | 0.05 | 0.02 | VDFW harvest data (2021–24) |
| Adult male harvest rate<sup>1</sup> | 0.15 | 0.03 | VDFW harvest data (2021–24) |

<sup>1</sup>Harvest rates are current ‘status quo’ conditions and subject to change under different management alternatives.

##### Density dependence

In addition to the baseline rates, we included a density-dependent regulatory process for reproduction based on an existing VDFW model and observed data. Thus, we consider a density-dependent reproductive rate, *γ*(*N*), for juvenile and adult females given by

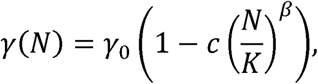

where

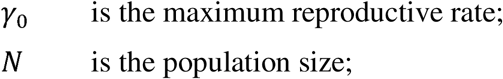

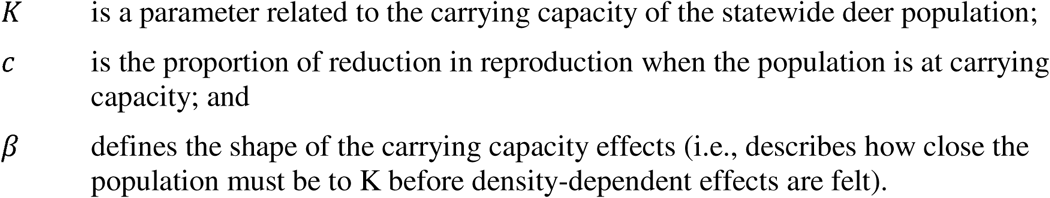

We described the proportional reduction parameter, *c*, as a uniform distribution with support between 0.5–0.9, and the shape parameter, *β*, as a uniform distribution with support between 3–4. The values for density dependence are similar to those considered in other Vermont wild deer population dynamics modeling efforts (DOEPOP, Vermont Department of Fish and Wildlife, written commun., 2024).

#### 8.3.3 CWD Dynamics in Wild Deer

Once the disease arrives in Vermont via one or more of the pathways described above, we modeled the intrastate disease growth process as an instantaneous rate, *λ*, composed of both direct and indirect transmission rates given by

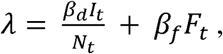

where

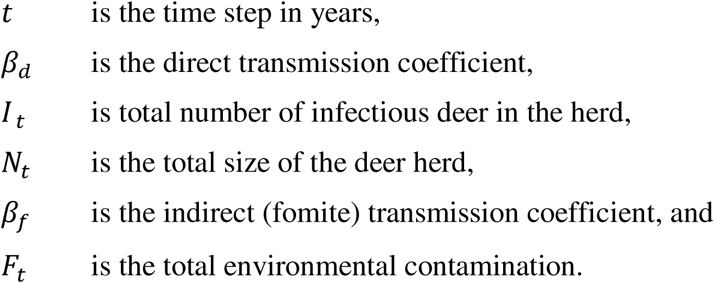

We then converted the instantaneous rate to a finite time (annual) infection probability for each susceptible individual in the population as 1 - exp (-*λ*). We calculated new exposed individuals at each time step using the per capita probability in a binomial random process where the number of trials equaled the number of susceptible animals. We modeled the state variable for environmental contamination, *F_t_*, as a function of live-animal shedding and prion decay given by,

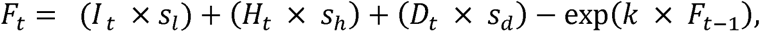

where

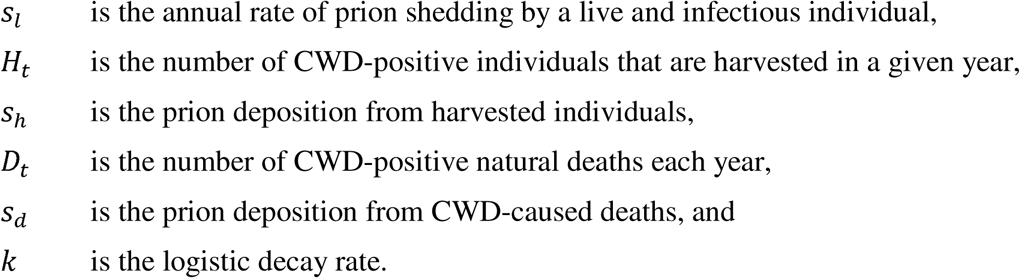

We assumed that all exposed individuals would transition into infectious individuals, if they survived the full year following exposure. The survival of infectious individuals with disease was assumed to be 25−50% lower based on study results from Wisconsin (https://dnr.wisconsin.gov/topic/research/projects/dpp/StudyResults; accessed June 5, 2026. Table 3 lists the required disease parameters, which were obtained either from empirical literature, expert elicitation, or exploration of the full range of support.

**Table 3.**
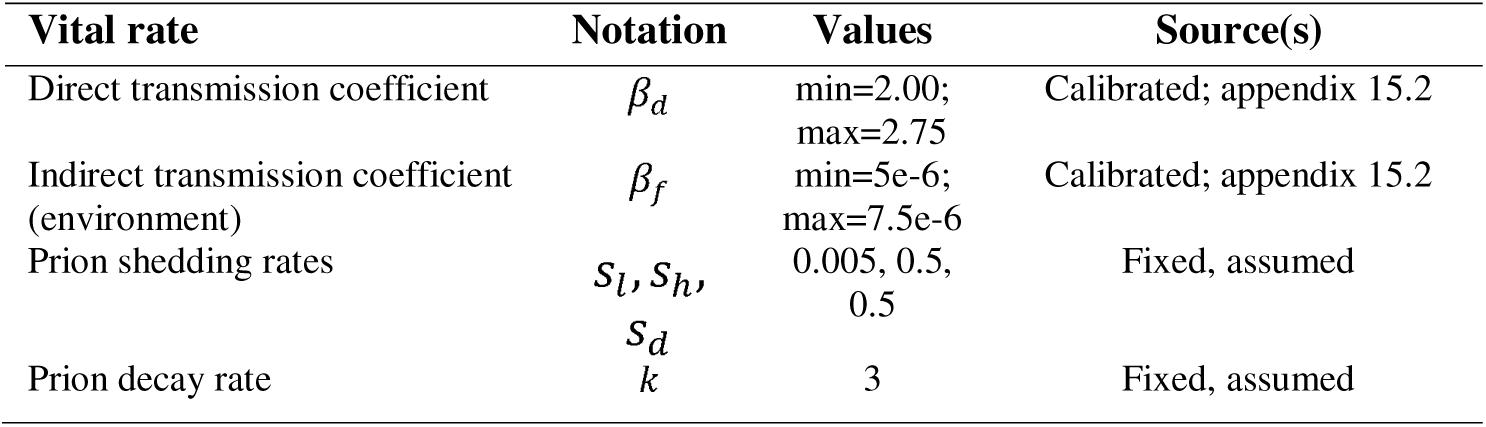
Estimates for parameters used to estimate chronic wasting disease transmission and decay in Vermont.

Beyond the spread of CWD within the wild population of Vermont deer, the disease could also spread across fencelines from the captive facilities.

### 8.4 Other Supporting Models

#### 8.4.1 Overview

Layered on top of the white-tailed deer population and disease models were submodels for CWD dynamics in captive deer and associated captive cervid farm years, forest regeneration potential, business revenues, surveillance, hunter participation, and agency costs (fig. 2).

#### 8.4.2 CWD Dynamics in Captive Deer and Farm Years

The model capturing the captive-wild interface informs the wild deer prevalence performance metric (PM 1b, table 1; through captive to wild fenceline transmission events) and the captive cervid farm years performance metric (PM 7, table 1; through both importation and wild to captive fenceline transmission events).

The number of wild deer that were newly infected each year via captive farms is a Poisson process governed by four quantities: 1) the density of infected captive cervids in infected deer per km^2^ (*d*_*captive.I,t*_), 2) the density of susceptive wild deer in susceptible deer per km^2^ (*d*_*wild.S,t*_), 3) a density scalar for the captive-wild contact rate (taken from Vercauteren and others, 2007; γ), and 4) the transmission efficiency of contact at fencelines (*β_fence_*).

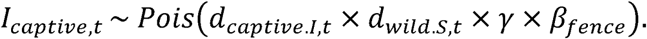

The statewide density of infected captive deer, *d*_*captive.I,t*_, builds on the cervid importation submodel described in 8.2.5. We assumed that disease prevalence in a farm would increase from its initial value after importation (*I_imports,t_*/100) to 0.25 in year 2, 0.50 in year 3, and that CWD would be detected by year 4. Infected captive deer density is calculated as the number of infected captive deer across all active facilities at time *t*, divided by the land area of the active captive facilities at time *t*. The statewide density of susceptible wild deer, *d_wild.S,t_*, is calculated using the deer population model and is the number of susceptible wild deer at time *t* divided by the total land area of Vermont.

The density scalar, *γ*, is computed from Vercauteren and others (2007), who report the only empirical captive-wild contact rate in the literature. We adjust that contact rate based on captive-wild densities in that study to generate *γ*, which, when coupled with wild susceptible and infected captive densities, yields the expected number of annual contacts between wild susceptible and infected captive deer annually. The successful transmission of CWD across the fenceline was assumed to be the product of the contact rate and transmission efficiency (*β_fence_*), which came from Mathiason and others (2009).

The metric for captive cervid farm years was calculated as the number of years of operation for each current cervid farm summed over all current cervid farms; thus, this metric is directly affected by farm depopulation and closure upon the detection of CWD. In the case of importation, we assumed that CWD would always be detected by the 4^th^ year post-introduction, and that farm depopulation would occur. In addition to importation, fenceline contacts with CWD-positive wild deer could also lead to CWD entering captive facilities and future depopulation.

For this second pathway, we again used a Poisson process governed by animal densities, a contact rate, and transmission efficiency. The infected wild deer and susceptible captive deer densities were calculated by dividing the count of infected or susceptible animals in that year by the land area of either Vermont or of active farms. We again used the density scalar for contact rate across the fence, as estimated from Vercauteren and others (2007), and transmission efficiency from Mathiason and others (2009). In the event of successful wild-captive transmission, we again allowed newly infected farms to increase in prevalence deterministically, as described above, with depopulation occurring in year 4. Upon depopulation, the remaining farm years associated with that farm were lost.

#### 8.4.3 Forest Regeneration

The performance metric associated with forest regeneration (PM 3, table 1) was calculated from total abundance in years 25 and 50 as estimated from the white-tailed deer population model. To inform the relationship between deer abundance and the regeneration potential in Vermont forests, we first calculated a statewide deer density at each time horizon. We then used a regression model specified in Boucher and others (2004) that estimated the total deciduous sapling density (saplings/hectare) as a function of deer density (deer per km^2^).

#### 8.4.4 Business Revenues

The performance metric that focused on business revenues (PM 6; table 1) was intended to capture any local economies that may be affected by CWD prevention and management actions. Most acute in those effects are the relationships between hunters and the expenditures that they make at small businesses associated with deer check stations, deer processors, and taxidermy establishments. Thus, business revenues were calculated using annual harvest estimates from the deer population model. For each yearly harvest estimate, we calculated the total annual revenue from businesses associated with hunting; 60% of the harvest is registered at check stations, which receive one dollar per deer, 50% of the harvest is sent to processors for approximately $125 dollars per deer, and 5% of the harvest is sent to taxidermists for approximately $800 dollars per deer (Vermont Department of Fish and Wildlife, written commun., 2024).

#### 8.4.5 Disease Surveillance

In general, agencies have available several potential CWD sampling opportunities, including active sampling by targeted culling (such as clinical-suspect animals), and passive (opportunistic) sampling that collects hunter-harvested, roadkill, or contaminated fomite samples. In our modeling environment, we considered both clinical-suspect animal testing and hunter-harvest testing as potential opportunities.

For hunter-harvested sampling, we assumed the State would take a random sample of harvested deer and test them for CWD. Because this was based on harvest, the sample would include higher numbers of age and sex classes that have higher rates of harvest (e.g., males). Thus, we assumed that the number of deer in each age, sex, and disease class could be given by a multivariate hypergeometric distribution

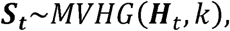

where ***S***_*t*_ is the vector of animals sampled by age, sex, and disease state, ***H***_*t*_ is the vector of all sex and age classes from the harvest that were available for testing, and k is the total sample taken. We simulated the sampling, then incorporated sampling diagnostic information for false positives and false negatives (0.98 and 0.99, respectively; National Academies of Science, Engineering, and Medicine, 2025), and finally, summed all individuals of the infected category to determine which animals tested positive.

The other sampling approach was focused on testing clinical-suspect animals. To approximate this type of testing, we preferentially selected animals from the population that were in advanced age classes (25% of the adult animals), as well as 50% of the infected disease class assuming that not all disease-positive animals would be showing clinical signs. We then drew a random sample from that subset of the population using the multivariate hypergeometric distribution as described previously.

In addition to the agency-directed sampling efforts, we also allowed hunters to submit their deer heads directly and modeled the testing rate as a function of apparent prevalence (fig. 4). This dynamic only set in following the initial detections of CWD and increased as disease trends tracked higher over time. In terms of the magnitude of voluntary testing, we considered strategies with increased outreach messaging about CWD to result in higher rates of sample submission compared to strategies that did not effectively communicate disease risks to public health.

**Figure 4.**
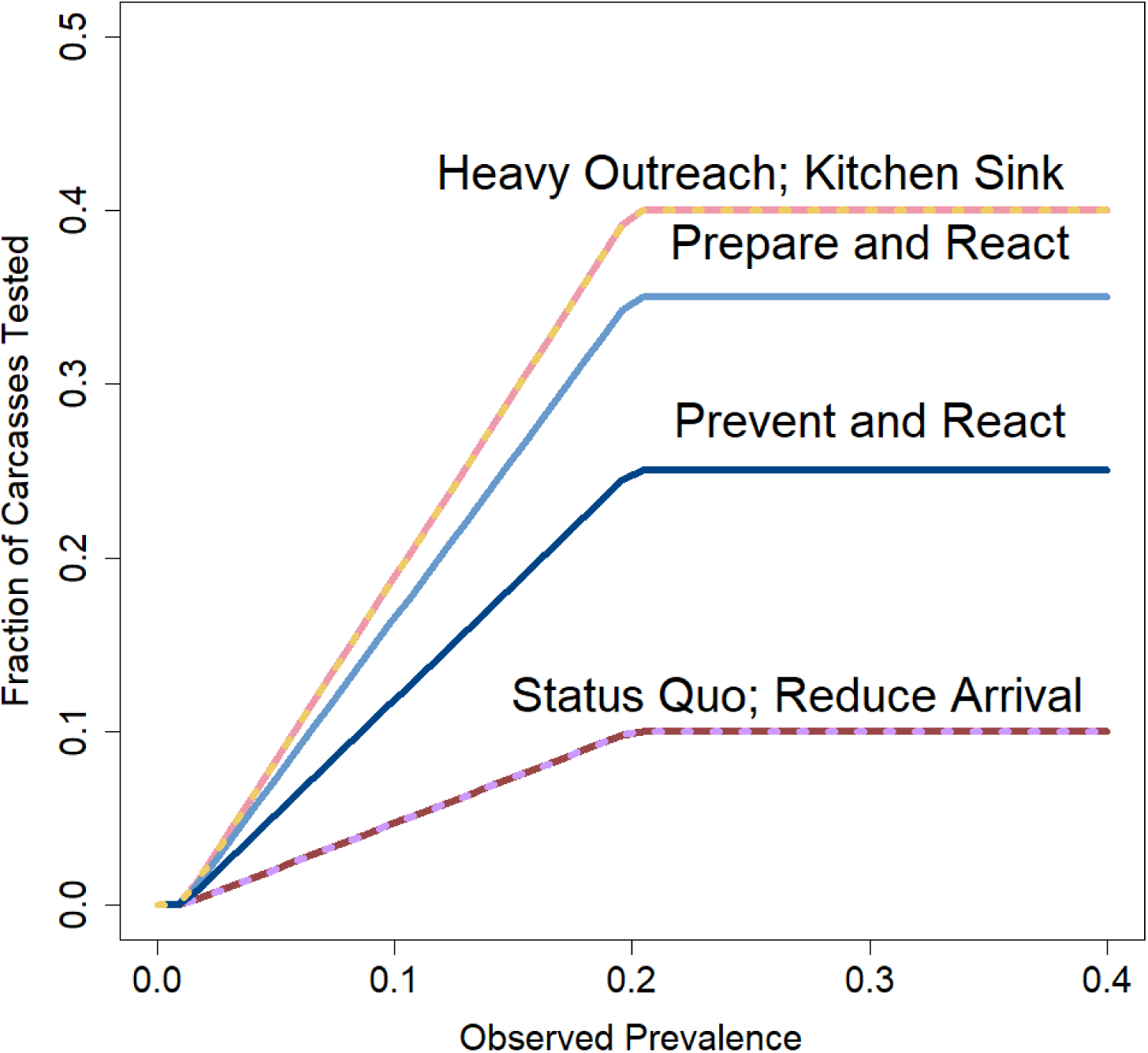
Fraction of hunters seeking voluntary testing of carcasses as a function of estimated prevalence of chronic wasting disease (CWD) in the Vermont deer population. Different CWD management strategies assumed different levels of outreach, resulting in different voluntary testing rates.

#### 8.4.6 Hunter Participation

In the near-term, we expect a decline in hunter participation over the next ten years at an annual loss of 0.75%, based on data from VDFW and an expectation that the >60-year-old hunter cohort would continue to age out of the hunting population, without corresponding recruitment of younger hunters. Following the ten-year period, we then assumed that annual hunter numbers would stabilize. These rates and periods are similar to dynamics observed in other states, such as Wisconsin (Mohr and others, 2025). Additionally, we allowed for an additive loss of hunters from the detection of CWD and the risk-based messaging from the agency. Despite both of these dynamics leading to declining hunter numbers, we assumed that there were hunters who would continue to go afield regardless of any changes to the deer abundance or regulations; we set this floor at 55,000 hunters, based on VDFW expectations.

#### 8.4.7 Agency Costs

The management of CWD is costly. For each action, we asked participants on the technical team to estimate expenses associated with management actions, accounting for both cost and frequency. We did not adjust for net present value or any expected inflationary changes to U.S. currency. Those estimates were included in the modeling work and summarized as a total across the time period (25- or 50-years). The breakdown for individual actions and the agencies responsible can be found in table 4.

**Table 4.** Cost breakdown for each management action that was considered in the decision analysis by agency. [VDFW; Vermont Department of Fish and Wildlife; VAAFM, Vermont Agency of Agriculture, Food and Markets]

| Action | Agency | Description | Cost<br>(frequency/<br>unit) |
| --- | --- | --- | --- |
| *Proactive costs* |  |  |  |
| Out-of-state hunter-harvested carcass regulations | VDFW | The cost associated with implementing restrictions on out-of-state carcass importation to limit the number entering the state | \$0 |
| Hunter-harvested carcass disposal guidelines | VDFW | The cost associated with implementing guidelines on in-state carcass disposal for hunters, taxidermists, and processors | \$40,000<br>(one time) |
| Preemptively close captive facilities | VAAFM | The cost associated with preemptively closing all captive cervid farms in the State | \$150,000<br>(one time) |
| Restrict captive cervid imports | VAAFM | The costs associated with restricting the number of captive cervids imported into the State | \$650<br>(per year) |
| Double fencing | VAAFM | The cost associated with preemptively constructing an additional fence around each captive cervid farm in the State | \$2,125,159<br>(one time) |
| Surveillance (pre-detection) | VDFW | The costs associated with testing deer for CWD | \$100<br>(per deer) |
| *Reactive costs* |  |  |  |
| Sharpshooting | VDFW | The costs associated with sharpshooting deer in areas where CWD-infected deer are detected | \$180<br>(per deer) |
| Surveillance (post-detection) | VDFW | The costs associated with testing deer for CWD, including hunter-requested testing | \$100<br>(per deer) |
| Close infected cervid farms | VAAFM | The cost associated with closing a captive cervid farm when CWD is detected in the herd | \$750,000<br>(per farm) |

### 8.5 Effects of Management Actions

#### 8.5.1 Actions Taken at Captive Deer Farms

The actions at captive deer facilities were assumed to affect two primary dynamics, the importation rate of infectious and healthy individuals through increasingly stringent standards on the permitting of cervid importation, and the fenceline contact rate between captive and wild deer being affected by actions to double-fence facilities. We assumed that stringent importation standards reduced the import rate by 75% (Vermont Agency of Agriculture, Food and Markets, written commun., 2024) whereas double fencing reduced contact rates by 67% (Schultze and others, 2023). Finally, the closure of facilities effectively stopped any risk of captive-wild CWD transmission in both directions.

#### 8.5.2 Proactive Regulations

We assumed that proactive actions, including a total ban on importation of deer parts from all States and increased messaging about CWD risks, would reduce the rate of carcass import and improper disposal by 25% each, at maximum (i.e., 25% reduction in importation of high-risk parts and 25% reduction in improper disposal). As for the out-of-state deer that go to taxidermists and processors, we assumed a reduction of 50% in improper disposal given the ability of agencies to develop more targeted messaging and outreach to these groups.

#### 8.5.3 Outreach and Education

We considered a broad range of outreach and communication efforts that aligned with the general approach that each alternative took toward CWD and the risks presented (appendix 15-1). In some cases, like those discussed in the section above, outreach and education efforts were considered to enhance the ability of an action to be successful. This was especially relevant to outreach associated with import restrictions and carcass disposal, hunter participation in CWD sampling efforts, and the continued engagement of hunters when CWD prevalence reached higher levels (fig. 4). Additionally, while we did not consider a specific quantitative effect for many of the actions listed in Appendix 15.1, there is widespread acknowledgment that outreach and education is a critical component to all the alternatives we evaluated.

#### 8.5.4 Targeted Removal of Deer

One of the potential reactive management actions is targeted removal of deer in an area surrounding detection of one or more CWD-positive individuals, through incentivized harvest, sharpshooting, or other methods of removal. To model the efficacy of this action as a function of the desired sharpshooting effort and the statewide CWD prevalence at first detection, we assumed a simplified case at the stage of early arrival to the State. We assume that all the infected deer within the State are within a single cluster (dark teal circle with radius *r*_1_ in fig. 5), that a single detection has occurred (which is, of course, located within the infected cluster), and that the sharpshooting (or other removal) effort is concentrated in a circle surrounding the detection (orange circle with radius *r*_2_ in fig. 5). If the sharpshooting effort is too small (*r*_2_ << *r*_1_), the orange circle will be entirely within the teal circle and only a small fraction of the infected population will be removed. If sharpshooting effort is very large (*r*_2_ >> *r*_1_), then the orange circle will entirely encompass the teal circle, all of the infected animals are at least subject to the removal effort, and the fraction removed will depend on the proportion of deer removed within the removal radius. In the intermediate case (as depicted in fig. 5), the efficacy of the effort depends on the degree of overlap between the two circles.

**Figure 5.**
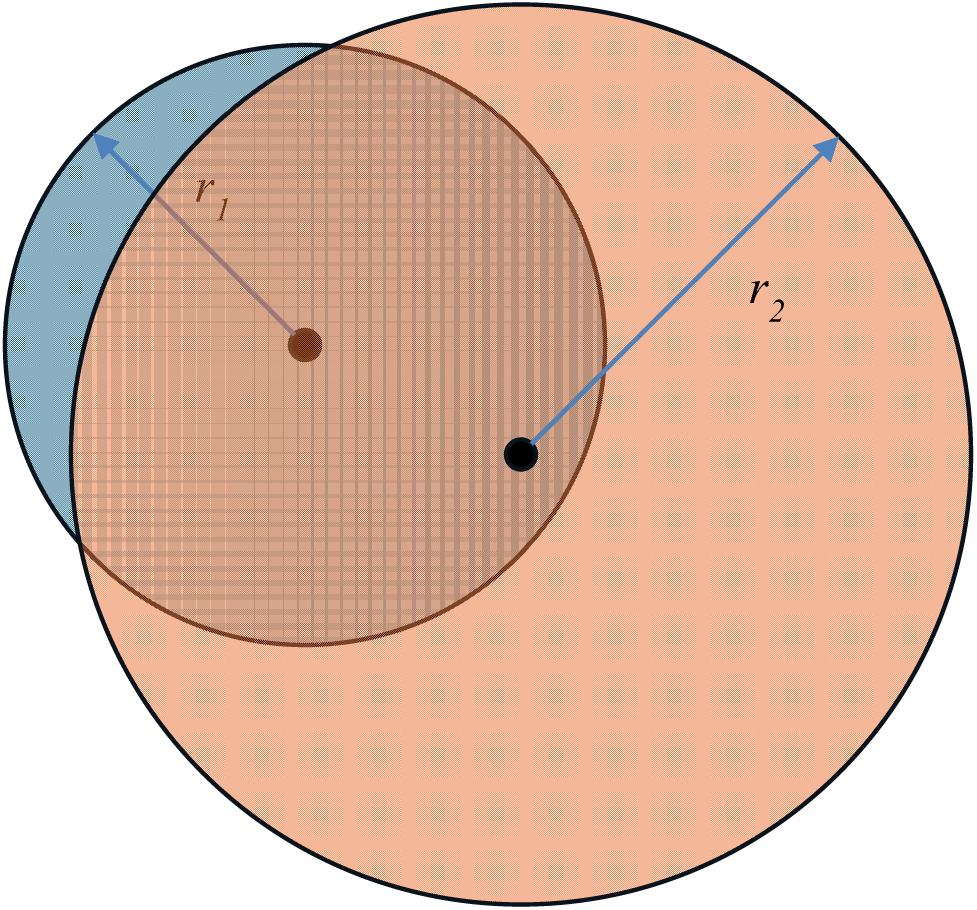
Overlapping concentrations of deer infected with chronic wasting disease (dark teal) and sharpshooting effort (orange). The overlapping area is the portion of the infected population that is subject to sharpshooting.

To estimate the radius of the infected cluster (*r*_1_), we assumed that all the infected deer in the State are within a single cluster, the prevalence within the infected cluster is *p*_max_ = 40%, and the deer density is *d* = 10/mi^2.^ Thus, the radius of the infected cluster is given by

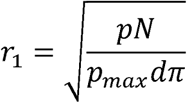

where *p* is the statewide prevalence of CWD, and *N* is the statewide deer population size (hence, *pN* is the number of infected deer statewide).

To estimate the radius of the targeted removal effort (*r*_2_), we assumed the removal area was designed so that a removal rate of *f* (default, *f* = ¾) is expected to achieve the desired number of removals (*S*). Thus, the radius of the targeted removal effort is given by

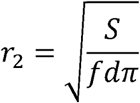

where *S*, *f*, and *d* are as defined above.

To model the effect of targeted removal, we derived the expected fraction of the infected cluster that is removed, accounting for the number of infected animals, deer density, removal effort, and removal efficiency (refer to details of the derivation in Appendix 15.3). The fraction of the infected cluster that is subject to sharpshooting (or other targeted removal), that is, the fraction of the teal circle in figure 5 that overlaps with the orange circle, decreases sharply with increases in the statewide prevalence of CWD (fig. 6), because as the prevalence increases, so does the size of the infected cluster. Thus, targeted removal can be effective, but only if CWD is detected early, when the statewide prevalence is very low (<0.005).

**Figure 6.**
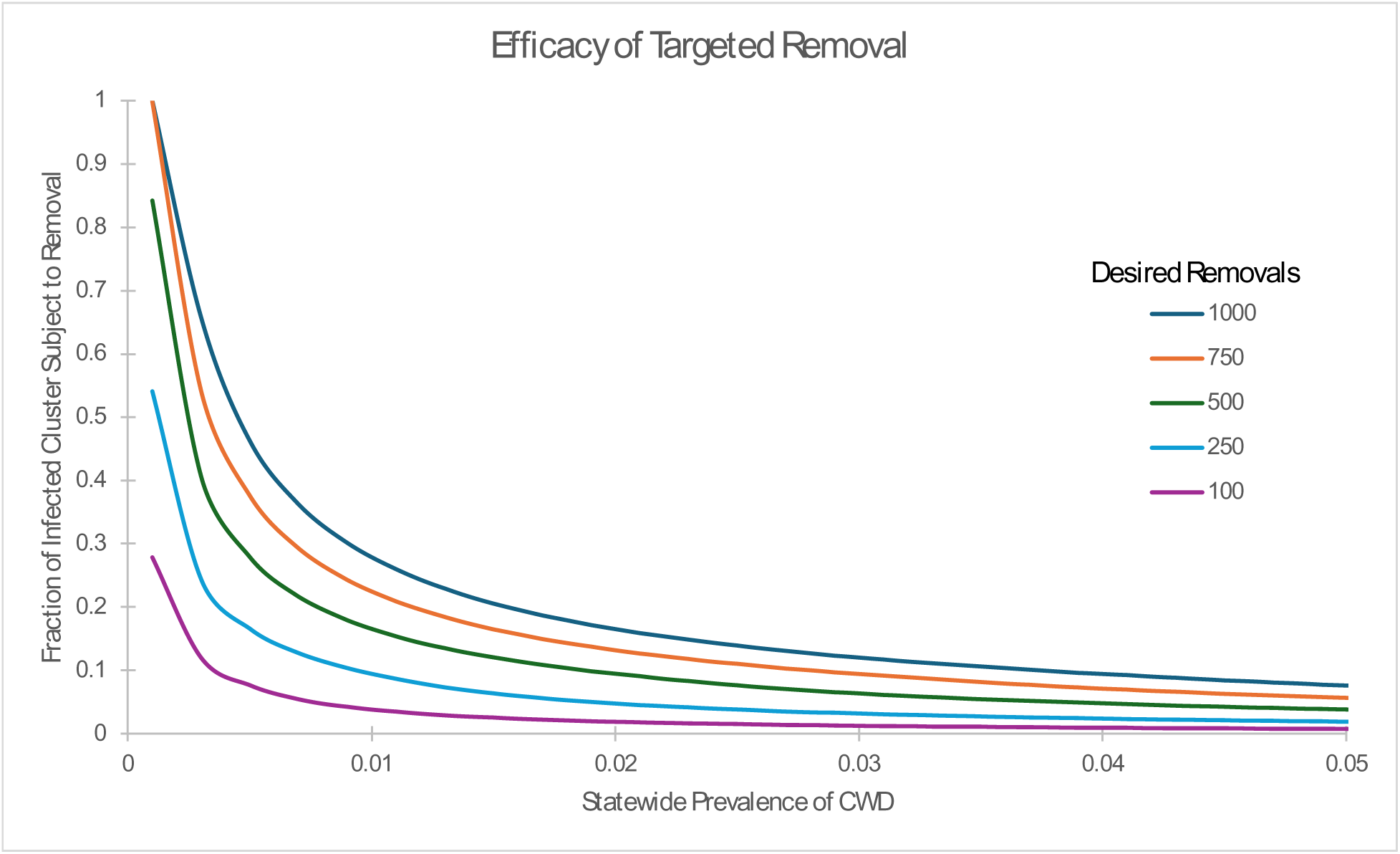
Fraction of infected cluster subjected to targeted removal as a function of the statewide prevalence of chronic wasting disease (CWD) in deer, for five different levels of desired removal.

#### 8.5.5 Increased Hunting Pressure

We assumed that hunting pressure could be increased by 10–20% under more liberal harvest policies, only in scenarios following the new detection of CWD. We considered this increase to apply evenly among all age and sex classes and to be based on existing harvest rates between 2020–2024. The increase was also layered on top of the approximated losses of older generation hunters, and those that stop hunting because they are risk averse to CWD.

## 9 Phase I – Consequence Analysis Results

We first analyzed the set of six alternatives that were designed, in part, to understand the relative roles that the proactive, reactive, and surveillance actions had in achieving the performance metrics. Because CWD arrival, detection, and prevalence drive many of the outcomes of the six alternatives, we present these results first. We found varying performance among the eight metrics across the alternatives, in terms of the arrival, spread, and long-term consequences of CWD and its associated effects.

### 9.1 Chronic Wasting Disease Dynamics

#### 9.1.1 Arrival and Detection of CWD in Vermont

Before exploring the outcomes of interest, it is valuable to understand two intermediate processes that affect the results—the time-to-arrival and time-to-detection of CWD in Vermont. Under the *Status Quo* strategy, the mean time-to-arrival of CWD in wild deer, from all pathways combined, is 7.7 years (range:1–26 years), with 53% of the first arrivals coming by wild deer dispersal across the landscape, 17% of the first arrivals coming from disposal of infected carcasses or parts on the landscape, and 30% of first arrivals coming from captive cervid farms. Under the *Heavy Outreach* strategy, the mean time-to-arrival is similar to *Status Quo* at 7.7 years (range:1–29 years). Under the *Reduce Arrival* strategy, the mean time-to-arrival increases to 9.5 years (range:1–29 years). The *Prepare & React* strategy does not have any proactive interactions, so the arrival time is about the same as for *Status Quo* (7.7 years, range:1–28 years). The *Prevent & React* strategy has some of the same proactive actions as *Reduce Arrival*, and a similar expected arrival time (9.5 years, range:1–30 years). Finally, the *Kitchen Sink* strategy has the longest expected arrival time, at 10.6 years (range:1–29 years). The differences in arrival time across strategies are small because there are not proactive actions that affect the dominant pathway of arrival (deer dispersing across the landscape).

Considering each pathway individually, introduction from natural wild deer movements had the soonest mean time-to-arrival estimate at 13.2 years, on average (SD=7.6, fig. 7). Following closely behind are introductions from carcass transport at 14.7 years (SD=11.5 years). Actions intended to delay the arrival from carcass transport, importation restrictions and guidelines for disposal, extended this average by 5.3 years to 20.0 (SD=14.2). Finally, status quo captive cervid import was estimated to result in CWD arrival in wild deer in 26.5 years, on average (SD=23.9). If captive cervid imports were to be reduced and double fencing installed, the average extends to 76.6 years (SD=65.4), with CWD never spreading to wild deer from captives in 14.7% of simulations.

**Figure 7.**
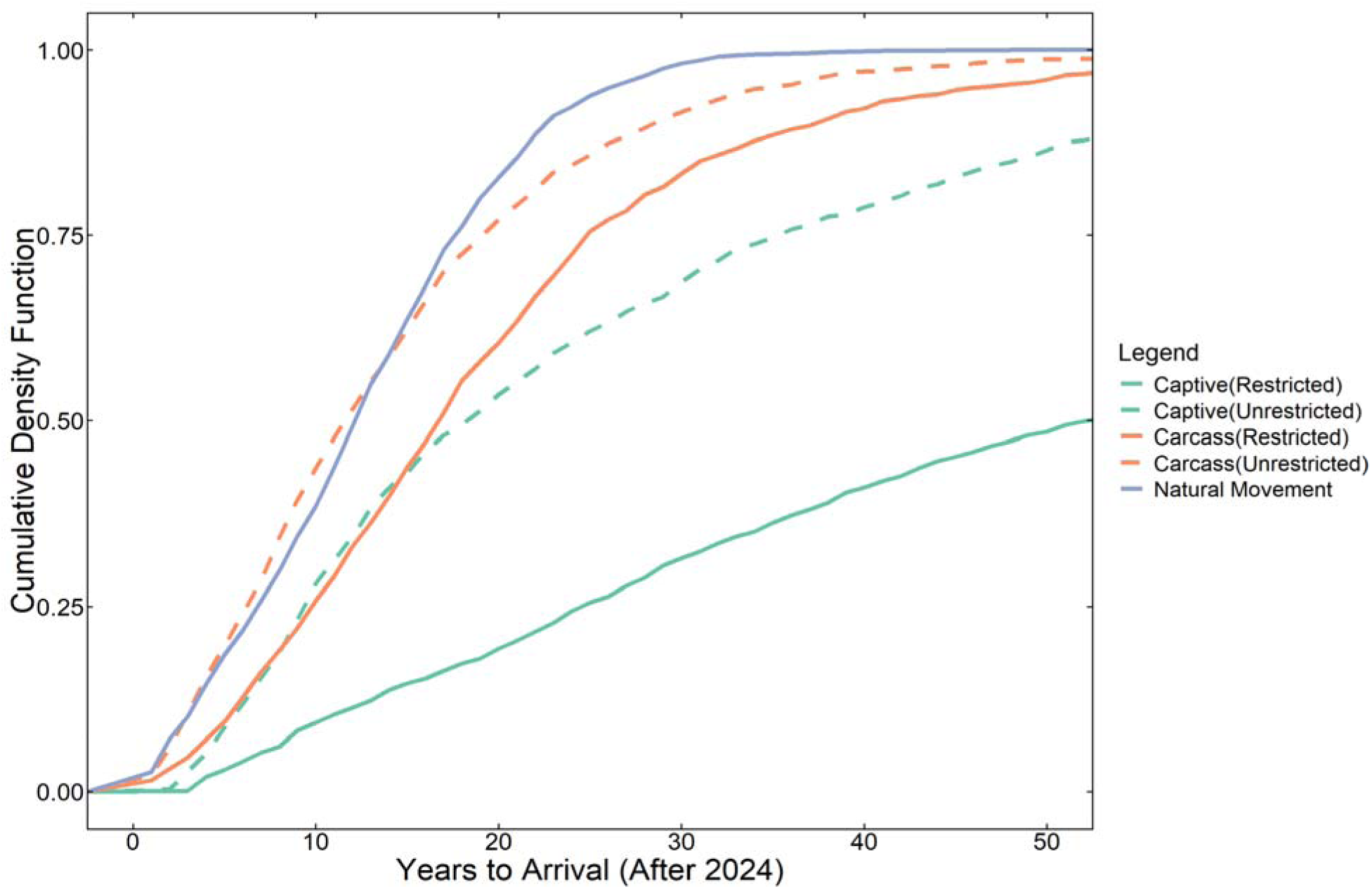
Empirical cumulative density functions for five different pathway scenarios for the arrival of chronic wasting disease (CWD) in Vermont. The first, natural movements of deer, results in earlier arrivals in Vermont wild deer, followed by carcass transport, and captive cervid imports. Carcass import restrictions and guidelines for carcass disposal extend the time-to-arrival from carcass imports, whereas captive cervid import restrictions and double fencing of facilities extends time-to-arrival from captives. Under captive cervid import restrictions and double fencing, CWD never spread to wild deer from captives in 14.7% of simulations in 200 years.

The mean time between arrival and detection of CWD in Vermont is directly affected by the surveillance strategy, with the *Status Quo* surveillance strategy showing a 29.7 year lag between arrival and detection; the moderate surveillance strategy (used in *Reduce Arrival* and *Prevent & React*) showing a 21.0 to 22.4 year lag between arrival and detection; and the maximal surveillance strategy (used by *Heavy Outreach*, *Prepare & React*, and *Kitchen Sink*) showing a 15.1 to 16.6 year lag.

Combining these two processes, the expected time from now until first detection of CWD in the state is longest under the *Status Quo* (37.4 years), shortest under *Heavy Outreach* and *Prepare and React* (22.8 to 23 years), and between 27.2 and 31.9 years in the other strategies.

#### 9.1.2 CWD Dynamics in Wild White-Tailed Deer

Following the introduction of CWD into the Vermont deer herd, the disease grows in the wild population as described previously. The growth patterns were summarized for the second performance metric, mean prevalence of CWD in the statewide deer population at 25 and 50 years from present. The differences in mean prevalence across alternatives are relatively small at year 25 (0.97% under *Status Quo* and 0.08% under *Kitchen Sink*) but become more pronounced at year 50 (23.6% under *Status Quo* and 4.0% under *Kitchen Sink*). The other alternatives fell within this range, with *Heavy Outreach* and *Reduce Arrival* outcomes being closer to status quo, and the reactive strategies, *Prepare and React*, and *Prevent and React* having lower prevalence.

### 9.2 Wild Deer Population Dynamics

The second performance metric associated with the health of the deer herd tracked the mean size of the deer population at 25 and 50 years. Initially, all the strategies show an increase in the deer population, as the older generation ages out of the hunter population and harvest rates decrease. Eventually, all strategies result in declines in the deer population, but the strategies differ in timing and cause of declines. There are three primary factors that affect the size of the deer population over time: the prevalence of CWD, which eventually depresses the population growth rate; the harvest rates achieved through hunting regulations; and the number of active deer hunters. The patterns in performance across alternative strategies are different at 25 and 50 years, because the effect of CWD on the deer population is largely not felt in the first 25 years and therefore the population projections under all six alternatives remain comparable to the current deer population size (∼146,000 deer).

At the 50-year mark, the mean deer populations start to diverge among the six alternatives, with those strategies that had higher CWD prevalence also having fewer deer (table 5). The 50-year population is smallest under the *Heavy Outreach* strategy (121,516), and largest under the *Kitchen Sink* strategy (140,841). In between those two strategies, the strategies with more aggressive reactive CWD measures had larger projected populations (e.g., *Prepare and React)* compared to those that had limited reaction to CWD detection (e.g., *Status quo*) (table 5).

**Table 5.** Consequence table showing the expected performance over 50 years of six alternative strategies against eight performance metrics for the management of chronic wasting disease (CWD) in Vermont white-tailed deer. In each row, the alternative that performs best is shaded green, the alternative that performs worst is shaded pink, and strategies that perform better than the status quo (as of 2024) are shaded yellow. Strategies that perform worse than the status quo are unshaded. [#, number; $, U.S. dollar; &, and; ha, hectare; yr, year]

| **Performance Metric** | **desired direction** | **Status Quo** | **Heavy Outreach** | **Reduce Arrival** | **Prepare & React** | **Prevent & React** | **Kitchen Sink** |
| --- | --- | --- | --- | --- | --- | --- | --- |
| Size of deer population at year 50 | Max | 123,400 | 121,516 | 127,308 | 136,010 | 125,968 | 140,841 |
| CWD prevalence at year 50 | Min | 0.24 | 0.21 | 0.21 | 0.10 | 0.19 | 0.04 |
| Mean annual active hunters over 50 years | Max | 61,029 | 59,132 | 60,428 | 59,105 | 60,232 | 59,716 |
| Sapling density at year 50 (saplings/ha) | Max | 2,907 | 2,913 | 2,895 | 2,867 | 2,899 | 2,851 |
| Mean total cost to agencies (\$ over 50 yr) | Min | 5,483,964 | 10,194,222 | 6,154,641 | 7,212,745 | 7,665,994 | 4,587,311 |
| Total CWD-positive animals consumed (#) | Min | 28,727 | 21,545 | 22,971 | 9,725 | 20,387 | 3,153 |
| Net annual revenue for associated business (\$) | Max | 1,692,209 | 1,762,274 | 1,705,280 | 1,764,873 | 1,762,104 | 1,801,300 |
| Cervid farm operations (total farm-years) | Max | 358 | 359 | 392 | 380 | 391 | 0 |

### 9.3 Hunting Participation and Human Consumption of CWD-infected Meat

The third performance metric tracks the mean number of active hunters over 25 or 50 years, where the active hunters are people who both have a license and participate in the hunting season. Over time, two factors affect the number of active hunters: the aging of the Baby Boomer generation out of the active hunting population; and a reduction in participation after detection of CWD in the state. There is a noticeable loss of hunters predicted over the next 10 years, similar for all strategies. The differences across the alternative strategies are small over both 25 and 50 years. Over the first 25 years, the average number of active hunters is lowest for the *Heavy Outreach, Prepare and React,* and *Kitchen Sink* strategies, because they have the earliest detections of CWD (owing to both an earlier arrival and a higher effort at surveillance) and thus, the earliest decrease in hunter participation. The best performing strategy at 25 years is the *Status Quo* strategy because moderate surveillance effort results in a longer time-to-detection for CWD and therefore hunters have less time to react to it. There are fairly similar outcomes at the 50-year time horizon.

### 9.4 Forest Regeneration

The fundamental objective associated with forest health has many dimensions, but through discussion with VFPR and USFS, the focus of this analysis was the aspect of forest health most affected by deer density—regrowth. Based on data from a study in Quebec, deciduous sapling density was assumed to vary linearly with the deer density. The sapling density at years 25 and 50 showed the opposite pattern across alternatives as deer density, those alternatives with more deer had fewer saplings compared to those with less deer, albeit the range was narrow (25-year: 2,831−2835 and 50-year: 2851−2912 saplings/ha).

### 9.5 Costs for Agencies

The costs associated with CWD management vary by agency and strategy; the performance metric shows the mean annual costs for VDFW, VAAFM, VDH, and VFPR over 25 or 50 years. For VDFW, costs arise from surveillance, voluntary testing by hunters, sharpshooting, and various outreach-related actions. The cost of VDFW outreach actions ranges from $0 per year (*Status Quo*) to upwards of $220,000 per year (*Kitchen Sink*), with all strategies other than *Status Quo* and *Prepare* & *React* including at least one new full-time employee (annual cost of $156,000) in both the proactive and reactive time periods. For VAAFM, costs (beyond what is associated with *Status Quo*) arise from closing and decontaminating cervid facilities and from proactive double-fencing. Proactive double-fencing of all 9 facilities was considered in the *Reduce Arrival* and *Prevent & React* strategies at a one-time cost of $2.12M (for all 9 facilities); reactive depopulation of herds once CWD was detected in them was considered in all strategies, at a cost of $750,000 (per herd). The *Kitchen Sink* strategy proactively depopulates and closes all facilities, at a total one-time cost of $150,000. The costs associated with depopulation prior to CWD detection are much lower for VAAFM because producers can bring to market uninfected animals, whereas after a CWD detection the value of animals is lost and indemnity payments to producers instead may come from the State. For VDH, costs are associated with the intensity of outreach efforts and range from $7,800 annually for reactive messaging (*Status Quo* and *Reduce Arrival*) to the development of a database to track potential human exposures ($85,000 one-time cost in *Heavy Outreach, Prevent & React, Prepare & React,* and *Kitchen Sink*). Costs for VFPR individual actions range from $1,600−3,200 per year with the *Kitchen Sink* being most costly (at $6,400 prior to and $14,400 after first detection) and the *Status Quo* being least costly (at $0).

### 9.6 Business Revenues

The seventh performance metric reflects the effects of CWD and its management on businesses associated with deer hunting, notably, deer meat processors, taxidermists, and local convenience stores that commonly host deer check stations. The alternative strategies differ because of differences in the size of the deer harvest and the regulations placed on processors and taxidermists. The differences across alternatives, however, are relatively small (the range between the highest and lowest annual revenue is about $109,000 compared to the *Status Quo* revenue of about $1.69M per year over 50 years).

### 9.7 Captive Cervid Operations

The final performance metric summarizes the number of farm-years (the sum of the duration of time that each of the nine farms stays operational) across strategies, accounting for farms that need to be depopulated either because they imported an infected animal, they were infected by a wild animal, or they were proactively closed. The maximum possible value for farm-years is 225 for the 25-year consequence table and 450 for the 50-year consequence table (table 5), both of which would occur if all nine farms remain operational for the entire time period. The patterns are similar across strategies for both time horizons. The *Kitchen Sink* resulted in 0 farm-years owing to proactive depopulation of all cervid farms whereas *Reduce Arrival* and *Prepare & React* had the largest values (392 and 391 farm-years after 50 years, respectively). These patterns were apparently due to reduced risk of infection from import because of additional import restrictions and reduced risk of infection from wild animals because of double-fencing. All other strategies resulted in intermediate values for this performance metric.

### 9.8 Full Consequence Table and MCDA

The overall tradeoffs among strategies suggested that those strategies with large monetary and staff-time investments in surveillance and reactive actions, notably the *Prepare and React* and *Kitchen Sink* strategies, provided large decreases in the overall prevalence of CWD compared to those strategies that were less intensive (e.g., *Status Quo*) and focused more on preventative measures (e.g., *Reduce Arrival, Prevent and React*). These stark differences among strategies made the deliberation among alternatives difficult for the agencies involved. This contrasting performance, while expected based on the design of the alternatives, nevertheless presented challenges to understanding how important the variation among the six strategies was, and which if any, performance metrics would guide the selection of a viable alternative. To aid this deliberation, we conducted an MCDA approach whereby we elicited estimates of the relative importance of each performance metric using swing weighting (von Winterfeldt and Edwards, 1986).

The swing weighting exercise asked policymakers to use the values in the 50-year consequence table and to consider future outcomes where all but one performance metric could be maximized. We asked the participants to first rank and then score each outcome from most to least preferred. The CWD prevalence in wild deer in 50 years consistently had the highest weight (range across participants: 15-36%) among the performance metrics (table 6). As a result, those alternatives that had the lowest prevalence, *Prepare and React*, and *Kitchen Sink*, consistently outperformed alternatives with higher 50-year prevalence outcomes (fig. 8). Despite these consistent rankings, the first phase of the consequence analysis concluded with a consensus that the strategy needed to be refined with respect to costs and disease outcomes; additionally, the group wanted further clarity about the role that captive cervid importation had on 50-year outcomes. As a result, the set of alternatives was expanded and additional MCDA tools were brought to bear on the deliberation of alternative CWD management strategies.

**Figure 8.**
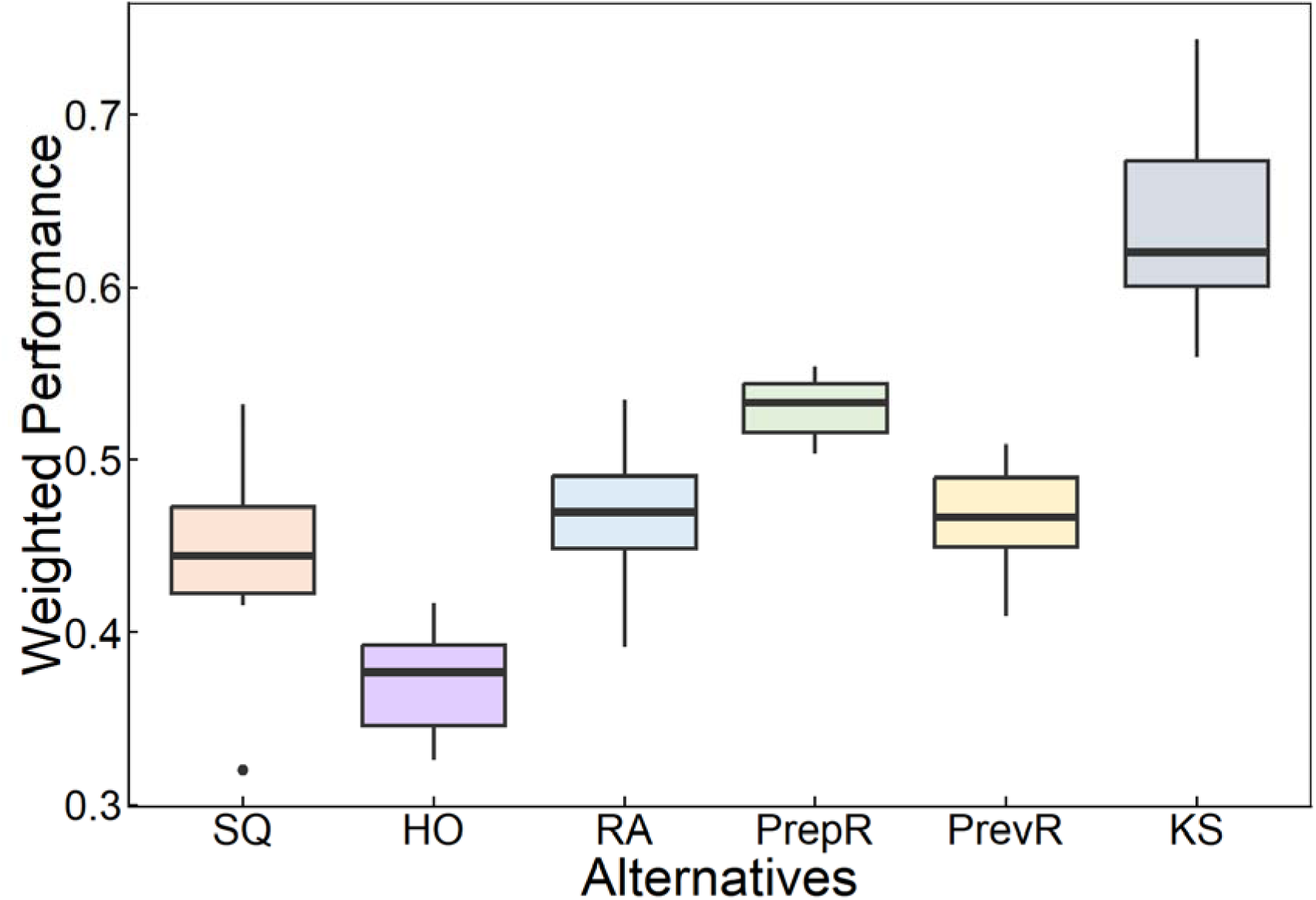
Weighted performance of the six alternatives for the decision analysis on chronic wasting disease management in deer in Vermont considering values provided by fourteen workshop participants from the Vermont Department of Fish and Wildlife and the Vermont Agency of Agriculture, Food and Markets and mean values for each of the eight performance metrics. The Kitchen Sink was consistently the best performing alternative regardless of which policymaker was used. Box plots depict the minimum, first quartile, median, third quartile, and maximum, with outliers depicted as single points. [SQ, Status Quo; HO, Heavy Outreach; RA, Reduce Arrival; PrepR, Prepare and React; PrevR, Prevent and React; KS, Kitchen Sink].

**Table 6.** Table showing the relative weights placed on each of the eight-performance metrics for the decision analysis on chronic wasting disease management in Vermont. The highest weight was placed on chronic wasting disease prevalence for 13 out of the 14 participants (staff from the Vermont Department of Fish and Wildlife, Vermont Agency of Agriculture, Food and Markets, Vermont Department of Health, Vermont Department of Forests, Parks and Recreation, U.S. Fish and Wildlife Service, U.S. Department of Agriculture Animal Plant and Health Inspection Service)

| Performance metric number | Performance metric | Participant Number |  |  |  |  |  |  |  |  |  |  |  |  |  |
| --- | --- | --- | --- | --- | --- | --- | --- | --- | --- | --- | --- | --- | --- | --- | --- |
|  |  | 1 | 2 | 3 | 4 | 5 | 6 | 7 | 8 | 9 | 10 | 11 | 12 | 13 | 14 |
| 1a | Size of deer population in 25 and 50 years | 0.14 | 0.08 | 0.09 | 0.05 | 0.14 | 0.13 | 0.02 | 0.12 | 0.10 | 0.08 | 0.17 | 0.13 | 0.13 | 0.15 |
| 1b | CWD prevalence in 25 and 50 years | 0.20 | 0.17 | 0.36 | 0.18 | 0.20 | 0.15 | 0.22 | 0.16 | 0.20 | 0.27 | 0.22 | 0.17 | 0.17 | 0.19 |
| 2 | Average annual number of active hunters over 25 and 50 years | 0.16 | 0.13 | 0.09 | 0.17 | 0.16 | 0.13 | 0.13 | 0.13 | 0.12 | 0.05 | 0.13 | 0.07 | 0.09 | 0.15 |
| 3 | Density of deciduous saplings at year 25 and 50 (saplings/hectare) | 0.12 | 0.10 | 0.11 | 0.17 | 0.19 | 0.11 | 0.09 | 0.11 | 0.10 | 0.14 | 0.11 | 0.14 | 0.09 | 0.09 |
| 4 | The average cost per year over 25 and 50 years | 0.06 | 0.16 | 0.09 | 0.14 | 0.15 | 0.11 | 0.14 | 0.13 | 0.16 | 0.25 | 0.11 | 0.16 | 0.15 | 0.15 |
| 5 | The total number of CWD positive deer harvested over 25 and 50 years | 0.18 | 0.15 | 0.22 | 0.14 | 0.02 | 0.14 | 0.20 | 0.15 | 0.18 | 0.14 | 0.17 | 0.16 | 0.19 | 0.17 |
| 6 | Average annual net revenue for hunting-related businesses over 25 and 50 years | 0.08 | 0.12 | 0.02 | 0.13 | 0.10 | 0.13 | 0.10 | 0.10 | 0.12 | 0.05 | 0.07 | 0.12 | 0.17 | 0.11 |
| 7 | Cumulative captive cervid farm-years over 25 and 50 years | 0.04 | 0.08 | 0.02 | 0.02 | 0.06 | 0.11 | 0.11 | 0.10 | 0.02 | 0.01 | 0.02 | 0.05 | 0.02 | 0.00 |

## 10 Phase II – Scenario Results

The emerging results during Phase I highlighted the important tradeoffs among surveillance, proactive, and reactive elements of a CWD management plan. However, based on the consequences across strategies and the relative weights placed on the performance metrics, not all metrics were deemed critical to distinguishing the right management strategy. Thus, the next phase of the analysis (Phase II) focused on a more refined set of central questions. Specifically, the multi-agency decision-making team was interested in knowing:

1. How much preventative work is needed to cost-effectively slow down the arrival of CWD, and hence the long-term prevalence?
  a. What contribution do proactive measures at captive cervid facilities make to reducing long-term prevalence?
  b. How effective are other proactive measures in reducing prevalence?
2. What level of reactive measures most cost effectively reduce long-term CWD prevalence?
  a. Are there diminishing returns in investment in sharpshooting?
  b. How does investment in surveillance for detection of CWD interact with reactive measures to reduce long-term prevalence?
3. What is the most cost-effective combination of proactive, surveillance, and reactive actions to reduce long-term prevalence of CWD in Vermont?

To investigate these questions, we simulated 256 additional strategies, looking systematically at combinations of different levels for the action elements. We first present the results of the strategies on prevalence, and then describe results that also consider costs.

The analysis of the first 40 scenarios provided insights into the first three related questions: the relative effects of the proactive actions, including captive cervid farm closures and double fencing, in reducing long-term prevalence of CWD in the wild deer population. Of the five possible combinations of actions at captive deer facilities, we found that all the actions associated with captive facilities reduced long-term prevalence in wild deer by about 1.0- to 1.5-percent at year 50 (or about 2,100 fewer CWD-positive deer in that year, assuming a population size of 140,000) compared to the status quo approach (fig. 9).

**Figure 9.**
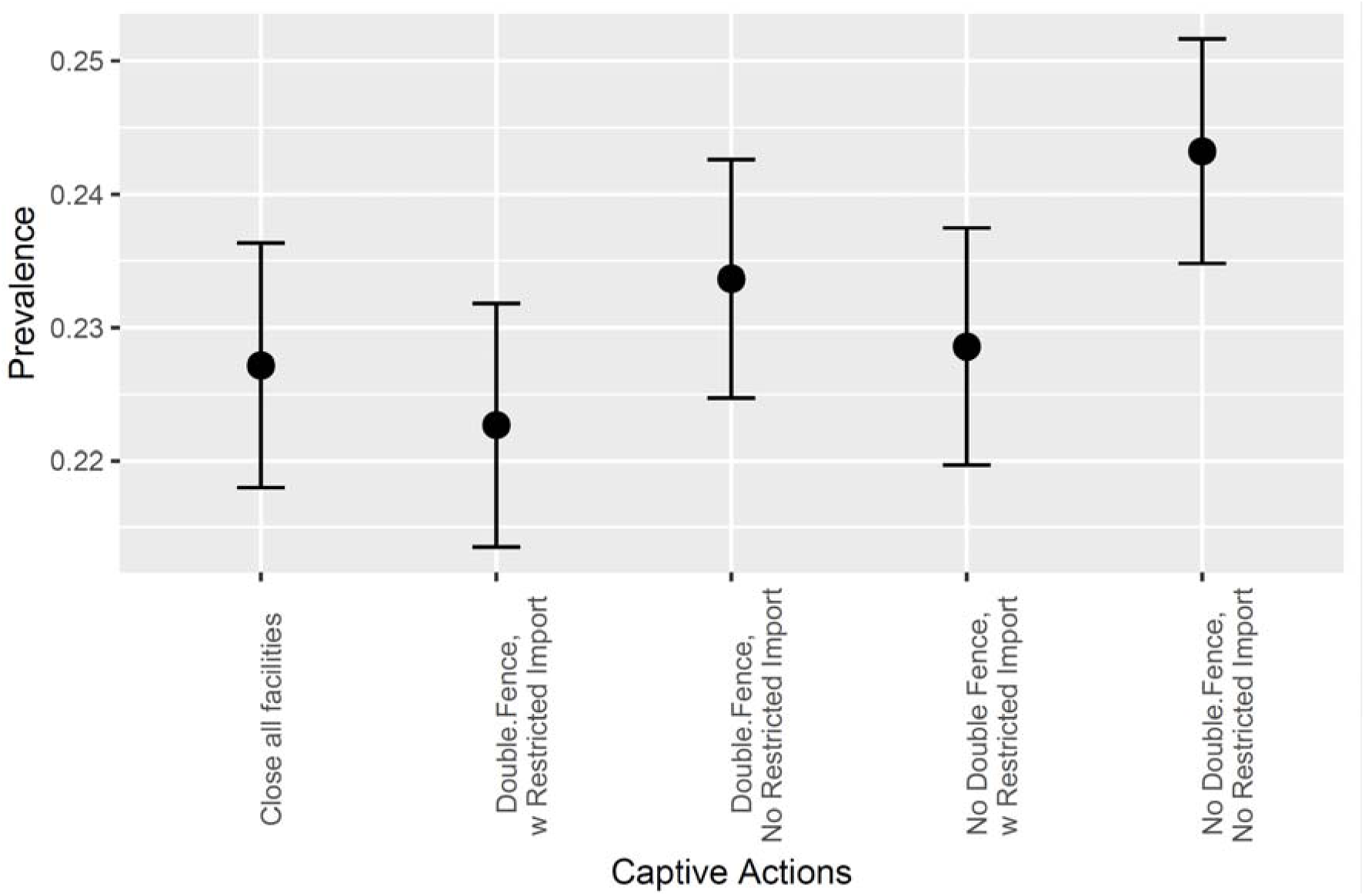
Effectiveness of proactive measures at captive facilities in Vermont. Prevalence of chronic wasting disease in the wild Vermont deer population at year 50 of the simulation, for 5 alternative approaches to captive deer management, at low levels of other proactive measures, and at low reactive levels. Symbols show the mean prevalence, error bars show the 95-percent confidence interval for the mean.

The patterns in the effects of the proactive actions at captive farms roughly hold up in combinations with other proactive actions taken to interrupt CWD introduction pathways involving wild deer (fig. 10). Restricting import of carcasses by residents hunting out-of-state reduces long-term prevalence by about 1-percent after 50 years, and issuance of carcass disposal guidance to hunters, taxidermists, and processors reduces prevalence by about 0.5-percent.

**Figure 10.**
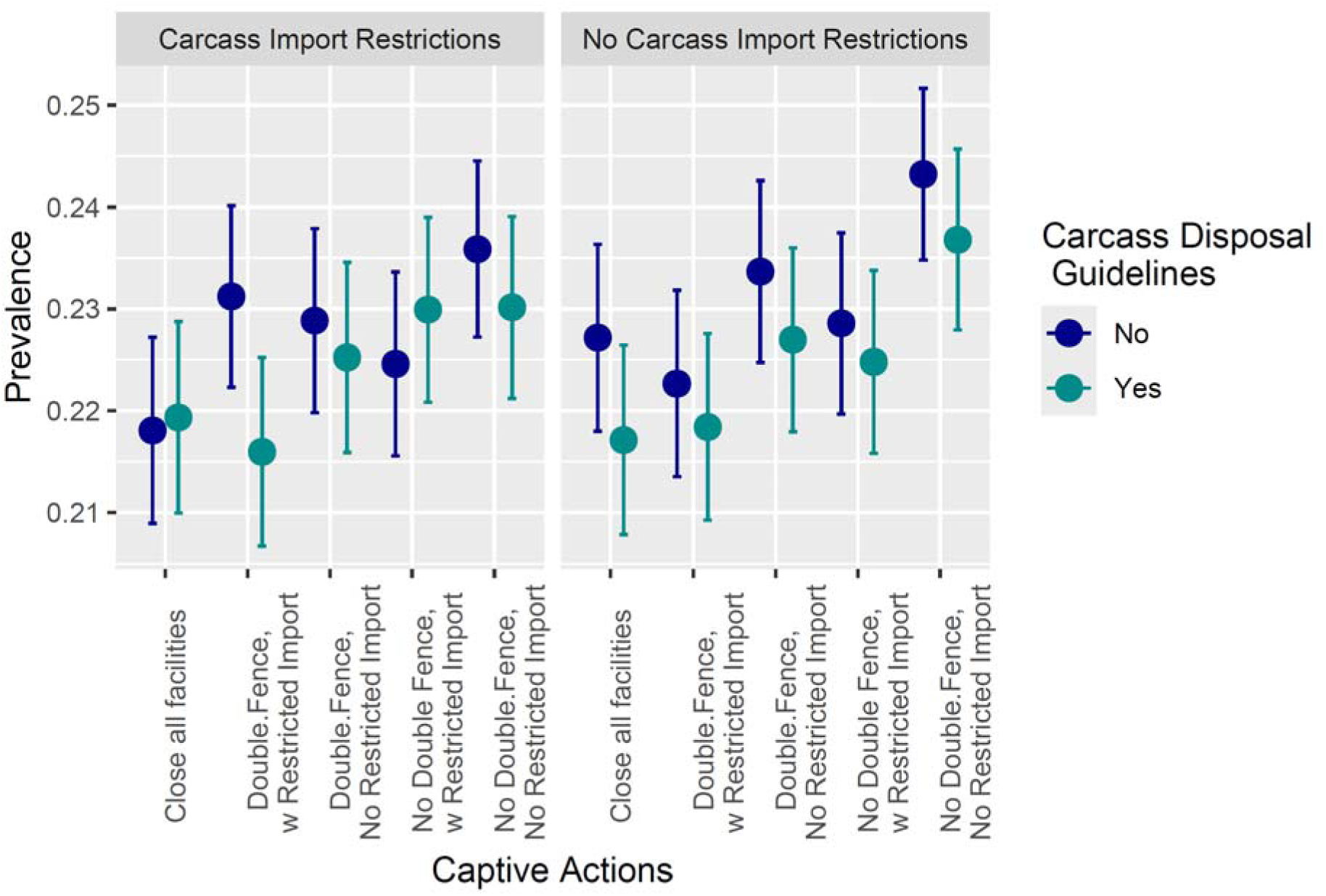
Effectiveness of all proactive measures, at low reactive levels. Prevalence of chronic wasting disease in the wild Vermont deer population at year 50 of the simulation, for 5 alternative approaches to captive deer management, in combination with 2 levels of restrictions on carcass movement by hunters hunting out-of-state, and two levels of guidance for carcass disposal, at low reactive levels. Symbols show the mean prevalence, error bars show the 95-percent confidence interval for the mean.

The effects of the different proactive actions, however, are swamped by the collective effects of reactive actions (fig. 11), with high reactive effort reducing long-term prevalence by about 13-percent compared to low reactive effort.

**Figure 11.**
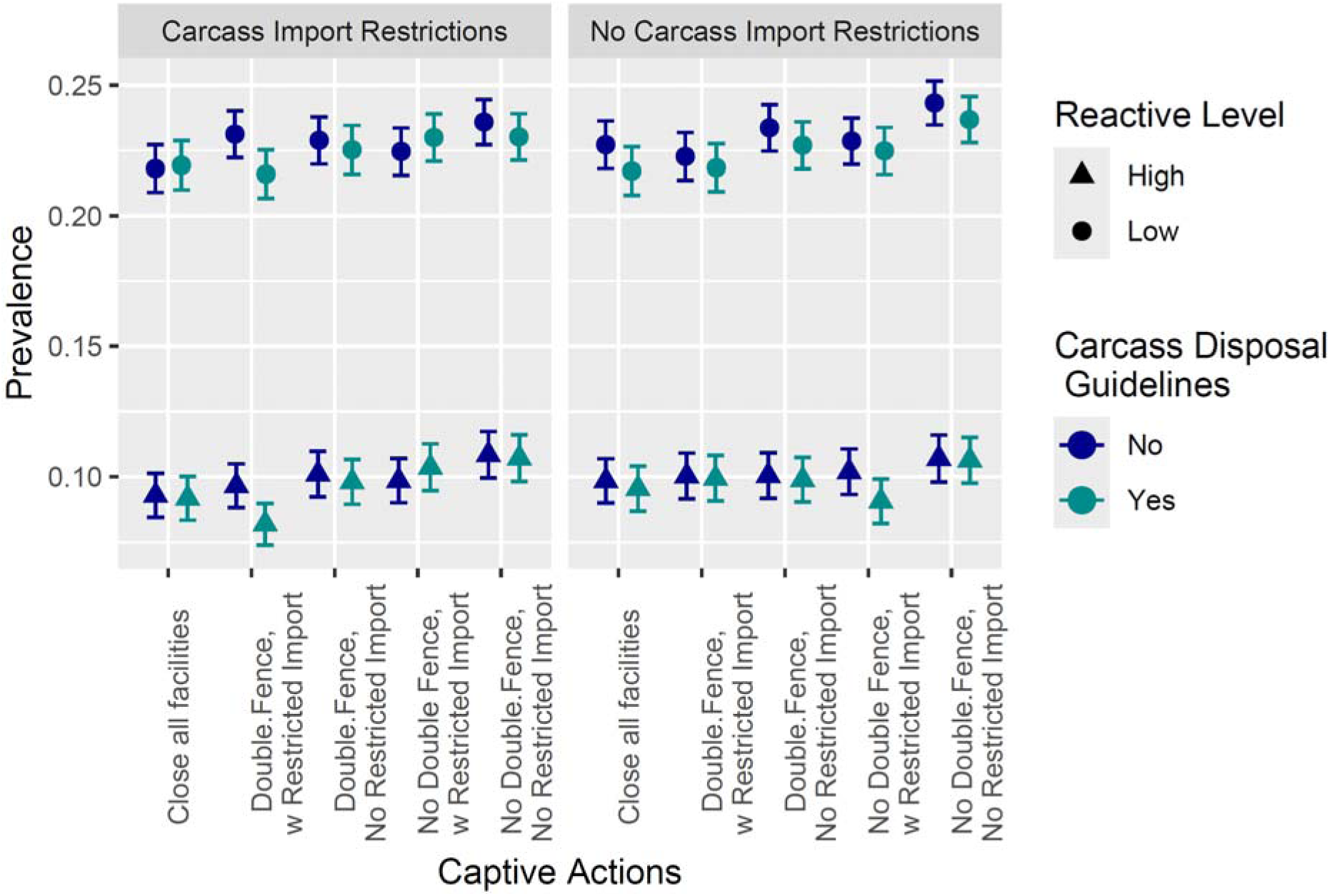
Comparison of proactive and reactive measures for the management of chronic wasting disease in white-tailed deer. Prevalence of chronic wasting disease in the wild Vermont deer population at year 50 of the simulation, for 5 alternative approaches to captive deer management, in combination with 2 levels of restrictions on carcass movement by hunters hunting out-of-state, and two levels of guidance for carcass disposal, at both high and low reactive levels. Symbols show the mean prevalence, error bars show the 95-percent confidence interval for the mean.

The second set of questions analyzed by an additional 216 scenarios focused on the efficacy of reactive measures at reducing long-term CWD prevalence. All the key reactive elements of the strategies (pre-detection surveillance, post-detection harvest rate, and sharpshooting) contribute to reducing the long-term prevalence of CWD in the wild population, with varying effectiveness (fig. 12).

**Figure 12.**
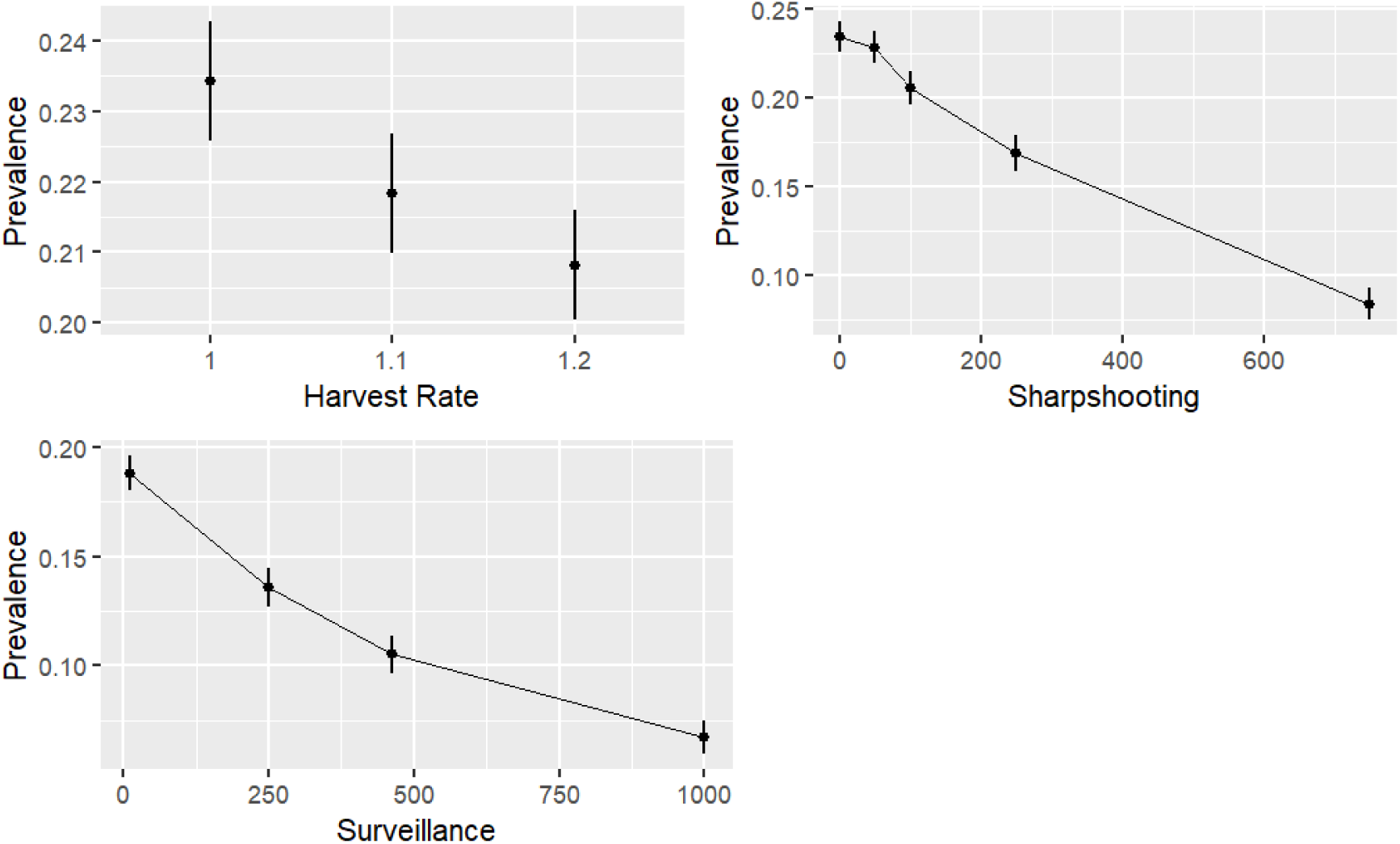
Effects of individual reactive measures. Prevalence of chronic wasting disease in the wild Vermont deer population at year 50 of the simulation, for (A) three levels of post-detection harvest rate (with sharpshooting at 0/yr and surveillance at 250/yr); (B) five levels of sharpshooting (with harvest at status quo and surveillance at 250/yr); and (C) four levels of pre-detection surveillance (with post-detection harvest at 1.2 and sharpshooting at 250/yr). The proactive actions are at status quo (low) levels. Symbols show the mean prevalence, error bars show the 95-percent confidence interval for the mean.

Elevating the harvest rate after detection of CWD helps to reduce the density of deer, which reduces CWD transmission rates, and slows the spread of the disease. This effect is small compared to the effects of the other reactive elements. Increasing the harvest rate by 10% lowers the prevalence at year 50 by about 1.5% and increasing the harvest rate by 20% lowers prevalence by about 2.5%, under moderate surveillance and no sharpshooting (fig. 12A). This effect appears to be additive to the other elements, and so the effect size holds in other combinations of sharpshooting and surveillance (figs. 13-14).

The effects of sharpshooting and surveillance interact with each other because increased surveillance allows detection of CWD at lower prevalence, which makes the sharpshooting effort more effective (fig. 13). At very low levels of surveillance (testing of 10 deer that look unhealthy, per year), sharpshooting is not very effective; removing 750 deer/year only reduces long-term prevalence by about 5%, because detection occurs so late. As surveillance effort increases, the effectiveness of sharpshooting increases, so that with surveillance of 463 carcasses/year, sharpshooting 750 deer/year decreases the long-term prevalence by about 20% (fig. 13C). At very high levels of surveillance (1,000/year), the non-linearity of the sharpshooting effect becomes evident, with attenuating effects at high levels of effort (fig. 13D).

**Figure 13.**
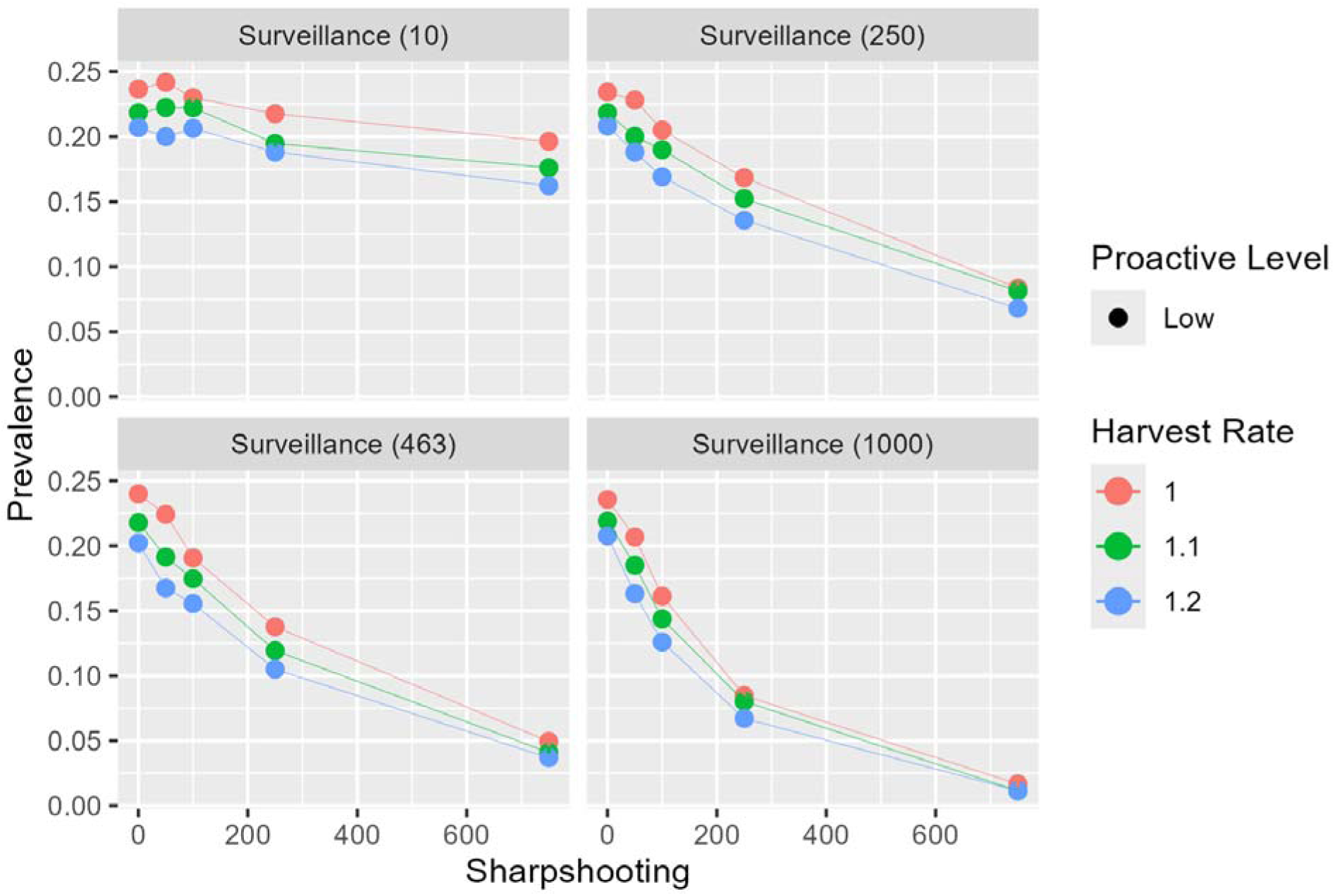
Effect of sharpshooting in the context of pre-detection surveillance. Prevalence of chronic wasting disease in the wild Vermont deer population at year 50 of the simulation as a function of sharpshooting effort (removals/year), at four levels of pre-detection surveillance, three levels of post-detection harvest rate, and at a low level of proactive effort. Symbols show the mean prevalence.

The benefit of pre-detection surveillance primarily comes through its ability to direct sharpshooting; the effect of post-detection harvest levels is not strongly affected by surveillance. When there is no sharpshooting, increasing pre-detection surveillance has no benefit (fig. 14). The benefit of increased surveillance becomes particularly noticeable once the sharpshooting rate rises to 100−250/year (fig. 14). The combination of very high surveillance (1,000/year) and high sharpshooting (750/yr) results in very low long-term prevalence of CWD (fig. 14). At these levels, any new incursion of CWD to the state is discovered at a stage when sharpshooting resources can eradicate the incursion or keep growth in check.

**Figure 14.**
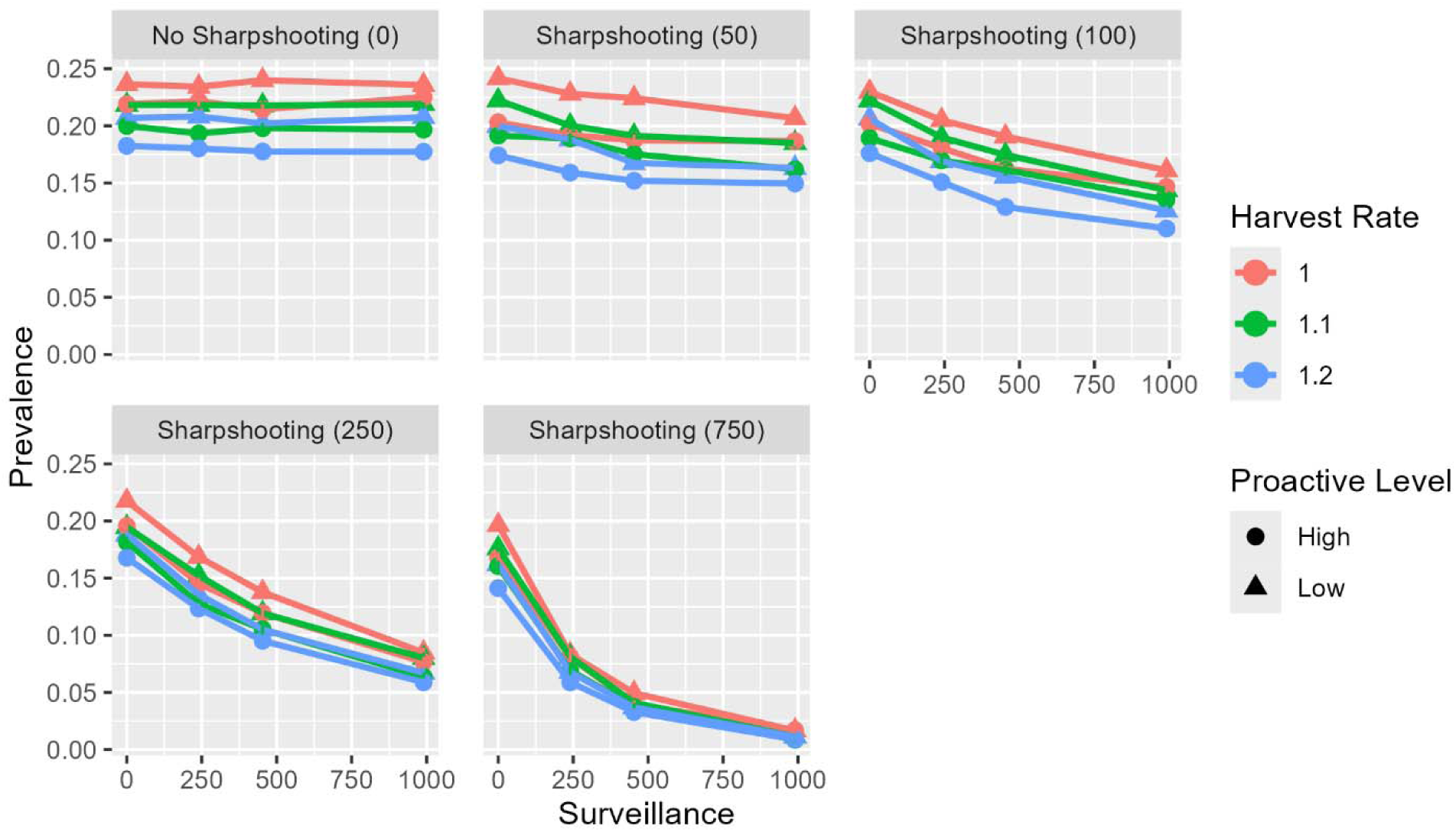
Effect of surveillance in the context of investment in sharpshooting. Prevalence of chronic wasting disease in the wild Vermont deer population at year 50 of the simulation as a function of pre-detection surveillance (deer sampled/year), at five levels of sharpshooting effort, three levels of post-detection harvest rate, and two levels of proactive effort. Symbols show the mean prevalence.

## 11 Phase II – Multi-Objective Optimization Results

### 11.1 Pareto Efficiency

While there are several actions that can be taken to effectively reduce long-term prevalence of CWD in Vermont, these benefits come at a cost. Proactive measures at captive facilities, pre-detection surveillance, and post-detection sharpshooting can all be quite expensive. To investigate which actions are most cost-effective and where the gains might be worth the costs, we created “Pareto plots” that graphically illustrate the relationship between the long-term prevalence and cumulative 50-year management costs. As an overview, consider the plot of all 256 scenarios (fig. 15). Since we would like to minimize both the long-term prevalence and the management costs, the best performing strategies are toward the bottom left of the figure. In that part of the figure, there is a frontier (known as a “Pareto frontier”) where you must spend more money to decrease the prevalence further. Everything above and to the right of the frontier is a “dominated alternative” in the sense that it is possible to find a strategy that is both more effective (lower prevalence) and cheaper (lower cost). Only one of the original six alternative strategies (*Kitchen Sink*) is on the Pareto frontier. But it is worth noting that this plot only considers two of the eight fundamental objectives, so there may be reasons to choose alternatives that are not on the frontier, because they are advantageous with regard to an objective that is not plotted. In the next few paragraphs, we examine the cost-efficiency of the components of the strategies.

**Figure 15.**
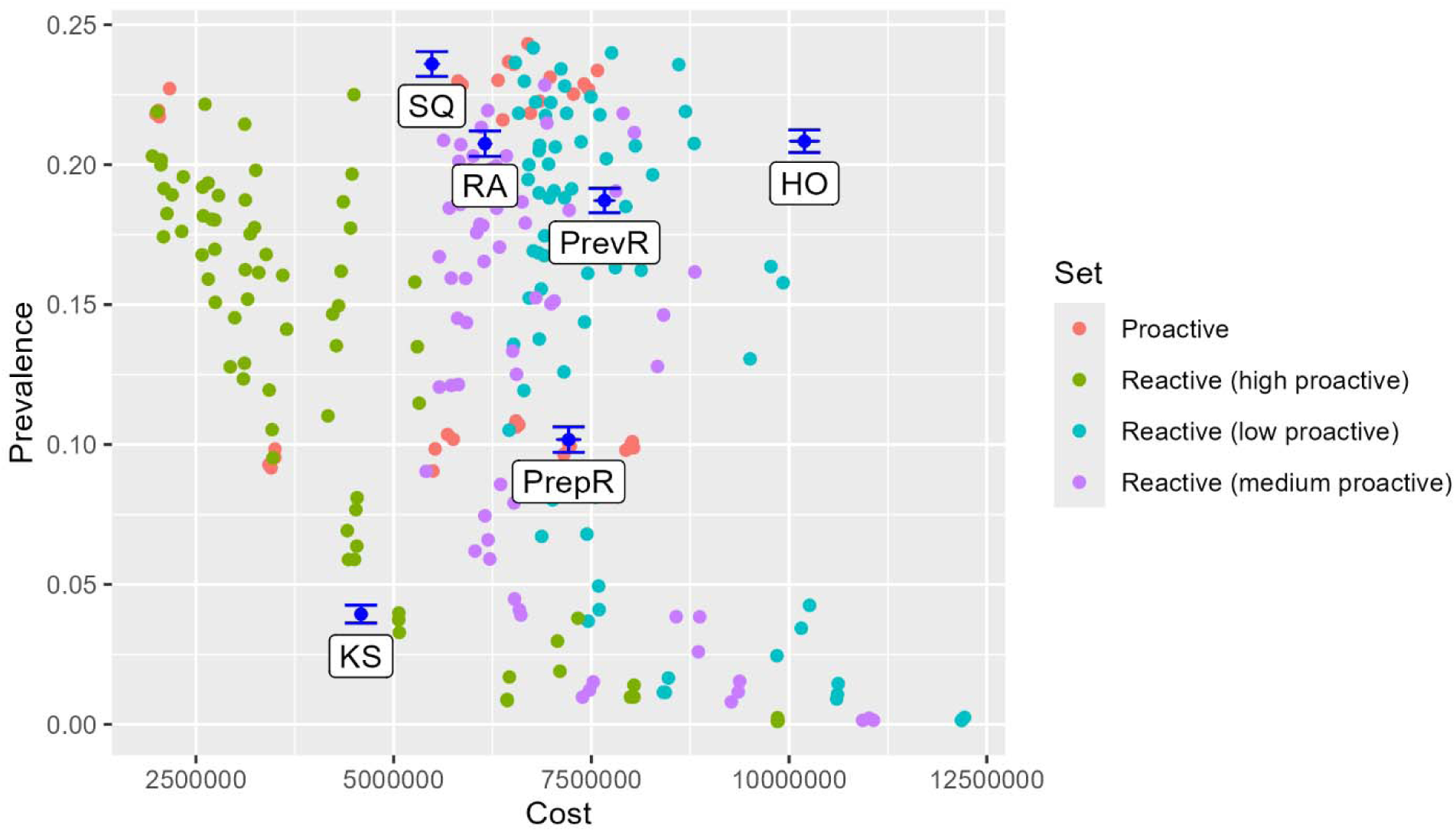
Pareto plot of cost-effectiveness. Prevalence of chronic wasting disease in the wild Vermont deer population at year 50 of the simulation plotted against the total cost to the management agencies over 50 years, for 256 new scenarios (in 4 sets) and the 6 original alternative strategies (blue). [SQ, Status Quo; HO, Heavy Outreach; RA, Reduce Arrival; PrepR, Prepare and React; PrevR, Prevent and React; KS, Kitchen Sink]

#### 11.1.1 Preemptive Captive Cervid Operation Closures

The cost-effectiveness of proactive measures at captive deer facilities is shown in figure 16, in the context of low reactive measures. The preemptive closure of all captive facilities results in a small reduction (about 1-percent) in long-term CWD prevalence in wild deer compared to the status quo, but more importantly, by closing those facilities before they become infected, it foregoes the future costs of closing infected facilities (estimated at $750,000 per infected farm). Notably, this also foregoes all the opportunity for those farms to operate in the interim, a trade-off that is not captured in the Pareto plot (fig. 15). Proactively double fencing all captive farms reduces the long-term prevalence slightly more, but is quite expensive, both because the fencing is expensive, and because a small fence failure rate still results in a need to close some infected farms in the future. (On average, the double fencing is estimated to cost $2.1M, but it only leads to avoidance of about $1.1M in farm-closure costs.) Finally, reducing the importation rate of captive cervids results in a small reduction in long-term prevalence (0.5 to 1.0-percent) as well as costs savings by lowering the risk of farm closure. The other proactive measures (increasing regulations on out-of-state carcass imports and increasing guidance about carcass disposal) produce reductions in prevalence and slight reductions in costs.

**Figure 16.**
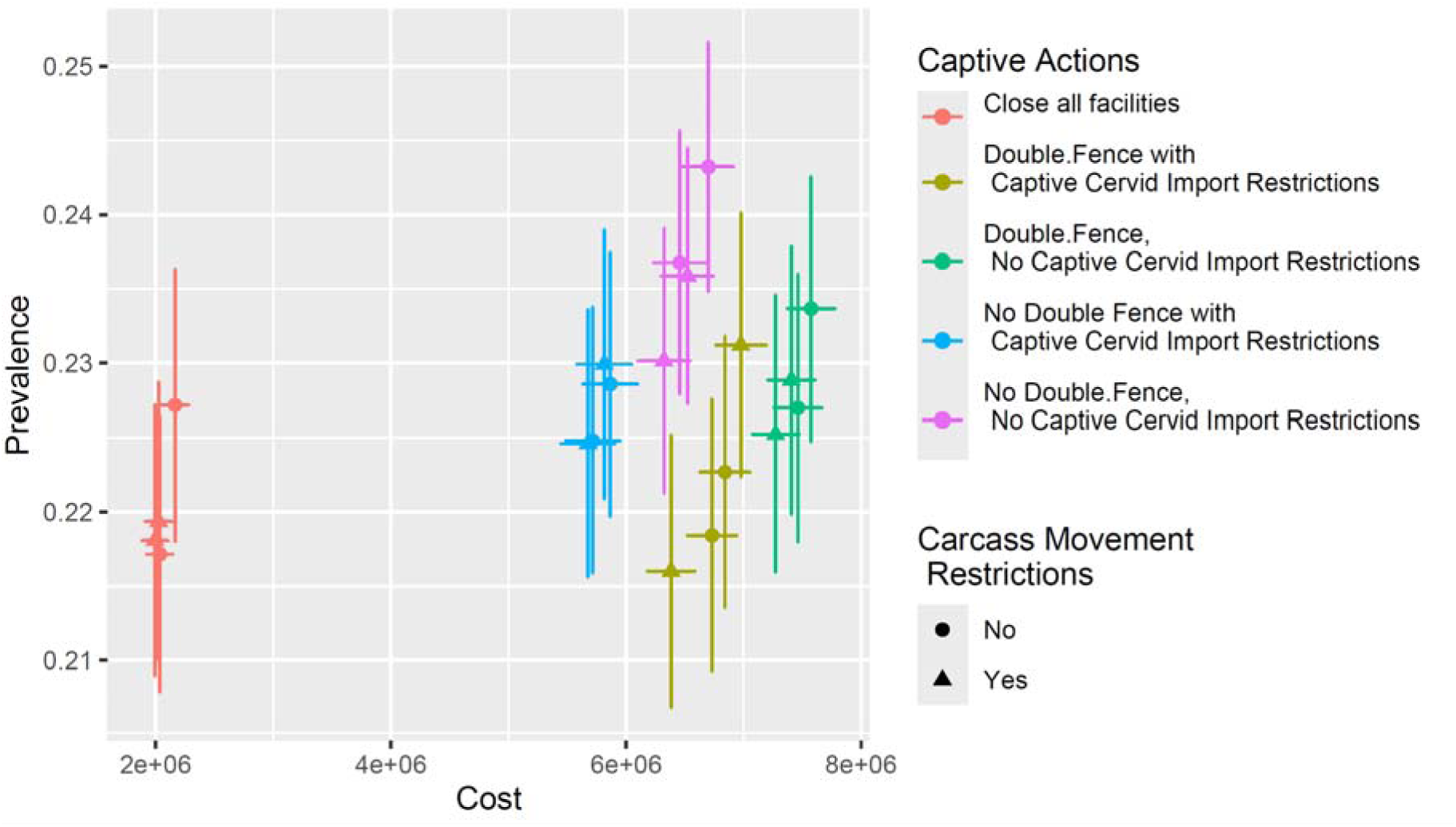
Cost-effectiveness of proactive measures for the management of chronic wasting disease in white-tailed deer. Prevalence of chronic wasting disease in the wild Vermont deer population at year 50 of the simulation plotted against the cost to the management agencies, comparing proactive actions, with reactive actions set at low levels (status quo surveillance, no sharpshooting, no post-detection increase in harvest). Error bars show the 95-percent confidence interval for the mean.

The cost-effectiveness of reactive measures is shown in figure 17, for the case where the proactive measures are at their maximum (notably, captive farms are proactively closed). In this case, a clear Pareto frontier appears, with further reductions in prevalence only possible with increased investment. Nearly all the Pareto-optimal solutions include a post-detection harvest rate increase of 1.2, which makes sense because no cost was attributed to this change in hunting regulations. With such a harvest rate, many combinations of sharpshooting and surveillance are Pareto-optimal—the exceptions are if surveillance is minimal (10/year) or if the surveillance investment outstrips the needs of the sharpshooting program.

**Figure 17.**
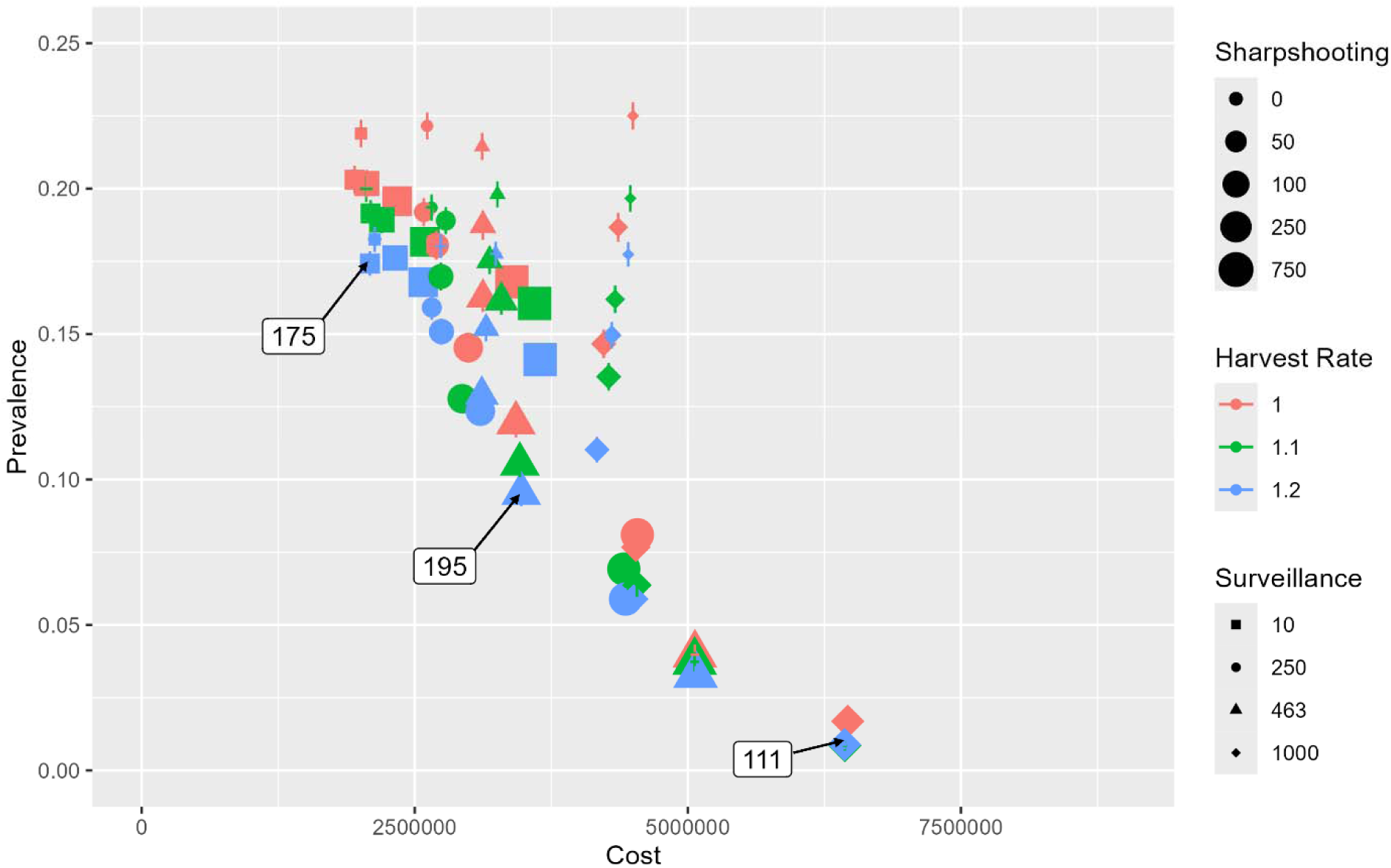
Cost effectiveness of reactive measures (with high proactive levels). Prevalence of chronic wasting disease in the wild Vermont deer population at year 50 of the simulation plotted against the cost to the management agencies, comparing reactive actions, with proactive actions set at high levels (preemptively close captive farms, impose carcass import restrictions, and increase carcass disposal guidance).

Three scenarios along the Pareto frontier illustrate the trade-off between prevalence and cost. Scenario 175 (surveillance of 10/year; sharpshooting 50/year) achieves a long-term prevalence of 0.174 at a total 50-year cost of $2.1M. Scenario 195 (surveillance of 463/yr; sharpshooting 250/yr) achieves a prevalence of 0.095 at a cost of $3.5M. Scenario 111 (surveillance of 1000/yr; sharpshooting 750/yr) achieves a prevalence of 0.009 at a cost of $6.4M. Thus, further gains in reducing long-term prevalence of CWD can be achieved with increased investment in surveillance and sharpshooting. The State’s decision, then, is in part influenced by how much it is willing to invest to lower the long-term CWD prevalence.

#### 11.1.2 No Preemptive Captive Cervid Operation Closures

The picture of cost-effectiveness is more complicated if the proactive measures do not include preemptively closing captive farms (fig. 18). Again, the most cost-effective strategies include a post-detection harvest rate increase of 1.2 (blue symbols). But now, the interaction of sharpshooting and surveillance does not all lie along the Pareto-frontier. Consider the strategies that include harvest at 1.2 and surveillance at 463 (the blue triangles): as the sharpshooting goes from 0 to 250, the prevalence decreases *and* the cost decreases. Even though there is an increase in cost owing to the sharpshooting, the total cost to the State *decreases* because the lower prevalence means fewer captive farms become infected; this lower cost of closure more than makes up for the increased sharpshooting costs. From sharpshooting of 250 to 750, an additional reduction in prevalence is possible, but it now comes at an additional total cost.

**Figure 18.**
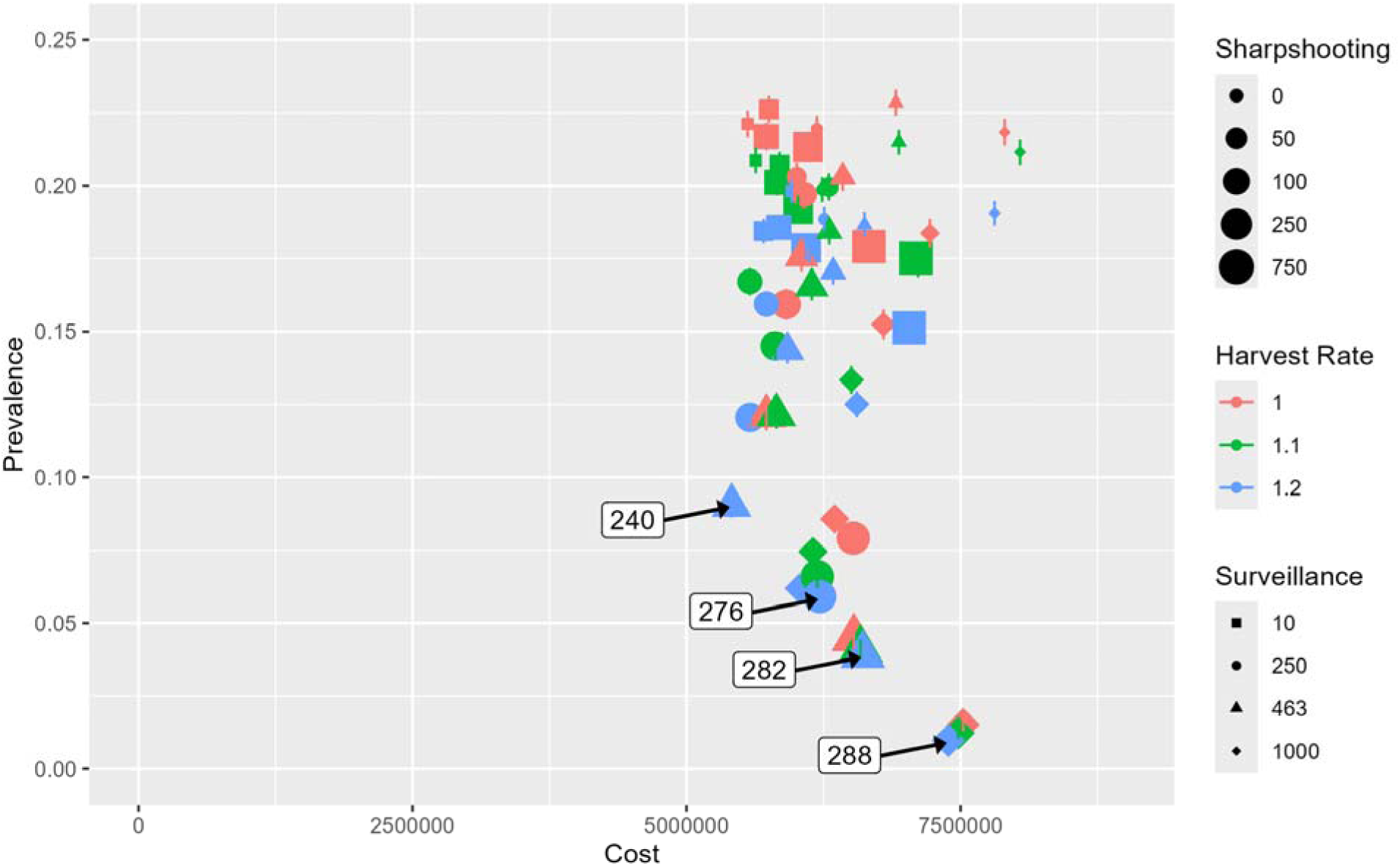
Cost effectiveness of reactive measures (with medium proactive levels). Prevalence of chronic wasting disease in the wild Vermont deer population at year 50 of the simulation plotted against the cost to the management agencies, comparing reactive actions, with proactive actions set at medium levels (keep farms open but reduce captive import rate, impose carcass import restrictions, and increase carcass disposal guidance).

Three scenarios along the frontier in figure 18 illustrate the trade-off between prevalence and cost. Scenario 240 (surveillance of 463/year; sharpshooting 250/year) achieves a long-term prevalence of 0.090 at a total 50-year cost of $5.4M. Scenario 276 (surveillance of 250/yr; sharpshooting 750/yr) achieves a prevalence of 0.059 at a cost of $6.2M. Scenario 282 (surveillance of 463/yr; sharpshooting 750/yr) achieves a prevalence of 0.039 at a cost of $6.6M. Scenario 288 (surveillance of 1,000/yr; sharpshooting 750/yr) achieves a prevalence of 0.010 at a cost of $7.4M.

The strategies on the frontier of figure 18 are not Pareto-optimal with regard to prevalence and cost alone (they fall to the right of the frontier on fig. 17), but they may be Pareto-optimal when prevalence and cost are considered along with the opportunity for captive farms to operate.

### 11.2 Consequence Table of Pareto Efficient Strategies

The seven new scenarios identified in figures 17 and 18 represent a set of alternatives that are Pareto-optimal with regard to long-term CWD prevalence, total agency cost, and opportunity to operate captive cervid farms. These alternatives are described in table 7.

**Table 7.** Description of seven pareto optimal strategies that span frontiers related to prevalence of chronic wasting disease in deer, costs to management agencies, and opportunities for agricultural activities. Common elements among all seven strategies include: increasing restrictions on import of out-of-state carcasses; increasing guidance on carcass disposal; and outreach to achieve 20% hunter-initiated testing of carcasses post-detection. [yr, year]

|  | 175 | 195 | 111 | 240 | 276 | 282 | 288 |
| --- | --- | --- | --- | --- | --- | --- | --- |
| *Proactive Elements* |  |  |  |  |  |  |  |
| Proactively close cervid farms | Yes | Yes | Yes | No | No | No | No |
| Reduce import of captives? | -- | -- | -- | Yes | Yes | Yes | Yes |
| Surveillance (pre-detection) | 10/yr | 453 | 1,000 | 453 | 250 | 453 | 1,000 |
| *Reactive Elements* |  |  |  |  |  |  |  |
| Hunting opportunity | 1.2 | 1.2 | 1.2 | 1.2 | 1.2 | 1.2 | 1.2 |
| Sharpshooting | 50/yr | 250 | 750 | 250 | 750 | 750 | 750 |
| *Results* |  |  |  |  |  |  |  |
| Mean time to arrival (yr) | 10.5 | 10.2 | 10.2 | 9.3 | 9.1 | 8.9 | 9.0 |
| Mean time to detection (yr) | +26.0 | +16.4 | +14.1 | +16.3 | +18.2 | +16.2 | +14.3 |
| Mean prevalence at detection | 0.028 | 0.002 | 0.001 | 0.002 | 0.003 | 0.002 | 0.001 |

A full consequence table for the seven new strategies shows the performance against all the fundamental objectives (table 8). Scenario 175 is the best performing of the seven strategies in terms of mean active hunters, sapling density, and total agency cost; Scenario 111 performs best with regard to size of the deer population, prevalence of CWD, and total CWD-positive animals consumed; and Scenario 288 performs best with regard to net annual revenue to associated businesses and cervid farm operations. The other strategies have intermediate performance on most metrics but may also present the best tradeoffs to the agencies.

**Table 8.** Consequence table showing the expected performance over 50 years of seven pareto-optimal strategies against eight objectives. In each row, the alternative that performs best is shaded green, the alternative that performs worst is shaded pink, and strategies that perform better than the status quo (as of 2024 regulations) are shaded yellow. Strategies that perform worse than the status quo are unshaded.

| Performance Metric | desired direction | 175 | 195 | 111 | 240 | 276 | 282 | 288 |
| --- | --- | --- | --- | --- | --- | --- | --- | --- |
| Size of deer population at year 50 | Max | 127,062 | 135,333 | 143,383 | 135,346 | 137,281 | 139,117 | 142,928 |
| CWD prevalence at year 50 | Min | 0.174 | 0.095 | 0.009 | 0.090 | 0.059 | 0.039 | 0.010 |
| Mean annual active hunters over 50 years | Max | 61,216 | 59,678 | 59,320 | 59,593 | 59,807 | 59,446 | 59,209 |
| Sapling density at year 50 (saplings/ha) | Max | 2,895 | 2,869 | 2,843 | 2,869 | 2,863 | 2,857 | 2,845 |
| Mean total cost to agencies (\$ over 50 yr) | Min | 2,089,321 | 3,474,550 | 6,433,961 | 5,410,681 | 6,214,995 | 6,609,199 | 7,388,282 |
| Total CWD-positive animals consumed (#) | Min | 18,688 | 9,884 | 924 | 10,341 | 6,640 | 4,208 | 1,228 |
| Net annual revenue for associated business (\$) | Max | 1,780,065 | 1,798,821 | 1,804,516 | 1,802,436 | 1,798,348 | 1,802,319 | 1,807,804 |
| Cervid farm operations (total farm-years) | Max | 0 | 0 | 0 | 408 | 412 | 414 | 420 |

A cost breakdown shows that the costs accrue in quite different ways for the seven new strategies (table 9). A significant component of the total cost to the state agencies is the cost to VAAFM related to captive farms. Scenarios 175, 195, and 111 incur upfront costs to preemptively close the existing cervid farms, but in doing so, forego future, more expensive costs of closing farms that become infected with CWD. On the other hand, Scenarios 240, 276, 282, and 288 incur some small expenses to reduce the rate of captive cervid importation, then incur greater expenses to close any farms that eventually become infected. From a future expense perspective, VAAFM benefits from investments made by VDFW to reduce prevalence of CWD in the wild deer population, because the risk of farms becoming infected is reduced by actions that reduce disease in wild deer. Sharpshooting costs, presumably borne by VDFW, can vary widely (from $121,000 over 50 years if the goal is only to remove 50 deer per year to $3,600,000 over 50 years if the goal is to remove 750 deer per year). Surveillance cost includes not only the cost of testing some targeted number of samples of hunter-killed carcasses, it also includes voluntary hunter-requested testing of meat prior to consumption in line with current public health guidance, which can be fairly extensive once outreach campaigns are successful and statewide prevalence rises.

## 12 Discussion

Based on the results of this analysis, CWD is a foreseeable, if not an imminent, risk to the wild and captive deer populations in Vermont. The work in this report was designed to support a multi-agency group in navigating a complex decision problem associated with the arrival and spread of CWD in a U.S. state. The problem was complex because of a multitude of tradeoffs in the performance of different strategies as well as uncertainties associated with the timing of arrival, disease spread dynamics, and the efficacy of actions designed to surveil for, and proactively and reactively manage the disease itself. Additionally, the suite of agencies working on the collaborative plan have differing missions and authorities and determining which sets of actions are most effective across these differing agency goals while also taking advantage of the expanded set of authorities is a difficult task. While there remain many open questions related to the spread to, and management of CWD in, Vermont, and other areas of the northeastern U.S., the decision analysis tools and computer models contained in this report are expected to support the development of CWD management planning documents.

The evaluation of alternatives occurred through two phases designed to overcome decision impediments related to uncertainty and multi-objective tradeoffs. The first phase focused on a discrete set of six alternatives that helped estimate the range of possible outcomes and the intensity of tradeoffs embedded in each alternative. The strategies were designed to understand how different, discrete combinations of intensive and lax surveillance, proactive, and reactive actions performed on the full set of performance metrics that were important to achieve. Through this evaluation, several things were made clear. First, the most important performance metric was to reduce the long-term prevalence of CWD in wild deer. Under status quo management, the expected arrival is in 7.7 years. Through some proactive efforts, this could be delayed until about 10.6 years, by reducing import of captive deer and import of out-of-state carcasses. Eventually, however, the main source of arrival will be dispersal of infected deer across the border from neighboring states or provinces, a process that had no associated management actions. Relatedly, the second important finding from the first phase of the project was that the *Kitchen Sink* strategy was preferred across all agencies, but uncertainty remained about whether the combination of actions within that strategy was the most efficient way to achieve low CWD prevalence. The final key finding from phase I was that most of the performance metrics had little effect on the ability to distinguish among strategies; however, the need to preserve captive farming opportunities was highly valued and this made the selection of any alternative that called for restrictions on existing farms difficult to do.

Moving into phase II of the project, the technical team focused primarily on optimizing the set of actions based on their ability to lower both costs and CWD prevalence simultaneously. Building on findings from phase I, it was also important to separate out those strategies that further restricted or eliminated captive cervid farming opportunities from those that maintained the already stringent set of management policies. In evaluating the actions, it was clear that the most effective way to manage CWD once it arrives will be early detection coupled with focused removal (through sharpshooting or related targeted efforts). This approach can handle isolated arrivals of CWD (through out-of-state carcass import or captive escape) and may even be able to keep a steady stream of arrivals through dispersal at bay, enough to possibly keep the prevalence of CWD in the State quite low over the next 50 years. The success of such an approach very much depends on the degree of investment in an appropriate mix of surveillance and sharpshooting, and the long-term acceptability of such an intensive management strategy. In other States, there has been some resistance by members of the public to long-term sharpshooting (e.g., Illinois and Missouri). Efforts to better understand the design and communication of targeted removals in terms of maintaining public support would be critical to adopting a strategy that relies on long-term implementation. Other proactive and reactive measures can contribute to the long-term suppression of CWD in the State, but to a lesser degree than surveillance and sharpshooting.

There are nuanced and important interactions between the outcomes valued by the different agencies. Notably, there are potential benefits to VAAFM of investment by VDFW to suppress CWD, through reducing the risk of captive cervid farms being infected by CWD from wild deer. As noted above, if captive cervid farms remain operational, then investment in surveillance (up to 463 deer/year) and sharpshooting (up to 250 deer/year) more than pays for itself in foregone costs to close infected cervid farms. Investment by VDFW to suppress CWD also has substantial potential benefits to VDH, by reducing the amount of CWD-infected venison consumed by humans. This benefit may be enhanced by cooperation between the agencies, as outreach by VDH could help increase the rate at which hunters have their deer tested for CWD.

It is important to highlight the existing actions by VAAFM that reduce CWD introduction risk to captive and wild deer. Under the status quo, VAAFM restricts imports to an average of 2.87 animals annually and only approves permits for animals originating in CWD-certified herds with mandatory reporting and quarantine requirements as stringent as those set by Vermont. In combination with other existing work to inspect, inventory, and test deceased captive animals for CWD, the benefit of this work is likely substantial. While we did not analyze any alternatives where current actions by VAAFM are reduced, it is possible that CWD risk to captive and wild deer would increase substantially with less stringent requirements.

The development of a long-term strategic plan for Vermont is a choice among myriad combinations of actions, but the analysis of a large number of scenarios herein helps to sort through the combinations. There are a number of elements that appear to be possible, in the sense that they produce benefits with relatively low costs:

- Continuing and even enhancing the already stringent regulations on the import of captive cervids, which may possibly be infected with CWD, moves toward eliminating one of the pathways of arrival.
- Increasing regulations and outreach regarding movement of out-of-state deer carcasses into Vermont could help to reduce another pathway of arrival.
- Increasing, or preparing to increase, deer harvest rates is a way to reduce deer density, which in turn could reduce CWD transmission rates once it arrives.
- Enhancing guidance on carcass disposal methods can reduce environmental transmission of CWD once it arrives.

The analysis herein suggests there are three key aspects of a long-term strategic plan that induce important trade-offs among objectives, and thus invite substantive deliberation:

- For VAAFM, how should captive cervid facilities be handled? Is it better to preemptively close them (before CWD arrives in the State) to offset future costs, or to keep them open for the benefit of private farming opportunity, knowing there may be eventual costs of closure of infected farms?
- For VDFW, how much to invest in surveillance for early detection of CWD arrival in the State?
- For VDFW, in partnership with USDA APHIS, how much investment should be prepared for sharpshooting or other targeted removal methods?

The seven new alternative scenarios presented in Tables 7−9 capture the trade-offs embodied in these questions, and the range of choices represents different ways to balance those trade-offs. During a meeting where the interagency team discussed phase II strategies, including a particular focus on the seven new Pareto-efficient alternatives, there was a collective interest in further developing strategy 282 for the Vermont CWD management plan (outline in Appendix 15.4). The elements of that strategy that were most attractive to the participating agencies include: it keeps prevalence low over the course of the next 50 years; it maintains the opportunity to continue captive cervid farming (albeit under conditions similar to or more restrictive than currently exist); and it leads to a delayed arrival of CWD and early detection. The drawbacks of 282 were discussed at length and include the need for substantial investment in CWD over the next decade to prepare for and react to its arrival.

**Table 9.** The cost breakdown for seven alternative strategies. The costs shown are the total cost over the 50-year simulation horizon, with the mean taken over 1,000 replicates. Costs are separated by proactive and reactive management.

|  | 175 | 195 | 111 | 240 | 276 | 282 | 288 |
| --- | --- | --- | --- | --- | --- | --- | --- |
| *Proactive costs* |  |  |  |  |  |  |  |
| Hunter Carcass Regulations | \$0 | \$0 | \$0 | \$0 | \$0 | \$0 | \$0 |
| Carcass Disposal Guidelines | \$40,000 | \$40,000 | \$40,000 | \$40,000 | \$40,000 | \$40,000 | \$40,000 |
| Preemptively Close Captive Facilities | \$150,000 | \$150,000 | \$150,000 | \$0 | \$0 | \$0 | \$0 |
| Restrict Captive Cervid Imports | \$0 | \$0 | \$0 | \$16,665 | \$17,742 | \$16,294 | \$15,122 |
| Double Fencing | \$0 | \$0 | \$0 | \$0 | \$0 | \$0 | \$0 |
| Surveillance (pre-detection) | \$35,543 | \$1,182,409 | \$2,333,700 | \$1,140,739 | \$657,375 | \$1,114,348 | \$2,226,400 |
| *Reactive costs* |  |  |  |  |  |  |  |
| Sharpshooting | \$121,113 | \$1,055,790 | \$3,464,505 | \$1,096,290 | \$3,065,175 | \$3,365,820 | \$3,609,360 |
| Surveillance (post-detection) | \$1,742,665 | \$1,046,351 | \$445,756 | \$1,097,987 | \$778,703 | \$631,986 | \$478,150 |
| Close Infected Cervid Farms | \$0 | \$0 | \$0 | \$2,019,000 | \$1,656,000 | \$1,440,750 | \$1,019,250 |
| **Total costs** | **\$2,089,321** | **\$3,474,550** | **\$6,433,961** | **\$5,410,681** | **\$6,214,995** | **\$6,609,199** | **\$7,388,282** |

The actual plan itself can be more nuanced than what this report considers and there is more work to do to build stakeholder support. For example, because arrival of CWD is not likely to be imminent, a surveillance plan could ramp up over the next 10 years or so, allowing VDFW to slowly develop the resources, and allowing hunters to slowly adapt to new protocols. There could also be some efficiencies in surveillance, especially in coordination with neighboring states (e.g., New York) and captive cervid farms, to deploy surveillance efforts in places where arrival is more likely. Another nuanced modification could be in the design of reactive sharpshooting. The scenarios herein assumed that once CWD was detected, then the intended level of sharpshooting would be implemented and continue indefinitely, but, in fact, reactive sharpshooting could be quite responsive to surveillance data on a year-to-year basis. That said, the decision framing elements including the objectives of the interagency team, the actions considered, and the computer models developed, may serve as the technical basis for the State’s response plan as it is developed.

## 13 Summary

This report focuses on the use of structured decision making to frame and evaluate the next Vermont chronic wasting disease (CWD) management plan for deer and moose. Chronic wasting disease is spreading across North America, and in locations where it occurs, it causes mortality of infected animals, declining population productivity, and threats to hunting, wildlife viewing, cervid farming, and agency budgets. To position the state of Vermont to address the threats posed by CWD, the Vermont Department of Fish and Wildlife worked with six other State and Federal agencies to evaluate proactive and reactive elements of their management plan. Those elements were analyzed in various combinations (scenarios) that captured differing philosophies behind CWD management as well as rough levels of investment around which to build a long-term strategic plan. We found that the agencies preferred strategies that kept CWD as low as possible while also preserving the opportunity to continue captive cervid farming and operating cost-efficient conservation programs.

## Acknowledgments

We thank the agencies and their staff that generated much of the data that informed the analyses in this report. We also thank two reviewers, M. Feehan and B. Hanley, for contributing their expertise and providing comments and revisions that greatly improved the paper.

## 15 Appendices

## 15.1 Description of Phase I Alternative Strategies

### Alternative 1: Status quo

This alternative calls for a minimal proactive and reactive approach to the management of chronic wasting disease (CWD). VDFW, Vermont Department of Fish and Wildlife; APHIS, Animal and Plant Health Inspection Services; VDH, Vermont Department of Health; VAAFM, Vermont Agency of Agriculture, Food and Markets.

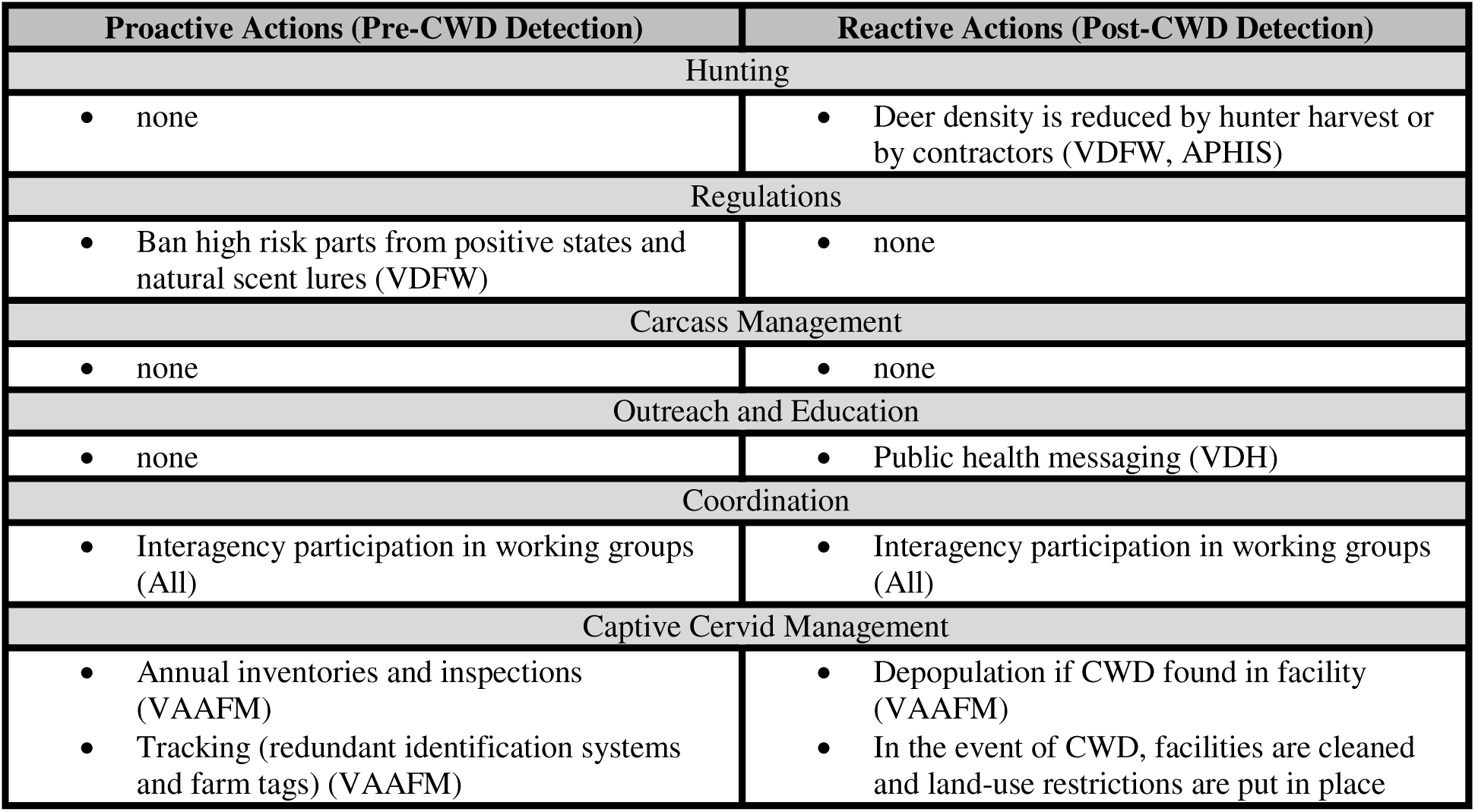
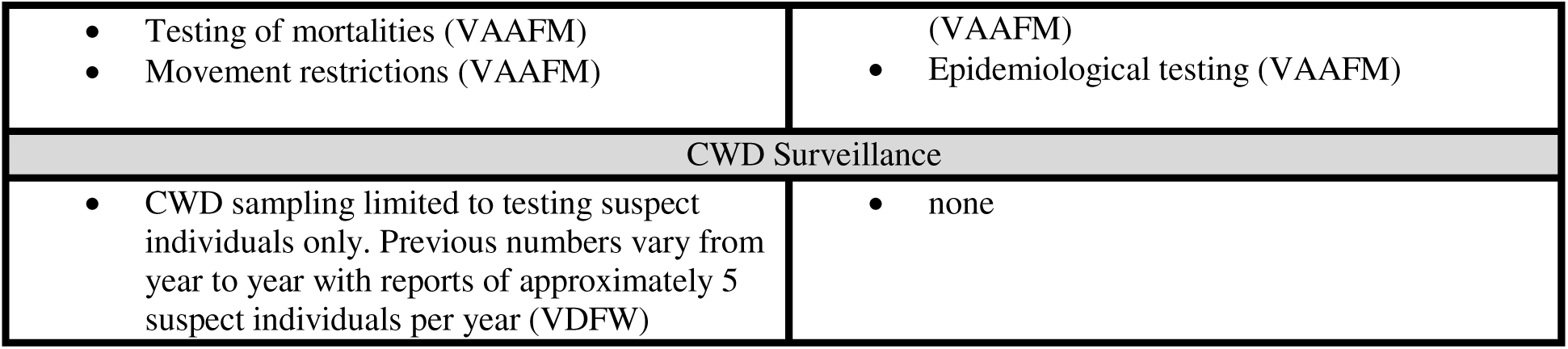

### Alternative 2: Heavy outreach to inform individual decisions

Outreach and education efforts emphasize important information about chronic wasting disease (CWD). By developing and disseminating outreach materials, task force agencies provide pertinent information that may inform voluntary action by hunters and the public. Additional resources are provided for surveillance to ensure that the information shared with the public is current. Outreach actions are used intensively as a proactive and reactive action, but all other actions are minimal prior to and after CWD detection. [VDFW, Vermont Department of Fish and Wildlife; USDA-APHIS, U.S. Department of Agriculture Animal and Plant Health Inspection Services; VDH, Vermont Department of Health; VAAFM, Vermont Agency of Agriculture, Food and Markets]

| Proactive Actions (Pre-CWD Detection) | Reactive Actions (Post-CWD Detection) |
| --- | --- |
| Hunting |  |
| • none | • Deer density is reduced by hunter harvest or by contractors (VDFW, USDA-APHIS) |
| Regulations |  |
| • Ban high risk parts from all states and scent lures (VDFW)<br>• No processing or taxidermy of out-of-state deer (VDFW) | • none |
| Carcass Management |  |
| • none | • Landowner guidelines (VDFW) |
| Outreach and Education |  |
| • Messaging during license sale, hunter education (VDFW)<br>• Guidelines for taxidermist/processor (VDFW)<br>• Messaging at hunt areas (VFPR, USFWS)<br>• Public health website (VDH)<br>• Sample collection training (USDA-APHIS)<br>• Cervid industry workshops (VAAFM)<br>• CWD grant proposals focused on outreach and education (APHIS) | • All actions in proactive (All)<br>• Expand public health messaging (VDH)<br>• Database of human exposures (VDH)<br>• Follow-up outreach to exposed humans (VDH) |
| Coordination |  |
| • Interagency participation in working groups (All) | • Interagency participation in working groups (All)<br>• Interagency maintenance of communications (e.g., spatial maps, signage) (All) |
| Captive Cervid Management |  |
| • Annual inventories and inspections (VAAFM)<br>• Tracking (redundant identification systems and farm tags) (VAAFM)<br>• Testing of mortalities (VAAFM)<br>• Movement restrictions (VAAFM)<br>• Emphasize non-CWD susceptible species for new farms (VAAFM) | • Depopulation if CWD found in facility (VAAFM)<br>• In the event of CWD, facilities are cleaned and land-use restrictions are put in place (VAAFM)<br>• Epidemiological testing (VAAFM)<br>• Double fencing encouraged (VAAFM) |
| CWD Surveillance |  |
| • Sampling to detect 1% CWD prevalence with 99% confidence statewide (VDFW) | • Monitor prevalence levels once detected (VDFW) |

### Alternative 3: Reduce the risk of CWD arrival

The emphasis is on actions that are proactive and occur before chronic wasting disease (CWD) is detected in new locations of the state of Vermont. The goal is to delay the arrival of CWD as long as possible and as such, actions are predominately proactive and highly intensive. [VDFW, Vermont Department of Fish and Wildlife; VFPR, Vermont Department of Forests, Parks and Recreation; USDA-APHIS, U.S. Department of Agriculture Animal and Plant Health Inspection Services; VDH, Vermont Department of Health; VAAFM, Vermont Agency of Agriculture, Food and Markets]

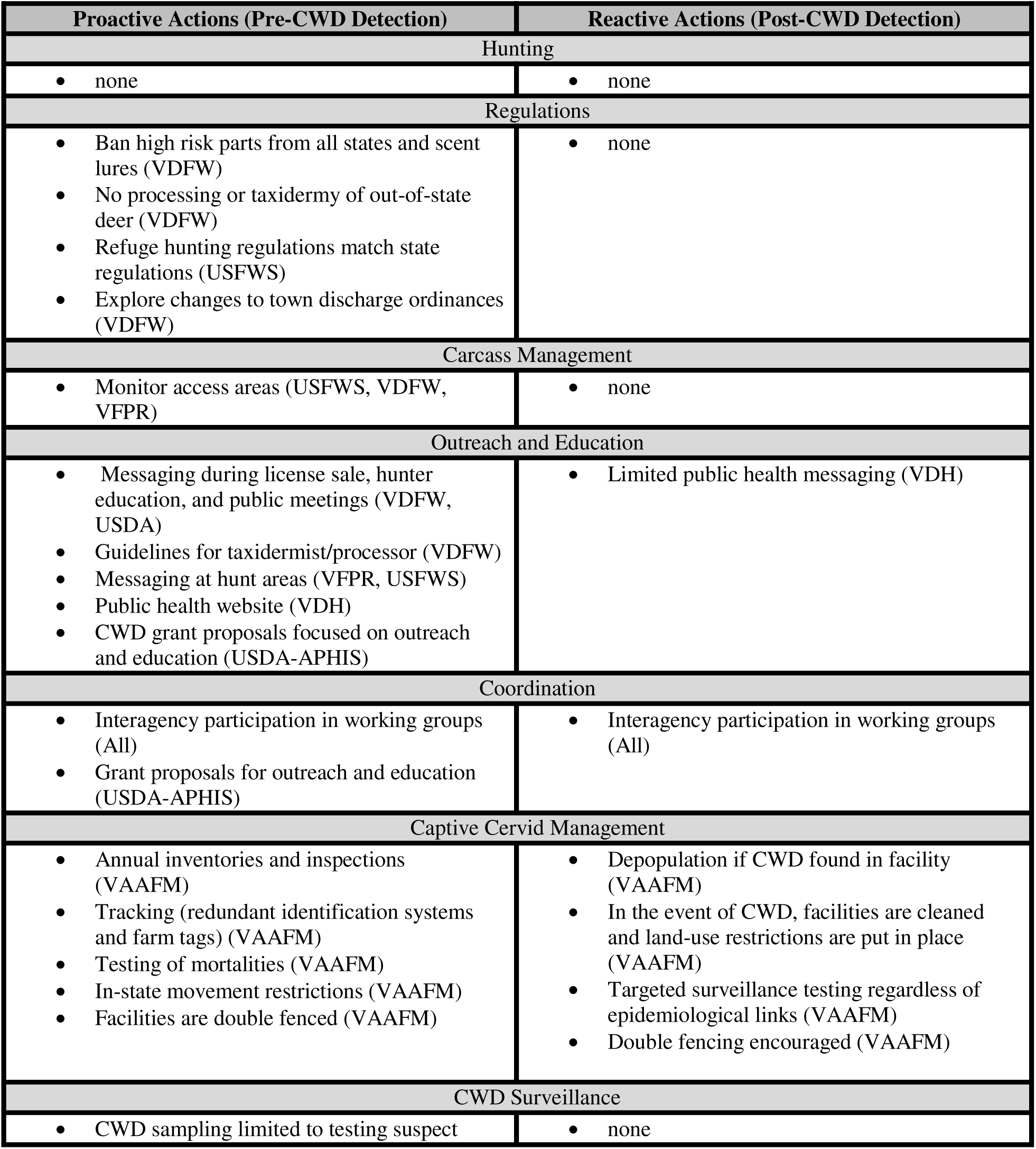
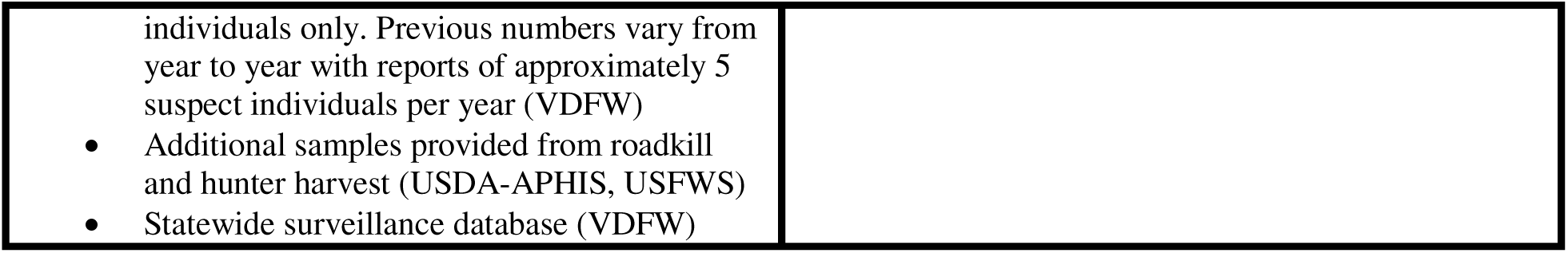

### Alternative 4: Prepare and react approach

This alternative creates opportunities for an aggressive reactive response to CWD detection and then implements those actions at first detection. Because the efficacy of response likely depends on early detection of CWD, there is also a large investment in surveillance in areas that are believed to be disease free. Pre-detection actions are limited to regulatory changes that allow for additional post-detection options in addition to other preparatory actions that allow for an intensive response. Reactive actions are expansive and include an expanded hunting season, targeted removal, and heavy outreach. [VDFW, Vermont Department of Fish and Wildlife; VFPR, Vermont Department of Forests, Parks and Recreation; USDA-APHIS, U.S. Department of Agriculture Animal and Plant Health Inspection Services; VDH, Vermont Department of Health; VAAFM, Vermont Agency of Agriculture, Food and Markets]

| Proactive Actions (Pre-CWD Detection) | Reactive Actions (Post-CWD Detection) |
| --- | --- |
| Hunting |  |
| • Work to increase hunting access (VFPR, USFWS, VDFW) | • Deer density is reduced by hunter harvest or by contractors (VDFW, USDA-APHIS) |
| Regulations |  |
| • Ban high risk parts from positive states and all scent lures from positive states (VDFW)<br>• No processing or taxidermy of out-of-state deer (VDFW)<br>• Refuge hunting regulations match state regulations (USFWS)<br>• Explore changes to town discharge ordinances (VDFW) | • Refuge hunting regulations match state regulations (FWS) |
| Carcass Management |  |
| • Monitor access areas (FWS, VDFW, VFPR) | • Landowner disposal guidelines (VDFW) |
| Outreach and Education |  |
| • Messaging during hunter education (VDFW)<br>• Guidelines for taxidermist/processor (VDFW)<br>• Messaging at hunt areas (VFPR, USFWS)<br>• Public health website (VDH)<br>• CWD grant proposals focused on outreach and education (USDA-APHIS) | • Messaging during hunter education (VDFW)<br>• Guidelines for taxidermist/processor (VDFW)<br>• Messaging at hunt areas (VFPR, USFWS)<br>• Public health website (VDH)<br>• CWD grant proposals focused on outreach and education (USDA-APHIS) |
| Coordination |  |
| • Interagency participation in working groups (All) | • Interagency participation in working groups (All) |
| Captive Cervid Management |  |
| • Annual inventories and inspections (VAAFM)<br>• Tracking (redundant identification systems and farm tags) (VAAFM)<br>• Testing of mortalities (VAAFM) | • Depopulation if CWD found in facility (VAAFM)<br>• In the event of CWD, facilities are cleaned and land-use restrictions are put in place (VAAFM) |
| • In-state movement restrictions (VAAFM)<br>• Imports are prohibited (VAAFM)<br>• Facilities are double fenced (VAAFM) | • Epidemiological testing (VAAFM)<br>• Cervid industry workshops (VAAFM) |
| **CWD Surveillance** |  |
| • Implement a surveillance program designed to detect CWD prevalence of 2.5% with 90% confidence statewide (VDFW)<br>• Additional samples provided from roadkill and hunter harvest (USDA-APHIS, USFWS) | • Monitor prevalence levels once detected (VDFW) |

### Alternative 5: Moderate intensity proactive and responsive approach

This alternative calls for moderate investments in proactive and reactive resources in the management of chronic wasting disease (CWD) in deer. There is also a focus on expanding the tools available to different agencies. Because reactive measures depend on detecting CWD, there is moderate surveillance implemented. [VDFW, Vermont Department of Fish and Wildlife; VFPR, Vermont Department of Forests, Parks and Recreation; USDA-APHIS, U.S. Department of Agriculture Animal and Plant Health Inspection Services; USDA-FS, U.S. Department of Agriculture Forest Service; VDH, Vermont Department of Health; VAAFM, Vermont Agency of Agriculture, Food and Markets]

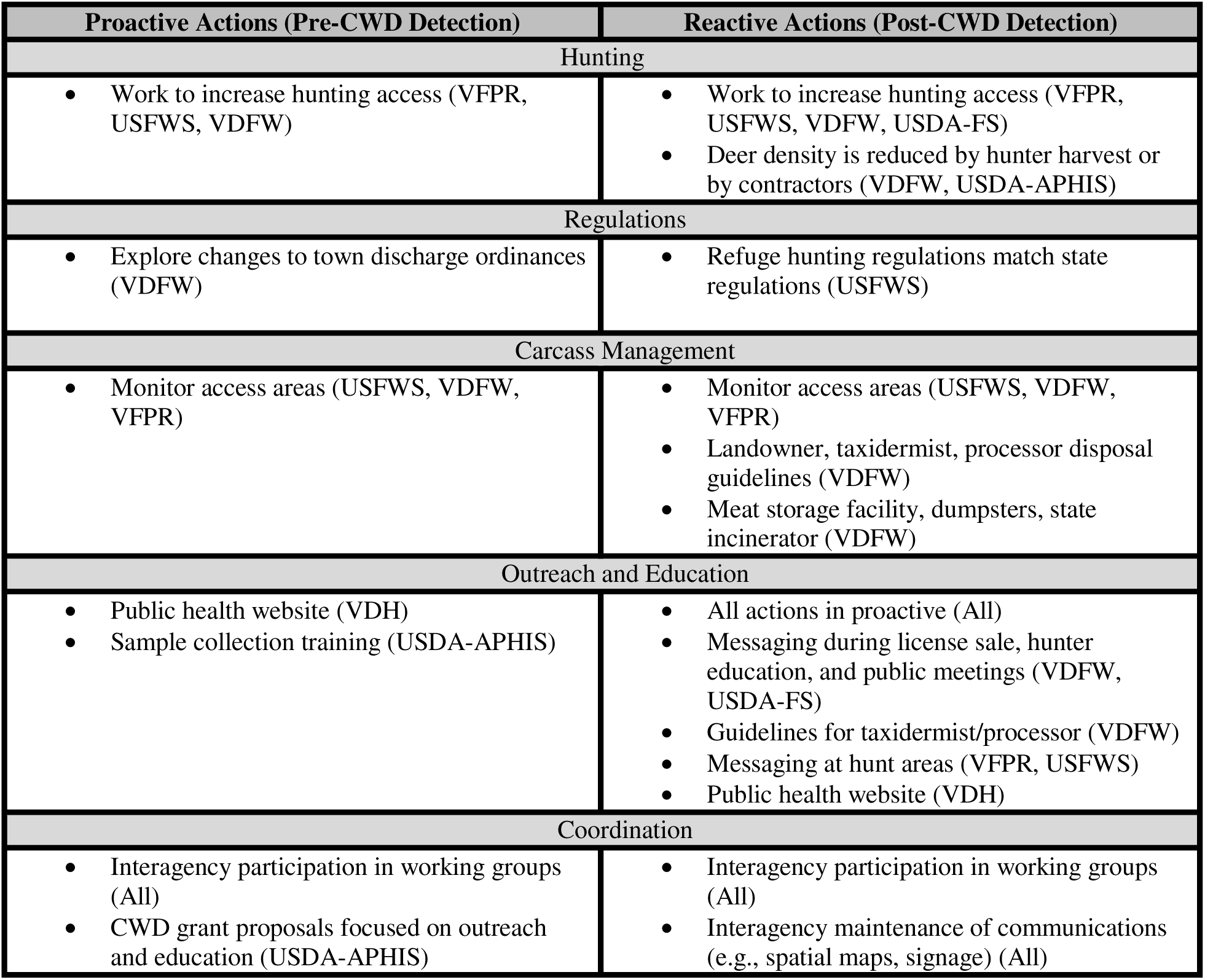
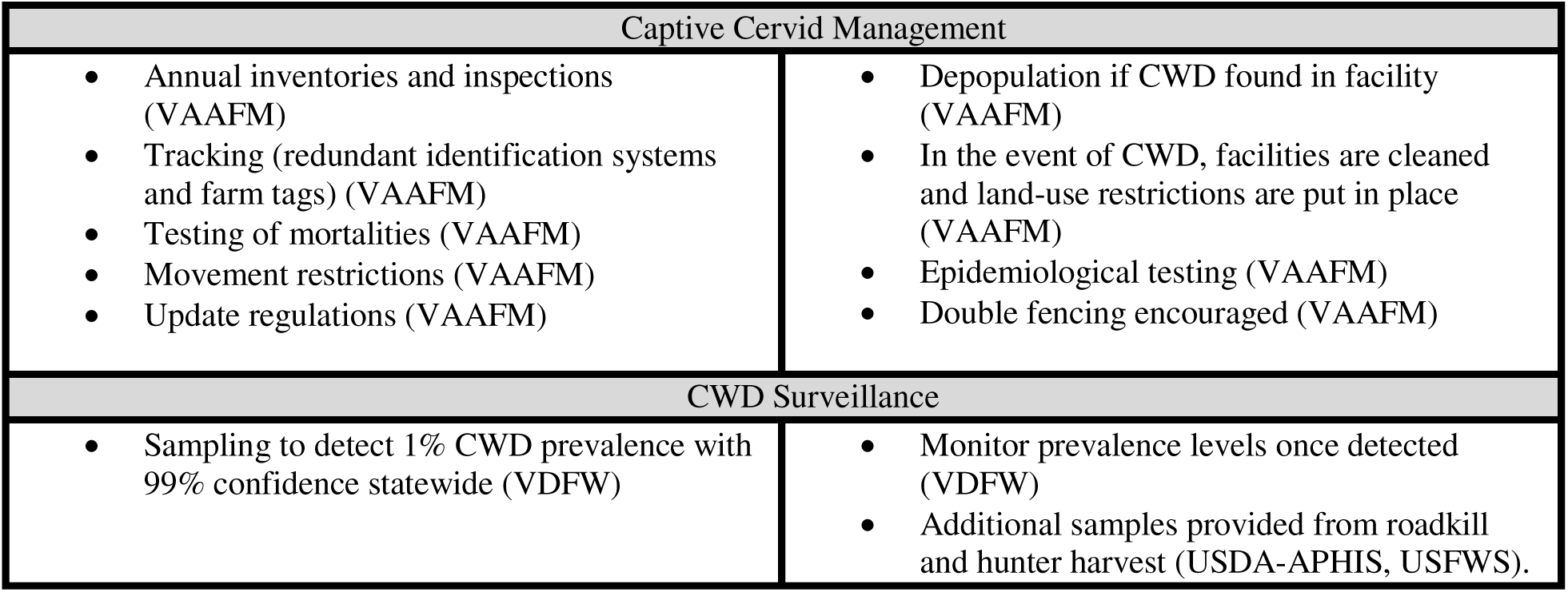

### Alternative 6: Maximum proactive and reactive chronic wasting disease (CWD) response

This alternative calls for all task force agencies to implement the most aggressive actions (both proactive and reactive) that they can. The goal is to explore the efficacy of an alternative that takes all possible actions, and to understand the negative implications of such a strategy. [VDFW, Vermont Department of Fish and Wildlife; VFPR, Vermont Department of Forests, Parks and Recreation; USDA-APHIS, U.S. Department of Agriculture Animal and Plant Health Inspection Services; USDA-FS, U.S. Department of Agriculture Forest Service; VDH, Vermont Department of Health; VAAFM, Vermont Agency of Agriculture, Food and Markets]

| Proactive Actions (Pre-CWD Detection) | Reactive Actions (Post-CWD Detection) |
| --- | --- |
| **Hunting** |  |
| Work to increase hunting access (VFPR, USFWS, VDFW) | Work to increase hunting access (VFPR, USFWS, VDFW, USDA-FS)<br>Deer density is reduced by hunter harvest or by contractors (VDFW, USDA-APHIS) |
| **Regulations** |  |
| Ban high risk parts and scent lures (VDFW)<br>No processing or taxidermy of out-of-state deer (VDFW)<br>Refuge hunting regulations match state regulations (USFWS)<br>Explore changes to town discharge ordinances (VDFW) | Refuge hunting regulations match state regulations (USFWS) |
| **Carcass Management** |  |
| Monitor access areas (USFWS, VDFW, VFPR) | Monitor access areas (USFWS, VDFW, VFPR)<br>Landowner, taxidermist, processor disposal guidelines (VDFW)<br>Meat storage facility, dumpsters, state incinerator (VDFW) |
| **Outreach and Education** |  |
| Messaging during license sale, hunter ed., and public meetings (VDFW, USDA-FS)<br>Guidelines for taxidermist/processor (VDFW)<br>Messaging at hunt areas (VFPR, USFWS)<br>Public health website (VDH)<br>Sample collection training (USDA-APHIS)<br>CWD grant proposals focused on outreach and education (USDA-APHIS) | All actions in proactive (All)<br>Expand public health messaging (VDH)<br>Database of human exposures (VDH)<br>Follow-up outreach to exposed humans (VDH) |
| **Coordination** |  |
| Interagency participation in working groups (All)<br>Establish process for CWD quarantine zones (USFWS) | Interagency participation in working groups (All)<br>Interagency maintenance of communications (All) |
| **Captive Cervid Management** |  |
| Captive cervids banned (VAAFM)<br>Existing farms depopulated or double fenced (VAAFM) |  |
| **CWD Surveillance** |  |
| Sampling to detect 1% CWD prevalence with | Monitor prevalence levels once detected |
| 99% confidence statewide (VDFW)<br>• Additional samples provided from roadkill and hunter harvest (USDA-APHIS, USFWS)<br>• Statewide surveillance database (VDFW) | (VDFW) |

## 15.2 Chronic Wasting Disease Growth Dynamics

**Figure 15.2.1:**
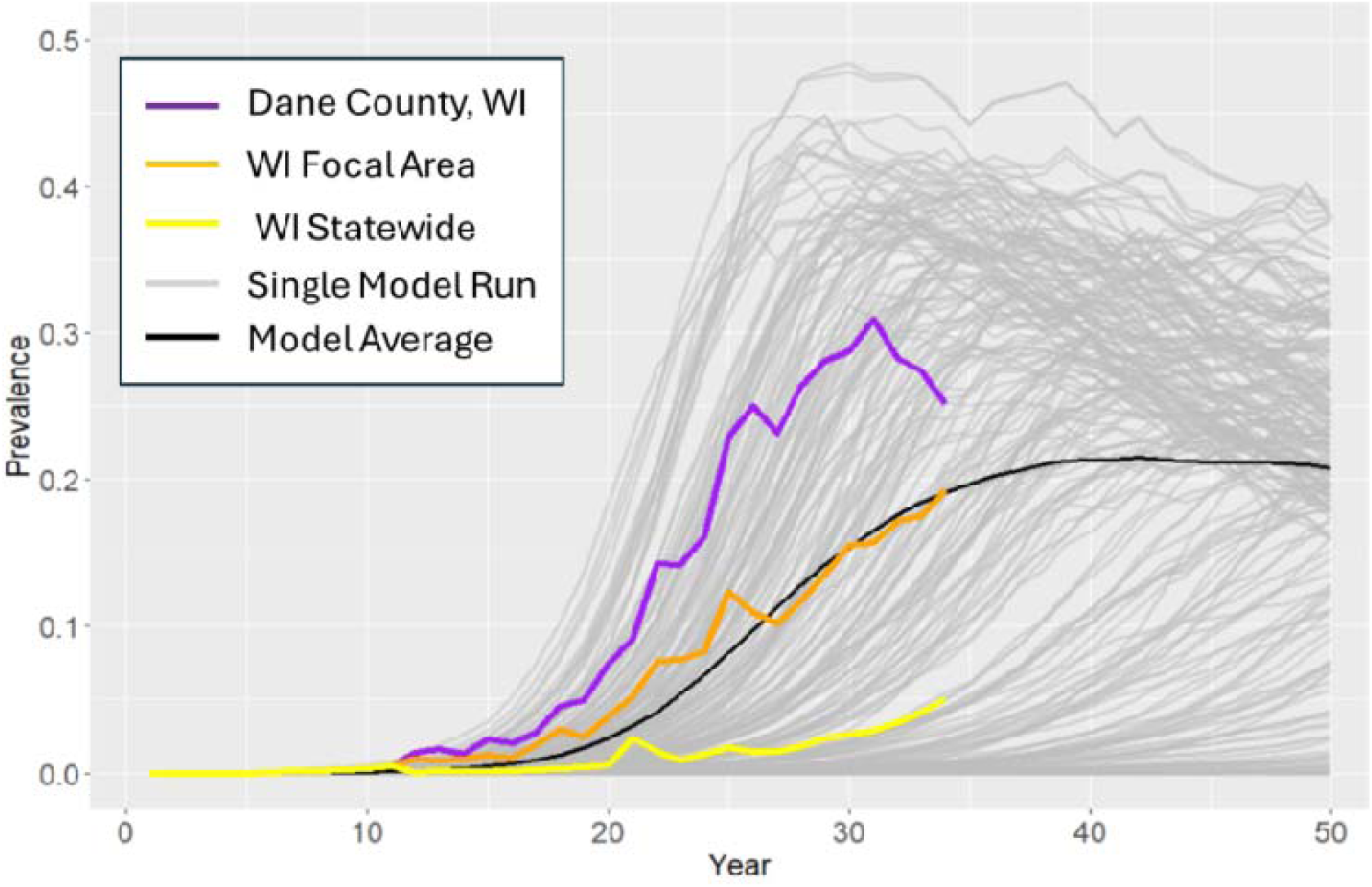
Model performance of chronic wasting disease (CWD) prevalence in deer under status quo conditions with disease introduced in year 1 (20 individuals). Also shown are observational data collected from Wisconsin from 2002-2025 at varying spatial scales. The orange focal area is roughly 2,000 mi^2^ larger than Vermont.

**Figure 15.2.2:**
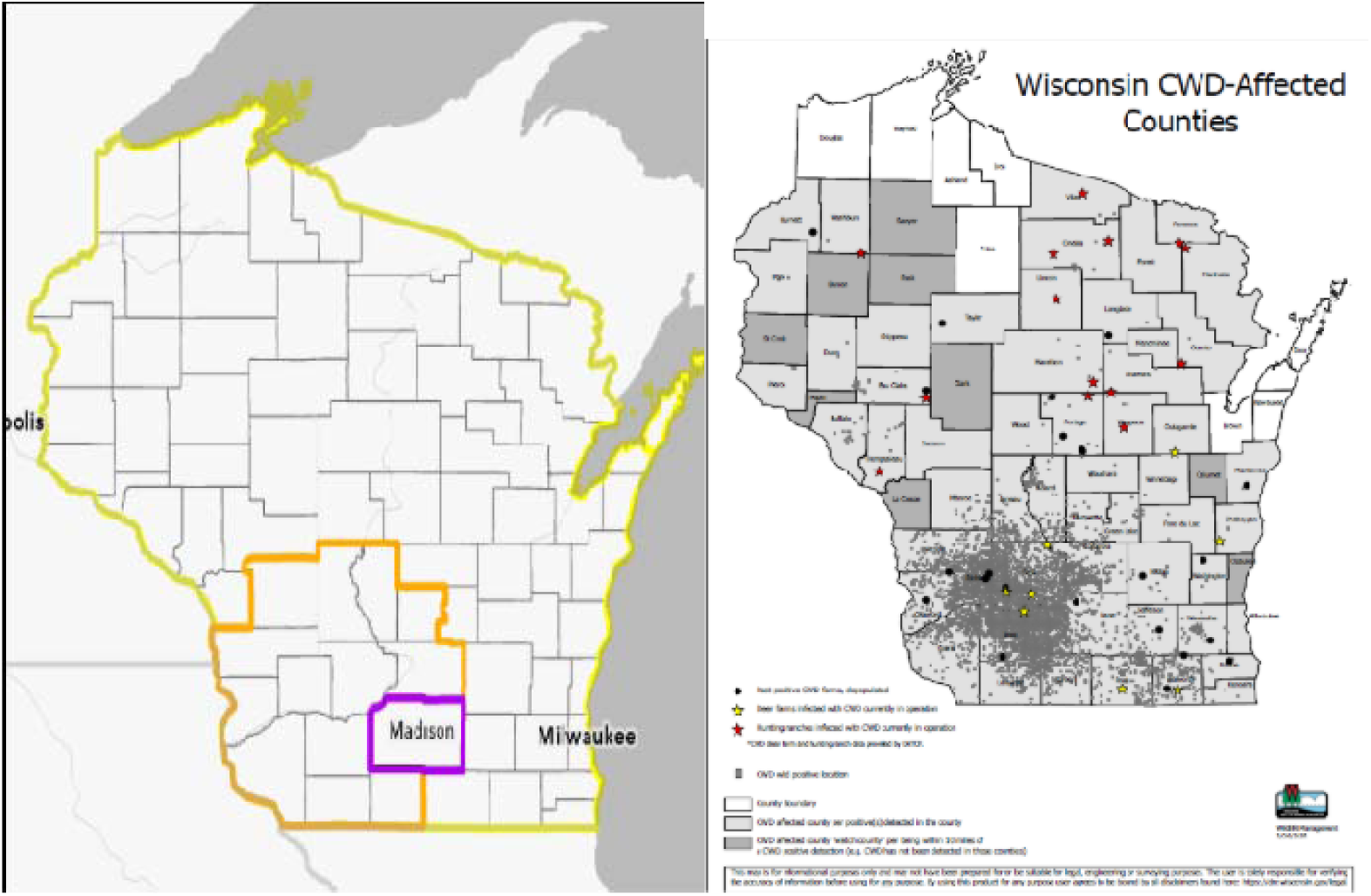
Panel A, Map that corresponds to the three areas of Wisconsin shown in figure 1. Panel B, observational data for chronic wasting disease (CWD) in Wisconsin deer collected 2002−2025.

## 15.3 Derivation of Efficiency of Targeted Removal

In figure 5, Area 1 (dark teal, with radius *r*_1_) is the area of the infected cluster. Area 2 (orange, with radius *r*_2_) is the area of the sharpshooting. We assume the infected cluster has a prevalence of *p*_max_ = 40%, so the number of deer in the cluster is the total number of infected deer (*I* = *pN*) divided by 40%. If the density is *d* = 10/mi^2^, then *r*_1_ = sqrt (*pN*/(*p*_max_*dπ).

We further assume that the sharpshooting area is designed so that a removal rate of *f* = ¾ of the population achieves the desired culling (*S*). Then the required radius is *r*_2_ = sqrt(*S*/(*f*\*d*π)). Given that the center of A_2_ falls somewhere within A_1_, what proportion of A_1_ falls within A_2_? That is, what proportion of the infected cluster is in the sharpshooting radius?

There are two limiting cases:

- Case 1: If the sharpshooting area is entirely within the infected area, then the fraction of Area 1 extracted is the ratio of Area 2 to Area 1.
- Case 2: If the sharpshooting area entirely encompasses the infected area, then the fraction of Area 1 extracted is 1 (100%).

Let

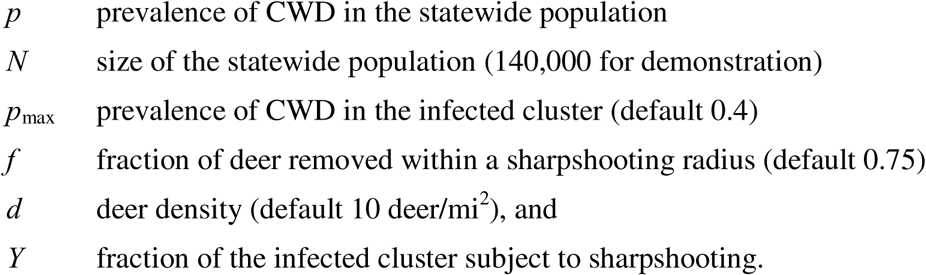

Then, the fraction of the infected cluster (Area 1) that is subject to sharpshooting pressure is

Case 1:

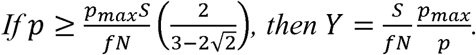

Case 2:

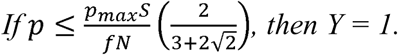

Case 3:

Otherwise, we need to calculate the area of intersection of two overlapping circles. The radius of the infected cluster is given by

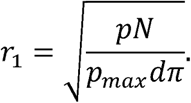

The radius of the sharpshooting effort is given by

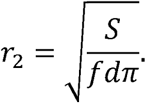

We assume that the center of the sharpshooting radius arises from the detection of a single infected animal, which must be within the infected cluster. If the infected animal is randomly located within the infected cluster (Area 1), then the distance between the centers of the two circles is a random variable with expected value

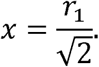

Then, with

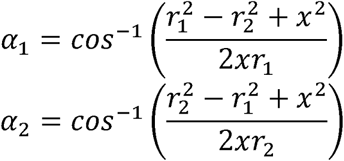

the fraction of the infected cluster that is within the sharpshooting radius is

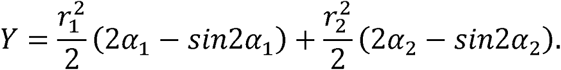

The fraction of infected animals removed is then the product of the fraction of the cluster subjected to sharpshooting (*Y*) and the fraction of deer removed within the sharpshooting radius (*f*).

## 15.4 Proposed Outline for a Proactive Response Plan

**Proactive Response Plan for Chronic Wasting Disease in Vermont**

Outline

Draft, 26 September 2025

This is a proposed, early draft of an outline for Vermont’s proactive response plan for chronic wasting disease in the state. At this stage, this outline is meant to prompt discussion of what elements need to be described in the response plan.

**Background**

**Purpose**

**Participating Agencies**

**Objectives of the Plan**

**Framework of the Plan**

**Coordination**

*Task Force membership*
*Task Force annual meetings*
*Task Force subcommittees*

**Proactive Actions**

*Deer harvest management*
*Captive cervid management*
  Import controls
  Annual herd inspections
  (Pre-emptive closure, double-fencing, etc.)
*Deer carcass management*
  Carcass transportation management (incl. out-of-state import)
  Carcass disposal management
*Public outreach*
*Other preparatory actions*
  Development of CWD-testing protocols and capacity
  Development of partnerships for targeted removals

**Surveillance**

*Wild deer surveillance*
  Pre-detection surveillance design
  Peri-detection surveillance design
  Post-detection surveillance design
*Captive deer surveillance*

**Responsive Actions**

*Establishment of disease management zones*
*Deer harvest management*
  Harvest regulations
  Hunter access to federal and state lands
*Targeted removal of wild deer*
*Captive cervid management*
  Enhanced precautions for uninfected herds in disease management zones
  Quarantine of suspect herds
  Depopulation and decontamination of CWD-positive herds
*Deer carcass management*
  Carcass transport across disease management zones
  Carcass disposal in disease management zones
*Hunter-requested testing of harvested deer*
*Public outreach*
  Hunter outreach
  Cervid farm outreach
  Public health outreach
  General public outreach in areas of targeted removal

**Responsibilities**

**References**

